# Genomic basis for RNA alterations revealed by whole-genome analyses of 27 cancer types

**DOI:** 10.1101/183889

**Authors:** PCAWG Transcriptome Core Group, Claudia Calabrese, Natalie R. Davidson, Nuno A. Fonseca, Yao He, André Kahles, Kjong-Van Lehmann, Fenglin Liu, Yuichi Shiraishi, Cameron M. Soulette, Lara Urban, Deniz Demircioğlu, Liliana Greger, Siliang Li, Dongbing Liu, Marc D. Perry, Linda Xiang, Fan Zhang, Junjun Zhang, Peter Bailey, Serap Erkek, Katherine A. Hoadley, Yong Hou, Helena Kilpinen, Jan O. Korbel, Maximillian G. Marin, Julia Markowski, Tannistha Nandi, Qiang Pan-Hammarström, Chandra Sekhar Pedamallu, Reiner Siebert, Stefan G. Stark, Hong Su, Patrick Tan, Sebastian M. Waszak, Christina Yung, Shida Zhu, PCAWG Transcriptome Working Group, Philip Awadalla, Chad J. Creighton, Matthew Meyerson, B.F. Francis Ouellette, Kui Wu, Huangming Yang, ICGC/TCGA Pan-Cancer Analysis of Whole Genomes Network, Alvis Brazma, Angela N. Brooks, Jonathan Göke, Gunnar Rätsch, Roland F. Schwarz, Oliver Stegle, Zemin Zhang

## Abstract

We present the most comprehensive catalogue of cancer-associated gene alterations through characterization of tumor transcriptomes from 1,188 donors of the Pan-Cancer Analysis of Whole Genomes project. Using matched whole-genome sequencing data, we attributed RNA alterations to germline and somatic DNA alterations, revealing likely genetic mechanisms. We identified 444 associations of gene expression with somatic non-coding single-nucleotide variants. We found 1,872 splicing alterations associated with somatic mutation in intronic regions, including novel exonization events associated with Alu elements. Somatic copy number alterations were the major driver of total gene and allele-specific expression (ASE) variation. Additionally, 82% of gene fusions had structural variant support, including 75 of a novel class called “bridged” fusions, in which a third genomic location bridged two different genes. Globally, we observe transcriptomic alteration signatures that differ between cancer types and have associations with DNA mutational signatures. Given this unique dataset of RNA alterations, we also identified 1,012 genes significantly altered through both DNA *and* RNA mechanisms. Our study represents an extensive catalog of RNA alterations and reveals new insights into the heterogeneous molecular mechanisms of cancer gene alterations.

## Introduction

Organs and tissues are formed by a complex assembly of numerous cell types and their functional heterogeneity is reflected in the diversity of their transcriptional profiles. Despite this, almost all tissue types can give rise to cancer, a process that involves dramatic changes in their transcriptomes. Transcriptional alterations often result from somatic changes in the cancer genome. For example, BCR-ABL1 fusions are caused by single translocation events in chronic myelogenous leukemia^1^ (CML), and HER2 overexpression is frequently the result of focal DNA amplifications^2^. Transcriptomic profiling can be highly informative even in the absence of detectable somatic mutations^3,4^ and even subtle differences in RNA splicing, isoform expression and promoter activation have been associated with cancer^5,6^. Gene expression is predictive of treatment response^7^ and patient survival without a known underlying mutation^8^.

The lack of driver mutations in some tumor samples has been attributed to the bias towards protein coding sequences in the search of recurrent genetic alterations^9,10^. Using whole-genome sequencing, it has been found that recurrent non-coding mutations can act as drivers of cancer, often by altering transcription. However, as non-coding mutations are rare and difficult to interpret, the link between mutations in the non-coding genome, alterations in the transcriptome, and the molecular transformation observed in cancer still remains largely unexplored.

Here, we report the joint analysis of matched transcriptome and genome profiling for 1,188 samples from 27 tumor types from the Pan-Cancer Analysis of Whole Genomes (PCAWG) project^11^, providing the largest resource of RNA phenotypes and their underlying genetic changes in cancer. RNA-Seq data were processed in a standardized way and then analysed to uncover cancer-specific transcriptome changes. We link these changes to somatic variations in the genome, as well as inherited germline background of the patient, highlighting the complexity of transcriptional changes in cancer and the genetic components of these variations. We demonstrate the importance of transcriptomics data in understanding how different dimensions of specific DNA alterations contribute to carcinogenesis, and map out the landscape of cancer-related RNA alterations.

## Results

### Unified data processing for PCAWG RNA sequencing data

For transcriptome analysis, we processed all tumor samples with RNA-Seq data from the PCAWG consortium provided by 30 ICGC projects. To harmonize these data across studies, we reanalyzed a total of 2,217 RNA-Seq libraries using a unified RNA-Seq analysis pipeline developed for this project (software availability described in **Methods**). Core components of this pipeline were spliced alignment of RNA-Seq data followed by gene expression quantification (**Extended Data Figure 1a**). We compared alternative alignment strategies using STAR^12^ and TopHat2^13^ (**Extended Data Figure 1a**), which yielded highly consistent gene expression quantifications (gene-level counts based on HTSeq^14^, **Extended Data Figure 2a**). Thus, we generated consensus gene expression measurements by averaging read counts for each gene, normalized by gene length, followed by upper-quartile normalization (FPKM-UQ)^15,16^ (**Extended Data Figure 2b**). FPKM-UQ quantification across the subset of TCGA samples were highly correlated (median correlation 0.95) with TCGA-reported gene expression using RSEM^17^ (**Extended Data Figure 3**). Transcript isoform-specific expression levels were estimated using Kallisto^18^.

After quality control filtering and merging of technical replicates, we obtained 1,359 RNA-seq profiles from 1,188 unique patients (**Methods**, **Extended Data Figure 4**), with between two and 154 samples per histotype (**Figure 1a**) and approximately equal numbers of male and female patients (**Figure 1b**). For 13 out of the 27 histotypes matched, adjacent normal tissue samples were available, giving rise to a total of 150 normal tissue samples (**Figure 1a,c)**. For additional normal coverage, we processed RNA-Seq data from 3,274 samples from the Genotype-Tissue Expression (GTEx) Consortium (version phs000424.v4.p1) using the same computational pipeline used for the PCAWG RNA-seq dataset. We also generated an adjusted expression dataset using PEER^19^ to account for unknown and technical covariates, as well as a conservative normalization based on quantile normalization. While we observe differences between GTEx and PCAWG samples, likely due to technical differences and batch variation, we find that overall the tissue dominates the expression patterns, suggesting that the combined set provides a useful resource (**Figure 1d, Extended Data Figure 5a**). Tumor purity varied across samples and was considered an additional covariate of expression patterns (**Methods, Extended Data Figure 5b**).

**Figure 1:**
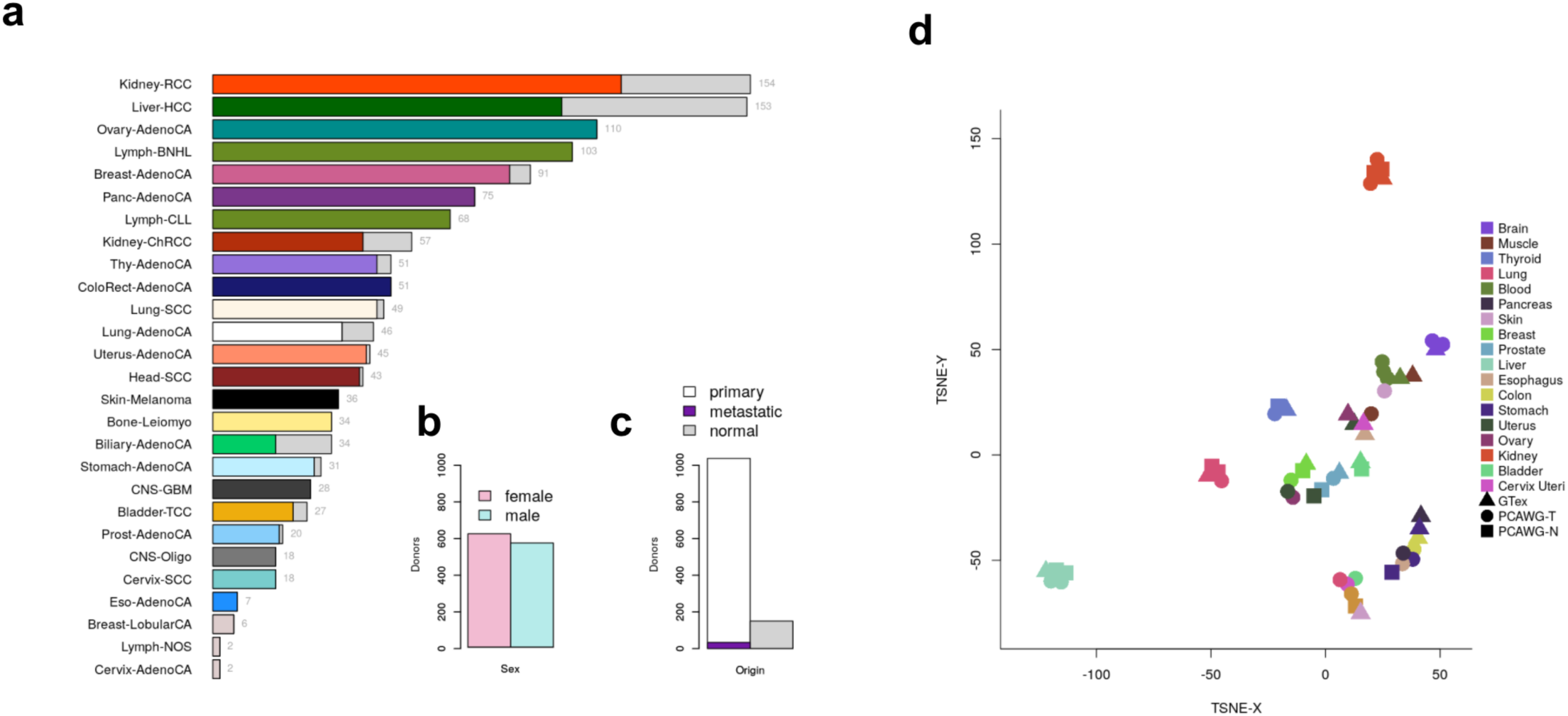
Pan-cancer expression profiling of 1,188 PCAWG donors. **a.** Tumor and normal RNA-Seq data from 27 histotypes. Total number of samples are given to the right of bars. grey bars, normal samples. **b.** Total number of tumor and normal samples from the PCAWG study. A subset of tumors (dark violet) were metastatic. **c.** Number of female versus male donors. **d.** t-SNE analysis of median gene expression aggregated within each project and each GTEx tissue.

### Cancer-specific *cis* germline regulatory variants highlight changes in regulatory landscape

To investigate the underlying mechanisms of different types of RNA alterations, we first focused on changes in mRNA expression level (**Extended Data Figure 6**). To identify heritable Allele Frequency (MAF) ≥ 1%) proximal to individual genes (± determinants of gene expression variability, we considered common germline variants (Minor Allele Frequency (MAF) ≥ 1%) proximal to individual genes (± 100kb around the gene) to map eQTL across the cohort (**Extended Data Figure 7a**) using a linear mixed model to account for population structure and other confounding factors (**Methods, Supplementary Table 1**). This pan-cancer analysis (pan-analysis) identified 3,509 genes with an eQTL (FDR ≤ 5%, hereafter denoted eGenes; **Methods, Supplementary Table 2**), enriched in transcription start site (TSS) proximal regions as expected from eQTL studies in normal tissues^20^ (**Extended Data Figure 7b**). Analogous tissue-specific eQTL analyses in seven cancer types with 60 or more patients identified between 106 (Breast-AdenoCA) and 472 eGenes (Kidney-RCC) (**Extended Data Figure 8a, Supplementary Table 2, Methods**).

To identify regulatory variants that are cancer-specific, we compared our pan-analysis eQTL set to eQTL maps from normal tissues obtained from the GTEx project^21^, adapting a strategy devised previously^22^. For the lead variant, *i.e.*, the most significant SNP of each eQTL, we assessed the marginal replication in GTEx tissues (P ≤ 0.01, Bonferroni-adjusted for 42 somatic tissues excluding cell lines, using SNPS in Linkage Disequilibrium (LD) with r^2^>0.8 for variants not tested in GTEx, **Methods**). 87.5% of eQTL (2,982 of 3,408 accessible eQTL variants, **Methods**) were replicated in at least one GTEx tissue, whereas 426 eQTL did not show any correspondence in GTEx tissues, suggesting cancer-specific regulation (**Figure 2a**, **Supplementary Table 3**). One such example is *SLAMF9*, a member of the CD2 subfamily, with known roles in immune response and cancer^23^ (**Figure 2b, Extended Data Figure 9a**). Similarly, we identified cancer-specific eQTL for genes with known roles in cancer such as *SLX1A*, a regulator of genome stability involved in DNA repair and recombination^24,25^ **(Extended Data Figure 9b**).

**Figure 2:**
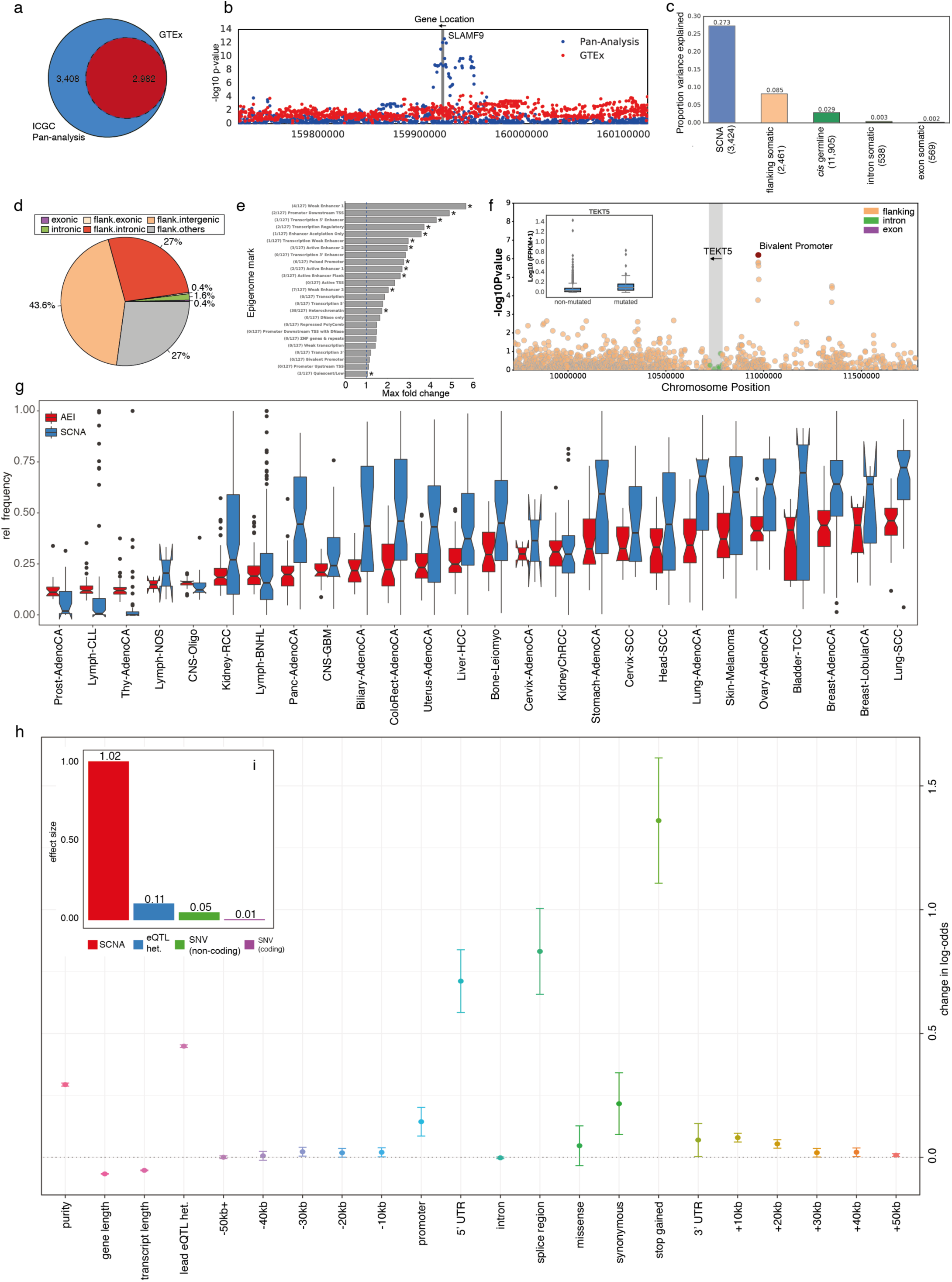
Germline and somatic SNVs associated with expression. The Venn diagram shows the number of eQTL identified in the PCAWGcohort and the fraction of QTL that replicate in the GTEx cohort. 426 eQTL were specific to the PCAWG cohort. Manhattan plot for SLAMF9, showing associations of the pan-analysis in the PCAWG cohort (blue) and in GTEx (red). **c.** Variance component analysis for gene expression levels (Methods). Shown is the average proportion of variance explained by different germline and somatic factors for different sets of genes (grouped based on component that explains the most variance), considering genes for which the largest variance component are i) somatic copy number alteration (SCNA), ii) somatic variants in flanking regions, iii) *cis* germline effects and iv) somatic intron and exon mutations, respectively. The number of genes in each set is indicated in parentheses. **d.** Breakdown of 567 genomic regions that underlie the observed cis somatic eQTL by variant category (Intronic = eGene intron; Exonic = eGene exon; Flank. = 2kb flanking region within 1Mb distance to the eGene start; Flank.intergenic = flanking region in a genomic location without gene annotations; Flank.intronic = flanking region overlapping an intron of a nearby gene; Flank.others = flanking region partially overlapping exonic and intronic annotations of a nearby gene). **e.** Maximum fold enrichment of epigenetic marks from the Roadmap Epigenomics Project across 127 cell lines. The number of cell lines with significant enrichments is indicated in parentheses (FDR ≤ 10%); asterisks denote significant enrichments in at least one cell line. **f.** Manhattan plot showing nominal p-values of association for TEKT5 (highlighted in gray), considering flanking, intronic and exonic intervals. The leading somatic burden is associated with increased TEKT5 expression (P=1.6.10^-06^) and overlaps an upstream bivalent promoter (red box; annotated in 81 Roadmap cell lines, including 8 ESC, 9 ES-derived and 5 iPSC cell lines). The inset boxplot shows a positive association between the mutation status and expression levels. **g.** Distribution of the number of genes with AEI (red) and SCNAs (blue) across the cohort. Cancer types with high chromosomal instability also exhibit highest amounts of AEI. **h.** Relative contribution of different types of somatic mutational burden and other co-variates to the likelihood of observing AEI. Promoter, 5’UTR, splice region and stop gain variants contribute most. **i.** Standardized effect sizes on the presence of AEI, taking only SCNAs, germline eQTLs, coding and non-coding mutations into account. In sum, SCNAs accounted for 86.1% of the total effect size, followed by germline eQTL (9.0%) and somatic SNVs (4.8%).

The majority of these cancer-specific regulatory variants could not be explained by differences in gene expression level between cancer and normal tissues (328/426 genes with at most a 2-fold increase in median gene expression compared to GTEx, *e.g., SLX1A*, **Extended Data Figure 9b**), whereas 98 genes did show evidence of cancer-specific upregulation and ectopic expression (*e.g., SLAMF9*). Among these were immunoglobulin genes and nine cancer/testis antigen encoding genes (CT genes) (**Extended Data Figure 8b**). Cancer testis genes are of interest for their known immunogenic properties^26,27^, and exhibit high expression in sperm and some cancers but are repressed in healthy tissues^28^. We also identified instances of eQTL that replicated in GTEx tissues but not in their corresponding normal tissues. One such example is *TEKT5*, which is expressed in our cohort but otherwise specific to testis in GTEx normal tissues, pointing to upregulation of selected genes in cancer (**Extended Data Figure 8c-e**). This catalogue of common and cancer-specific regulatory sites provides the basis for the following analysis and serves as a resource for analysis of gene regulation in future studies.

### Somatic *cis* eQTL mapping reveals widespread associations with non-coding variants

We explored the effect of *cis* somatic variation on gene expression by aggregating somatic variants in local burdens in genic and non-genic regions (**Methods**). Using these somatic burden elements, *cis* germline variants and SCNAs, we decomposed variation in gene expression of individual genes into these genetic components (**Methods, Figure 2c**). This analysis identified SCNAs as the major driver of expression variation (27.3% on average, **Figure 2c**), followed by flanking somatic and germline variants. Notably, *cis* germline effects, although exhibiting smaller effects on individual genes, explained the largest proportion of variance for 11,905 genes, compared to 3,568 genes, for which somatic factors explained most variation.

We also tested for associations between recurrently mutated intervals (burden frequency ≥ 1%, considering 2kb gene-flanking regions, exons and introns) and gene expression levels (**Extended Data Figure 10a-c**, **Methods**), assessing alternative strategies for burden estimation where burdens weighted by variant clonality maximised detection power (**Extended Data Figure 11a-d**). Genome-wide, this identified 649 somatic eQTL (FDR ≤ 5%; **Supplementary Table 4**) associated with 567 unique regions. Among these, 11 somatic eQTL were explained by mutational burdens in exons or introns, including genes with known roles in the pathogenesis of specific cancers such as *CDK12* in ovarian cancer^29,30^, *PI4KA* in hepatocellular carcinoma^31^, *IRF4* in leukemia^32^, *AICDA* in skin melanoma^33^, *C11orf73* in clear cell renal cancer^34^ and *BCL2* and *SGK1* in lymphoma^35^**, Extended Data Figure 12a-g**). The majority of eGenes (68.4%) involved associations with flanking non-coding intervals (272 intergenic, 172 intronic regions, **Figure 2d)**, and were due to mutations observed in multiple cancers (**Extended Data Figure 13a, Supplementary Table 4**). In contrast to germline variants, these associations tended to be located distal to the TSS (≥20kb, 88%), with larger effects on average than associations proximal to the TSS | *β* | =3.3 versus | *β* | =1.4, **Extended Data Figure 13b**), which points to the relevance of somatic mutations at distal regulatory elements. To assess to which extent structural variants (SVs) may have confounded the gene expression changes observed for the 649 eGenes, we performed a per sample analysis to assess the presence of SVs nearby the leading genomic intervals in each individual with associated mutational burden (**Methods**). This analysis identified 110 (17%) eGenes (**Supplementary Table 4)** with at least one SV close to the individual mutational burden (with a maximum distance observed between the burden and the SV of 40kb). Among the eGenes with SVs, we found immunoglobulin (Ig) genes to be the most prevalent class of eGenes with structural alterations (85/110 eGenes), which is particularly expected for B cell malignancies, where Ig genes are known target of genomic translocations ^36,37^. However, with the exception of Ig genes and few other genes (like *PACS2*), SVs do not seem to frequently co-localize with somatic burden for the 649 somatic eGenes identified.

We next tested the most significant flanking intervals per somatic eGene (lead flanking intervals) for enrichments in cell-type specific regulatory annotations comparing the overlap of true associations to distance-and burden-frequency matched random regions (**Methods**). This identified enrichments for 13 out of 25 epigenetic annotations (FDR ≤ 10%, **Figure 2e**, **Supplementary Table 5**), including poised promoters, weak and active enhancers and heterochromatin in more than two cell lines (**Figure 2e**), but no significant enrichment of TFBS (**Supplementary Tables 6**).

Poised or bivalent promoters are a hallmark of developmental genes and prepare stem cells for somatic differentiation^38^. Re-activation of poised promoters is one mechanism of upregulation of developmental genes in cancer, including CT genes^39^. CT genes were marginally more frequent among genes with somatic eQTL than expected (45/982, P=0.06, Fisher’s exact test), and we observed an enrichment for somatic eQTL in bivalent promoters for CT genes (P=0.04, Fisher’s exact test). Again, we find *TEKT5,* an integral component of sperm, that has been found to be aberrantly expressed in a variety of cancers^40^. We observed a positive association between *TEKT5* expression and somatic mutational burden (prevalently observed in non-Hodgkin lymphoma patients) in a bivalent promoter site close to the 5’ end of the gene (**Figure 2f**). The prevalence of developmental genes among the somatic eGenes was also consistent with a global enrichment (FDR ≤ 10%) for GO categories related to cell differentiation and developmental processes (**Supplementary Table 7**).

### Allele-specific expression captures cancer-specific dysregulation

To facilitate expression analysis on the level of individual haplotypes, we quantified allele-specific expression (ASE), adapting established quality control steps to cancer tissues^41^, where we pooled ASE counts across heterozygous variants within genes to maximize detection power (**Methods**). This allowed us to quantify ASE for between 588 and 7,728 genes per patient (median=4,112 genes with 15 or more ASE reads in 1,120 samples, **Extended Data Figure 14**, **Methods, Table 1**).

**Table 1:** RNA alteration data.

| RNA data | Synapse ID | Total number of gene alterations found | Mean number of gene alterations per donor |
| --- | --- | --- | --- |
| Extended RNA-Seq metadata | syn7416381 | NA | NA |
| Gene expression (PCAWG) | syn3104297 | 577,470 | 486 |
| Gene expression (GTEx) | syn8105922 | NA | NA |
| Transcript-level expression (PCAWG and GTEx) | syn5974793 | NA | NA |
| RNA fusions | syn3107127 | 5,900 | 5 |
| Alternative promoters | syn3354819 | 246,224 | 207 |
| Alternative splicing | syn3107126 | 208,463 | 175 |
| Allele-specific expression | syn5816006 | 545,918 | 460 |
| RNA editing (with a non-synonymous change) | syn7183004 | 167 | <1 |
| Combined gene-centric table (DNA and RNA alterations) | syn7241153 | 1,873,105 | 1,577 |

The table gives Synapse identifiers to RNA analyses reported in this study. To find a particular dataset, the url is: <https://www.synapse.org/#!/Synapse:<Synapse ID>> For example, for gene expression data, the full url is: https://www.synapse.org/#!/Synapse:syn3104297; NA, not applicable

To robustly identify genetic elements that contribute to somatic dysregulation, we considered ASE^42^ to test for allelic expression imbalance (AEI) (FDR ≤ 5%, binomial test, **Methods**). Across the cohort, we observed substantial differences in the fraction of genes with AEI between cancer types (**Figure 2c, Extended Data Figure 14**), and between cancer and the corresponding normal tissue, which appeared to be driven by variation in somatic copy number alteration (SCNA) prevalence^43^ (**Figure 2g, Extended Data Figure 15a,b**).

We used a logistic regression model to identify the determinants of AEI, accounting for the germline eQTL genotype, SCNAs and the weighted mutational burden of proximal somatic SNVs stratified into functional categories (**Extended Data Figure 6**, **Methods**). In aggregate, SCNAs accounted for 86.1% of the total explained effect, confirming our findings from the somatic eQTL analysis, followed by germline eQTL lead variants (9.0%) and somatic SNVs (4.8%) (**Figure 2i**). While cumulatively, non-coding variants were more relevant than coding variants, somatic protein truncating variants (‘stop-gained’) triggering nonsense-mediated decay^44^ were the most predictive individually (**Figure 2h**). This was confirmed by a quantitative model on ASE ratios (**Extended Data Figure 16a-d**). SNVs within splice regions, 5’ UTR and promoters were also strongly associated with AEI presence and we observed a global trend of decreasing relevance of variants with increasing distance from the TSS (**Figure 2h**).

Our model allows for distinguishing AEI caused by germline SNPs, SCNAs and somatic SNVs by computing an average predicted score per gene across the cohort (**Methods, Supplementary Table 8**), which identified somatic AEI as predictive for genes with relevance in cancer (**Extended Data Figure 17**). Motivated by the observed cancer-specific germline regulation of cancer/testis antigen encoding genes (CT genes), we also used these model components to investigate sources of AEI in CT genes. Notably, CT genes were depleted when considering the full somatic score including SCNAs (25/476 CT genes in the top 10% of genes, 48 expected, χ^2^ test, P=6.10^-4^), but enriched in the AEI score based on SNVs only (66/476 CT genes in the top 10% of genes, 48 expected, χ^2^ test, P=6.10^-3^). One potential explanation is that repressed CT genes have to undergo somatic re-activation by SNVs before CN amplification. To elucidate this, we used mutation timing data^45,46^ (**Methods**), stratifying SNVs into the categories *early* and *late* (SNV occurred before and after SCNA at the same locus, respectively) and found strong over-representation of *early* SNVs in 329 out of 7,525 CT gene-patient pairs (216 expected, χ^2^ test, P=4.10^-14^).

### Promoter mutations are rarely associated with a change in alternative promoter activity

In addition to gene expression levels, we also aimed to characterize other RNA expression phenotypes of the cancer transcriptome (**Extended Data Figure 1b)**.

Promoter activity affects gene expression levels and the transcript isoforms expressed^47,48^. To identify active promoters, we combined the expression of isoforms initiated in transcription start sites that are identical or nearby, assuming that these are transcribed from the same promoter (**Methods**). This approach identified 44,639 active promoters (FPKM >0.1 in at least 1% of the patient cohort; **Table 1**). Since promoters were found to be recurrently mutated in cancer^9,10^, we specifically investigated the mutational burden in a 200bp window upstream of annotated promoters in the subset of the PCAWG cohort with matched expression and genome sequencing data (**Figure 3a**). The overall numbers of non-coding mutations in promoters reflect the number of mutations observed genome-wide, with melanoma showing the highest numbers (**Figure 3b**). Only 327 promoters show mutations in more than five samples, the majority of which occurs in Skin-Melanoma and Lymphoma (**Figure 3c**). Promoter mutations in Skin Melanoma are expected due to decreased nucleotide excision repair^49,50^, and frequent mutations in lymphoma can be attributed to activation-induced cytidine deaminase (AID), suggesting that functionally impactful mutations are rare. Indeed, the genes with the highest numbers of promoter mutations are *TERT, CXCR4, PAX5* and *CIITA*, among which only the *TERT* gene that encodes the Telomerase Reverse Transcriptase, a core component of the Telomerase complex, was found to be a driver mutation **(Figure 3d)**^10,51–53^**, indicating that functionally impactful mutations are not very common (Figure 3c,d**). When we calculated the pan-cancer association between promoter mutation burden and promoter activity, we found one significant association involving the possibly non-coding transcript *C12ORF77* with unclear biological relevance (**Extended Data Figure 18**). The major limitation for this analysis is a lack in power: firstly, driver mutations are very rare and only a subset of samples have both expression and mutation data, and secondly there is a reduced sensitivity to detect mutations in promoters compared to other genomic regions due to their GC content^53^. This effect is particularly pronounced for the *TERT* promoter^53^, which nevertheless showed the highest number of non-coding promoter mutations (**Figure 3d**). Promoter mutations have been described for the *TERT* gene^10,51,52^. *TERT* has three annotated promoters, the most frequent mutation occurs at the first promoter, which also includes a UV-related mutation hotspot (**Figure 3e**). While *TERT* does not show a significant association in the pan-analysis, we find that *TERT* promoter mutations are associated with increased promoter activity in individual cancer types^10^ (**Figure 3f**). However, the high number of promoter mutations makes *TERT* an exceptional example compared to all other promoters. Overall, we find that promoter mutation burden is rarely associated with a change in promoter activity and gene expression in the PCAWG cohort.

**Figure 3.**
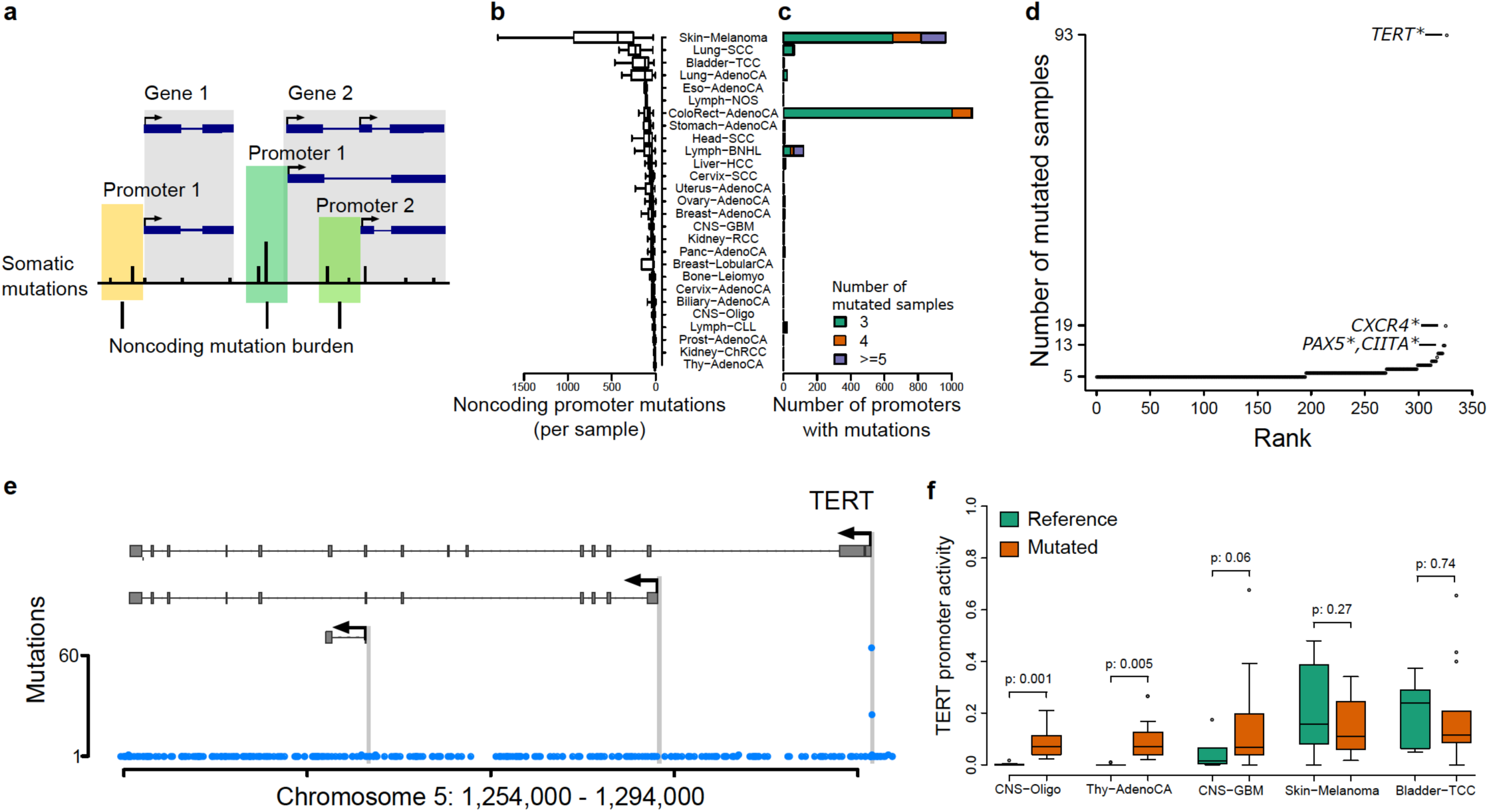
Promoter mutations and their association with expression. **a.** Schematic representation of noncoding promoter mutation burden calculation. **b.** Overview of noncoding promoter mutations per sample. **c.** Overview of the number of mutated promoters per tumor type for promoters with at least 3 mutated samples. **d.** Promoters ranked by the number of mutated samples across all cancer types in a 200bp window, *indicates cancer census genes. **e.** Shown is the TERT locus and the number of mutations observed at each position. The first promoter shows a highly recurrent non-coding mutation reported previously (Bojesen et al., 2013; Rafnar et al., 2009)**. f.** Comparison of promoter activity for mutated and non-mutated samples.

### Intronic mutations associated with splicing and exonization

We identified and quantified alternative splicing using *SplAdder*^54^, focussing on six splicing events types. We found an increase of unannotated alternative splicing events in tumor samples compared to non-tumor samples; for example, there are ≈30% more detected cassette exon events in Liver tumor samples than in matched normals or tissue matched GTEx samples (316,522 tumor, 279,148 normal, 234,710 GTEx; **Extended Data Figure 19a**). In total, *SplAdder* detected 595,041 alternative 3’, 386,734 alternative 5’, 1,226,253 cassette exon, 755,589 intron retention/novel intron, 47,889 coordinated exon skip and 505,515 mutually exclusive exon events in at least one sample of the cohort with Lymph-BNHL, Lymph-CLL, and Ovary-AdenoCA having the most novel events **(Table 1, Extended Data Figure 19b).** While splicing of samples from the same histotype covaries, we observe differences between GTEx and PCAWG cohorts **(Extended Data Figure 19c)**.

Based on our observations of a globally changed splicing landscape, we sought to specifically understand the relation between splicing changes and somatic mutations within introns and to overcome limitations of exome-only studies that are unable to characterize mutations further into the intron beyond the conserved GT/AG donor/acceptor consensus positions.

Focusing on cassette exon events which was the most frequently observed class of alternative splicing, we integrated the quantification of splice events with somatic variants and identified 5,282 mutations near exon-intron boundaries, 1,800 (34%) of which had a large impact (|Z-score| ≥ 3) (**Supplementary Table 9**). Consistent with previous findings using exome-sequencing^55^, a majority of mutations overlapping essential dinucleotides of the 5’ and 3’ splice sites have a strong effect on local splicing, 61% and 57%, respectively **(Figure 4a)**. Relative to a background mutation rate at random intron positions, we found strong associations between splicing outliers and mutations directly at or adjacent to splice sites, with the signal extending much further into the intron. Nearly a third of all mutations 226/469 in a window of 5nt into the intron from the 5’ site were considered impactful. **(Figure 4a, top)**. Almost all impactful mutations had a negative effect on splicing **(Figure 4a bottom, Z-score ≤ −3)** and only very few cases (4% total) enhanced splicing efficiency **(Figure 4a bottom, Z-score ≥ 3)**. Altogether, these results suggest that somatic mutations in the extended splice site region can be as detrimental to splicing as those in the canonical GT-AG dinucleotides.

**Figure 4.**
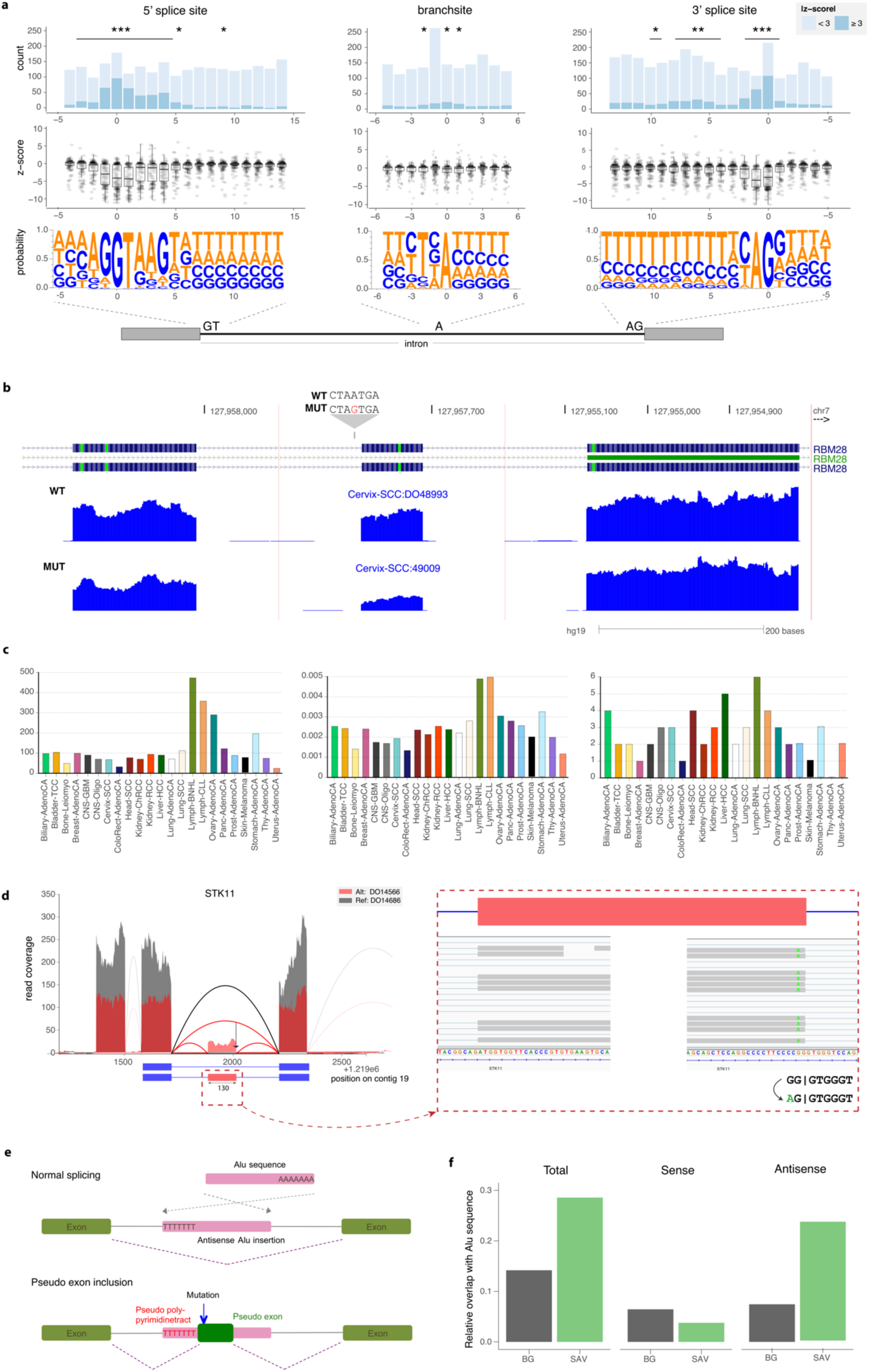
Position specific impact of somatic mutations on alternative-splicing. **a.** Top panel: Proportion of mutations near exon-intron junctions and at branchsites that impact exon skipping events. Impactful mutations are ones in which the percent spliced in (PSI) derived z-score is >= 3 (dark blue). Non-impactful mutations are ones in which the PSI derived z-score is <3. Intron positions that are significantly enriched for impactful mutations are denoted by asterisks (*** < 0.001, ** < 0.01, * < 0.05). Middle panel: Magnitude and direction of mutation-associated splicing alterations. Bottom panel: Sequence motifs of regions. **b.** Example splicing impact of a branch point mutation. UCSC genome browser RNA-seq coverage plots of cassette exon event in RBM28 between mutant and wild-type. Mutant (bottom track) harbors a A->G mutation 29 nucleotides upstream from acceptor site of affected exon. **c.** Distribution of novel cassette exon events only detected within the PCAWG cohort. Top panel shows total number of events per histology type. Middle panel contains the same number, normalized to the total number of cassette exons detected in the histology types. The bottom panel shows the number of exonizations per histotype (novel cassette exons co-located to a somatic alteration near the acceptor or donor of the exon). **d.** Example for exonization event in the tumor suppressor gene STK11. RNA-Seq read coverage for a part of the gene is shown in red for a donor carrying the alternate allele and in grey for a random donor with reference allele. The cassette exon event is shown as schematic below, with blue (red) boxes denoting constitutive (alternative) exons and blue solid lines introns. Zoomed in panels on the bottom show details from IGV visualization, highlighting a somatic mutation at the 3’ end of the cassette exon. The associated sequencing change is illustrated on the lower right corner, where the vertical bar denotes the exon-intron boundary. **e.** Alu-based exonization mechanism. Presence of Alu element in intron in antisense alone will still result in normal splicing (top). Specific mutation of Alu sequence creates novel splice site and creates exonization (bottom). **f.** Enrichment of SINE elements in SAVs compared to sequence background (left). Drawn separately for SINE elements overlapping in sense (middle) and antisense (right) direction.

For mutations in or near the poly-pyrimidine tract, we found a significant enrichment for mutations linked to outlier splicing (**Figure 4a**). Although it is known that *trans* factors that assist in branch site recognition, like *SF3B1*, are recurrently mutated in various cancer types^56^, a pan-analysis of the impact of branch site associated mutations in *cis* has not been performed. Based on recent branchpoint annotations^57,58^, we found 23 impactful mutations at branch site adenosines **(Figure 4a (middle), Figure 4b, Supplementary Table 10)**. Further, we measured positive selection for somatic mutations associated with splicing alterations at a gene-level using a permutation test. Our analysis recovered two known tumor suppressor genes, *TP53* and *FANCA* (FDR ≤ 1%) (**Supplementary Notes**, **Supplementary Table 11**).

Complementing our analyses of global shifts in the splicing landscape, we also studied the contribution of rare splicing associated variants (SAV) that appear in only a small number of samples. We applied the SAVNet approach^59^ which was designed to identify associations between rare somatic variants and local changes in alternative splicing (**Methods**, FDR ≤ 10%, **Extended Data Figure 19d**). SAVs can have various consequences, ranging from the disruption of existing donor and acceptor motifs, to the creation or activation of novel splice sites. In total, we could identify 1,901 SAVs (555/827 acceptor/donor disruptions, 155/364 accept/donor creations) (https://www.synapse.org/#!Synapse:syn3107126). Notably, 1,066 affected canonical splice sites, while the other 806 disrupted non-canonical sites or created novel splice sites. Interestingly, we find a two-fold enrichment of cancer genes in SAVs **(Extended Data Figure 19e)**.

Although we find that splice site creating SAVs (scSAVs) strongly concentrate near exon-intron boundaries **(Extended Data Figure 19f)**, 46.7% of scSAVs are further than 100bp away from the nearest annotated exon. Mutations at those sites generally changed the sequences towards the donor/acceptor motif consensus **(Extended Data Figure 19g)**, providing further evidence for the creation of novel functional splicing motifs. Focusing on novel splice sites deep in introns, we analyzed the extent of exonizations – the formation of novel exons within an intron. To estimate the number of such events within the PCAWG cohort, we filtered all cassette exon events to retain only those that do not occur in the annotation, normal samples, or the GTEx outgroup. Out of 67,254 novel cassette exons, we characterized 3,941 (6%) as exonization events with 45 being located in direct vicinity to somatic alterations **(Figure 4c, Supplementary Table 12)**. Several of those occur in cancer related genes, such as *STK11* **(Figure 4d)**, a well known tumor suppressor kinase acting on AMPK family proteins. As expected, the exonization event would cause a frameshift in *STK11*.

As shown previously, Alu sequences can have a strong impact on exonization^60,61^. They contain elements resembling consensus splice sites, that together with activating mutations, can lead to the formation of consensus splice sites to become a novel exon **(Figure 4e)**. We found a significant enrichment of scSAVs within annotated Alu sequences (P=2.8.10^-9^), particularly for Alus inserted in antisense direction (P=2.6·10^-15^) **(Figure 4f)**. Using pairwise alignments of each Alu sequence overlapping an SAV against the Alu consensus as a reference coordinate system, we found several hotspots of newly created splicing donor and acceptor sites, especially at position 279 close to the poly-T stretch **(Extended Data Figure 18h)**. Our results indicate that Alu sequence exonization, extensively studied in the context of primate genome evolution, is also frequently observed in cancer genome evolution.

### Patterns of gene fusion distribution across cancer indications

Gene fusions are an important class of cancer-driving events with therapeutic and diagnostic values^62^. We identified a total of 925 known and 2,372 novel cancer-specific gene fusions by combining the output of two fusion discovery methods and genomic rearrangement (SVs) information, and several filters were implemented to exclude artefacts or those also present in the GTEx or normal PCAWG samples^63^ (**Methods, Table 1**). Although most gene fusions appear only in one sample, we found that some of the fusion partners are highly recurrent (**Extended Data Figure 20a).**

For the 3,540 identified fusion events representing 3,297 unique gene fusions, we categorized them based on novelty, recurrence, known oncogenic gene partners for downstream analyses **(Figure 5a, Table 1)**. Similar to what was observed in the TCGA tumors cohort previously^64^, the average number of putative gene fusions per sample varies considerably across histological types (mean=3, median=2, sd=3). Most of cancer types (10 of 27) have less than one fusion per sample, while soft tissue-leiomyosarcoma harbour ∼14 fusions per sample on average.

**Figure 5:**
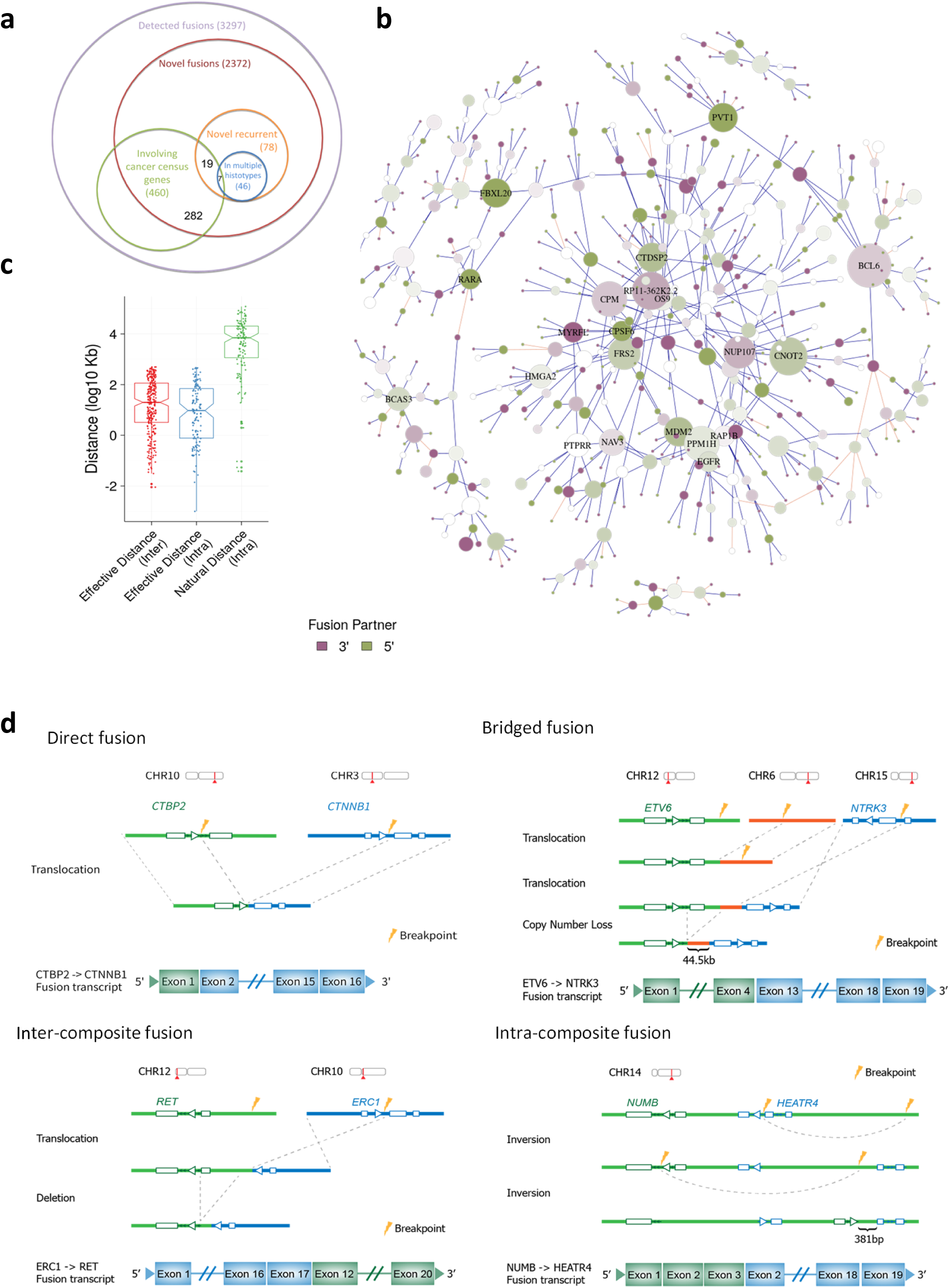
Structural rearrangements associated with RNA fusions. **a.** The number of all detected and novel fusions and their overlap with the cancer census genes. Majority of the fusions are present only in one sample, however, over half of the recurrent fusions are present in several cancer histotypes. From the novel recurrent fusions 19 involve cancer census genes and 7 of them are in multiple histotypes. **b.** Connected clusters of at least 10 genes. Genes are represented as nodes and the size of a node is proportional to the number of gene fusion partners. Two nodes are connected if one fusion was detected involving the two genes: an edge is colored blue if the fusion has matched structural rearrangements evidence and is colored orange otherwise. Nodes and connections are only shown between promiscuous genes. The color intensity indicates if a gene is involved more often in a fusion as 3’ (purple) or 5’ (green) gene or both (white). **c.** Supported rearrangements for composite fusions bring the fused segments of two genes significantly closer. Natural distance indicates the native distance between two related SV breakpoints. Effective distance indicates the distance between the final two breakpoints of the intra-composite/inter-composite fusions. **D.** Schematic representation of examples of different types of SV-supported fusions: *i) direct fusions, ii) bridged fusions, iii)* inter-composite fusions, and *iv)* intra-composite fusions. Bridged fusions are those composite fusions formed by a third genomic segment bridging two different genes. Only one of the possible orders of genomic arrangement is depicted in each case, with breakpoints highlighted as thunderbolts.

Only 71 of the 2373 (∼3%) novel fusions were recurrent, with the majority occurring only in one histotype, while 7 were found across multiple histotypes. Of the 27 most recurrent gene fusions **(Extended Data Figure 20a)**, 8 have been previously reported *(e.g., CCDC6-RET^65^, FGFR3-TACC3^66^, TMPRSS2-ERG, ESR1-CCDC170, PTPRK-RSPO3)* or independently detected in the TCGA cohort^67^*(e.g., GNS-NUP107, TRIO-TERT)*, while 6 were putative novel fusions including *NUMB-HEART4, AFM-FBF1, ESR1-AKAP12*, and *TRAF3IP2-FYN*. In addition to coding region fusions, we also observed 105 fusion transcripts involving the UTR regions of one gene and the complete coding sequences of another. These include a known fusion *TBL1XR1–PIK3CA* in a breast tumour and a notable novel example *CTBP2-CTNNB1* in a gastric tumour, in which the overall transcriptional output of *CTNNB1* is elevated, indicating a potential novel activation mechanism of oncogenes **(Extended Data Figure 20b)**.

Although most involved genes engaged with only one fusion partner, 35 genes had more than five partners. These “promiscuous” genes tended to be selective in being either a 5’ or 3’ partner, and were overrepresented in cancer census genes and in PCAWG’s cancer driver genes (one tailed Fisher’s exact test, odds ratio (OR)=8.66, P≤1.1e-15 and (OR)=12.27, P≤2.2e-16, respectively). Network analysis of promiscuous genes and their partners revealed that most genes belonged to small clusters but several larger clusters emerged. Focusing on clusters with at least 10 genes **(Figure 5b)**, we found that they were significantly enriched in cancer-related pathways (Benjamini-Hochberg corrected P ≤ 0.01) and in protein-protein interactions (P < 1.0e-7). For example, the known oncogene *BCL6* was involved in 15 different fusions, mostly as a 3’ partner with the breakpoints conserved. All such fusions contained the intact exon 2 of *BCL6* and seemed to co-opt the regulatory sequences of their 5’ fusion partners. This pattern had been reported previously in primary gastric high-grade B-cell lymphoma^68^. In general, the breakpoints and their positions (3’ or 5’) were often conserved in promiscuous genes and did not show association with other genomic features such as common fragile sites^69^ **(Extended Data Figure 20c)**, indicating that these genes tend to selectively fuse to other genes. Taken together the data suggests that at least some of the promiscuous fusion partners might play a functional role in cancer progression.

### Bridged fusions and evidence-based gene fusion classification

Our comprehensive data create an unprecedented opportunity to understand their genetic basis of gene fusions. The average number of gene fusions per histological type is highly correlated with the average number of SVs (Pearson correlation 0.95), supporting SVs as a major cause of gene fusions **(Extended Data Figure 20d)**. By examining somatic rearrangement events and fusions simultaneously, we found 2,618 fusion events that could be explained by single genomic rearrangements, with duplication as the predominant type.

Notably, a large number of fusions, including known fusions, namely *ETV6-NTRK3^70^*, could not be associated with any single SV event. The *ETV6-NTRK3* fusion was present in a head and neck thyroid carcinoma sample, linking exon 4 of *ETV6* to exon 12 of *NTRK3*. We found three separate SVs in the same sample: i) a translocation of *ETV6* (chr12:12,099,706) to chromosome 6 (chr6:125,106,892); ii) a translocation of *NTRK3* (chr15:88,694,049) also to chromosome 6 (chr6:125,062,387); and iii) an additional copy number loss (chr12:12,032,501 - chr12:12,099,705) spanning from *ETV6* intron 5 to the exact SV breakpoints (chr12:12,099,706), jointly bringing *ETV6* within 45 kb upstream of *NTRK3*, a distance that would allow transcriptional read-through^71^ or splicing^72^ to yield the *ETV6-NTRK3* fusion^73^ **(Figure 5d)**. Thus, the short chromosome 6 segment appeared to function as a bridge, linking two other genomic locations to facilitate a gene fusion. We term such products *bridged fusions*. This novel class of fusions are not uncommon. Out of a total of 436 fusions supported by two separate SVs, 75 are bridged fusions, with a median length of bridges of 3.7 kb **(Methods, Supplementary Table 13)**.

Aside from bridged fusions, 344 additional fusions are linked to more than one SV in the same sample. These multi-SV fusions are collectively termed *composite fusions*. For example, the known *ERC1-RET* fusion was only supported by an inter-chromosomal translocation and an intra-chromosomal rearrangement, resulting in the connection of *ERC1* to the exon 12 of *RET* **(Figure 5d)**. While fusion transcripts formed by two adjacent genes are often thought to be derived from transcription-induced chimeras, such chimera formation could be facilitated by composite DNA rearrangements. For one of the tumours with the recurrent *NUMB-HEATR4* fusion, we detected two consecutive inversions, bringing the NUMB exon 3 within 381 bp of the *HEATR4* exon 2 **(Figure 5d)**, down from the natural distance of 14 kb, which would might allow for fusion formation by splicing.

Based on the nature of underlying genomic rearrangements, we propose a unified fusion classification system **(Figure 5d, Extended Figure 20e)**. Overall, we identified 75 bridged fusions, 284 inter-composite fusions generated by a translocation linking two genes from different chromosomes followed by a second intra-chromosomal rearrangement, and 125 intra-composite fusions generated by multiple intra-chromosomal rearrangements. Notably, intra-composite fusion partners were brought significantly closer to each other, from the median natural distance of 6,836 kb to the median of 7.9 kb (Wilcoxon Rank Sum Test, P < 2.2e-16, **Figure 5c**). Inter-composite fusion partners also exhibited similarly short gene distances post-translocation **(Figure 5c)**.

While most fusions had direct or composite SV support, for the remaining 18%, including known fusions like *RHOH-BCL6^74^*,we did not detect SV evidence. Thus, either these genes were fused directly at the RNA level or the underlying supporting SVs escaped detection. The latter was evidenced by an observation that known fusions, such as *TMPRSS2-ERG^75^*, did not have consistent SV support in all samples where it was detected (in 4 out of 6 samples this fusion was supported by a deletion, while in the other two samples it did not have any SV support). On the other hand, the 340 SV independent, intra-chromosomal fusions had significantly closer breakpoints than those with SV support **(Extended Data Figure 20f)**. Since read-throughs for such close-by genes have been observed previously^73^, it is likely that certain functional fusions can also be generated by RNA readthrough events.

### Pan-cancer unified analysis reveal diverse modes of RNA-level alterations

Given our comprehensive set of RNA alterations, we sought to characterize the heterogeneous mechanisms of cancer genome and transcriptome alterations. To enable joint analyses of RNA and DNA alterations, we created a gene-level table, indicating the presence or absence of putatively functional events for each gene and patient. Alterations at the nucleotide, amino acid, exon, transcript, or gene level, were all mapped onto the most likely affected gene (**Methods**), and were filtered to exclude types of events that were unlikely to cause functional changes, such as synonymous substitutions or short in-frame insertions or deletions. In particular, we only retained non-synonymous SNVs and RNA editing events as well as splicing events that either induce a frameshift or the alternative region contains an HGMD variant^76^ of the category “damaging” (**Methods**). For quantitative alteration types (expression, splicing, alternative promoters, allele specific expression), the most extreme samples with outlying values within histotype were selected (**Methods, Extended Data Figure 1c**). The resulting binary gene-level table of RNA alterations enables meta-analyses of aberrations across patients, genes, pathways and in specific histotypes. We found no significant correlation between the sample purity and the frequency of outliers (**Extended Data Figure 21).** Over all genes, samples, and alterations we selected 1,871,689 events for further analysis, which was 1.17% of all alteration events. It should be noted that we chose to only include RNA alterations with potential functional effects and with strongest quantitative impact, resembling similar strategies for filtering DNA alterations (**Methods**)^77^. The exact number of alterations for each of the alteration types does depend on filter parameters and those were chosen to have a low observed alteration frequency across the samples. A summary of all identified RNA alteration events is given in **Table 1** and **Supplementary Table 14**.

Building on the gene-centric table, we characterized gene alterations at the RNA-level and contrasted these with DNA alterations (non-synonymous SNVs, SCNAs) identified through whole-genome sequencing analysis from the PCAWG consortium ^78^. To check the quality of the gene-level tables, we tested whether each of the alteration types exhibits cancer specificity. We performed gene set enrichment analysis for top genes ranked by their recurrence within each alteration type against the union of COSMIC cancer census genes^79^ and driver genes identified in the PCAWG cohort^80^. We found that all eight alteration types, six RNA and two DNA alterations, had a significant enrichment for cancer census genes as well as PCAWG driver genes (FDR ≤ 5%, hypergeometric test).

When comparing gene alteration frequencies across all histotypes (**Figure 6a, Extended Data Figure 22**), we note that different cancer types harbor distinct combinations of DNA-and RNA-level alterations (**Figure 6a)**. While, as expected, skin melanoma significantly exceed other cancers in the number of non-synonymous SNVs^81^ (Wilcoxon Rank Sum Test, P < 0.012), lymphatic cancers have low numbers of SNVs (Wilcoxon Rank Sum Test, P = 5.3. 10^-15^) but high incidences of alternative splicing (Wilcoxon Rank Sum Test, P < 2.2.10^-^6^^). While the overall numbers of gene fusions are dwarfed by other types of alterations across cancer types, breast & ovarian adenocarcinomas and soft tissue-leiomyosarcoma are more profoundly impacted by gene fusions (Wilcoxon Rank Sum Test, P < 1.2·10^-6^). Although oligodendrogliomas have relatively low number of SCNAs and non-synonymous SNVs (Wilcoxon Rank Sum Test, P = 0.0061, 1.1.10^-6^), their RNA-level alterations are more comparable with other tumor types, suggesting that transcriptomic alterations can be more impactful in certain cancer indications.

**Figure 6:**
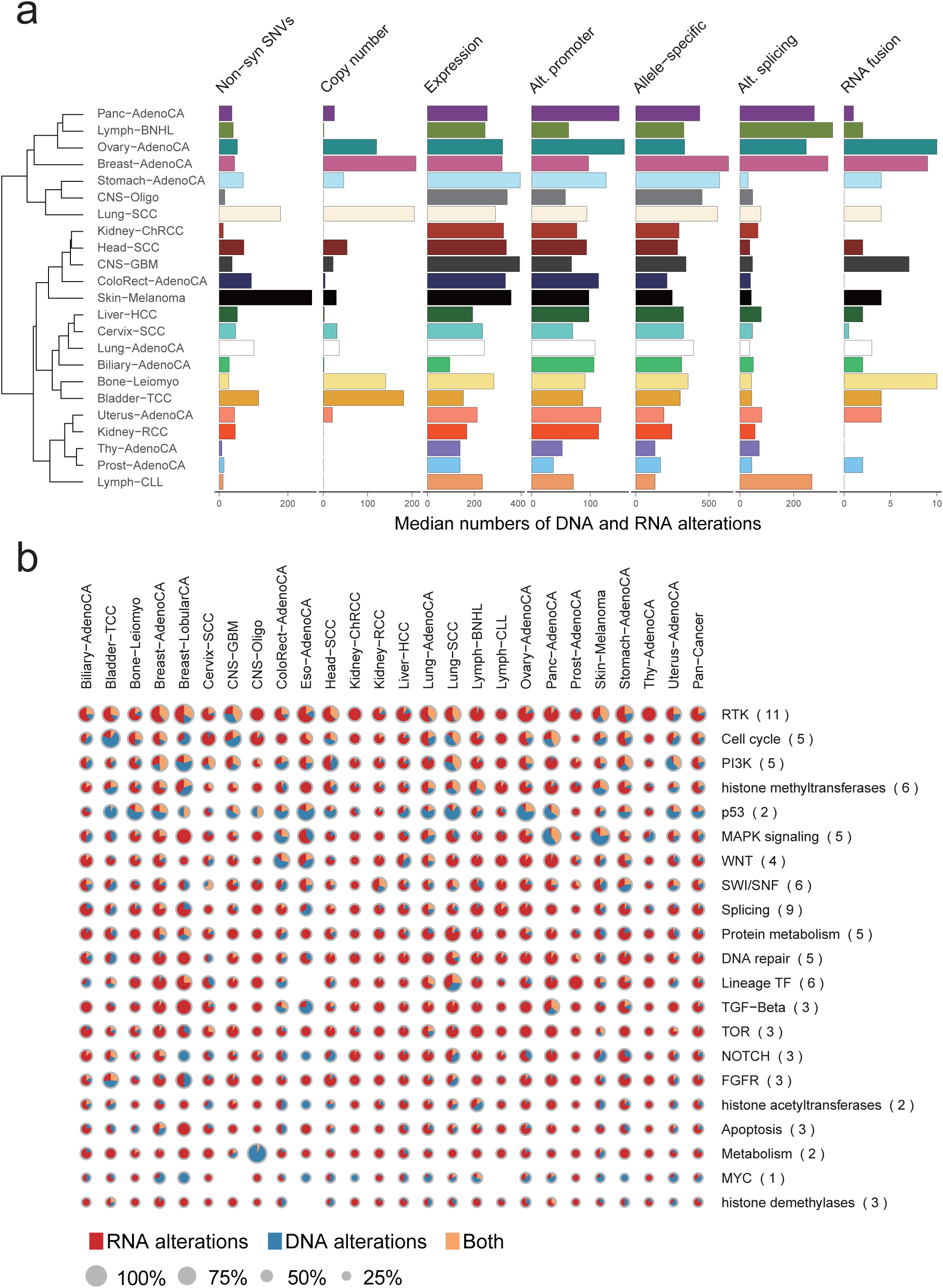
Global view of DNA and RNA alterations affecting tumors. **a.** Barplots showing the median numbers of different alterations across histotypes. Histotypes are ordered by hierarchical clustering based on the pattern of different types of alterations. Only histotypes with more than 10 patients are shown. **b**. Composite pie-charts showing the percentages of RNA alterations, DNA alterations or both, affecting sets of genes in well-characterized cancer pathways and known to be functionally altered in cancer. The sizes of circles represent the percentages of patients affected based on the given gene set. The columns indicate different cancer types.

To evaluate the extent that RNA changes provide additional mechanisms for cancer gene alterations, we examined DNA and RNA-level alterations both in individual genes (**Extended Data Figure 23a**) and in sets of genes in pathways with known roles in cancer^82,83^ (**Figure 6b**). We found that RNA alterations occur at a high proportion in many pathways, including the TOR and metabolism pathways. Even for genes or pathways typically associated with high non-synonymous alterations, such as the p53 pathway, there are also a sizable proportion of RNA alterations. Among the 576 samples altered in the p53 pathway, 131 (22.7%) of them carried only RNA alterations, indicating that neglecting transcriptomes would underestimate the degree of cancer pathway alterations. *MDM2*, in particular, is more frequently altered via RNA alterations than DNA alterations, while *TP53* is altered primarily through non-synonymous SNVs (**Extended Data Figure 23b**) although 13% of all TP53-impacted tumors exhibited changes at both DNA and RNA levels (**Extended Data Figure 23c**). In addition, while *IDH1* is predominantly altered at the DNA level in oligodendroglioma, its alterations in stomach adenocarcinoma are almost exclusively at the RNA level (**Extended Data Figure 23a**), indicating diverse modes of gene alterations that vary by histotypes.

### Co-occurrence of RNA and DNA alterations

The diverse types of alterations in this study enabled us to investigate trans-associations between different genetic and expression characteristics. Indeed, known genetic associations, such as the co-occurring mutations of *KRAS* and *PIK3CA^84^*, and those between *LATS2* and *NF2^85^*, could be recapitulated in this study (**Extended Data Figure 24a**). We then performed systematic co-occurrence analysis to identify all significant *trans*-associations involving cancer-related genes (FDR ≤ 5%, see **Methods**). By investigating how somatic mutations of known cancer genes may impact the expression of other genes, we found *MYC* and *NFKBIE* to be widely linked to dysregulation of many genes (**Figure 7a**), consistent with their known transcriptional regulatory roles in cancer^86–89^. Among other top-ranking genes, *CCND3* mutations co-occurred with *MYC* mutations (P = 6.10^-14^), in agreement with their reported joint function in cancer^90^. Notably, *B2M* mutations are associated with multiple expression changes (**Figure 7a**, **Extended Data Figure 24b**). Pathway enrichment analysis of the top 100 genes associated with all *B2M* alterations, including gene fusions and expression outliers, indicates that the most impacted genes are involved in immune systems and DNA repair (FDR ≤ 1%) (**Figure 7b, Extended Data Figure 24c**). *B2M* encodes an MHC I heavy chain protein involved in antigen presentation and has been previously linked to immune escape^91^. In Lymph-BNHL, the histotype with the most *B2M* altered tumors, donors with altered *B2M* tend to carry more non-synonymous mutations (Wilcoxon Rank Sum Test, P = 0.0097) (**Extended Data Figure 24d**). Therefore, we hypothesize that tumours with *B2M* alterations may better tolerate DNA repair deficiencies.

**Figure 7:**
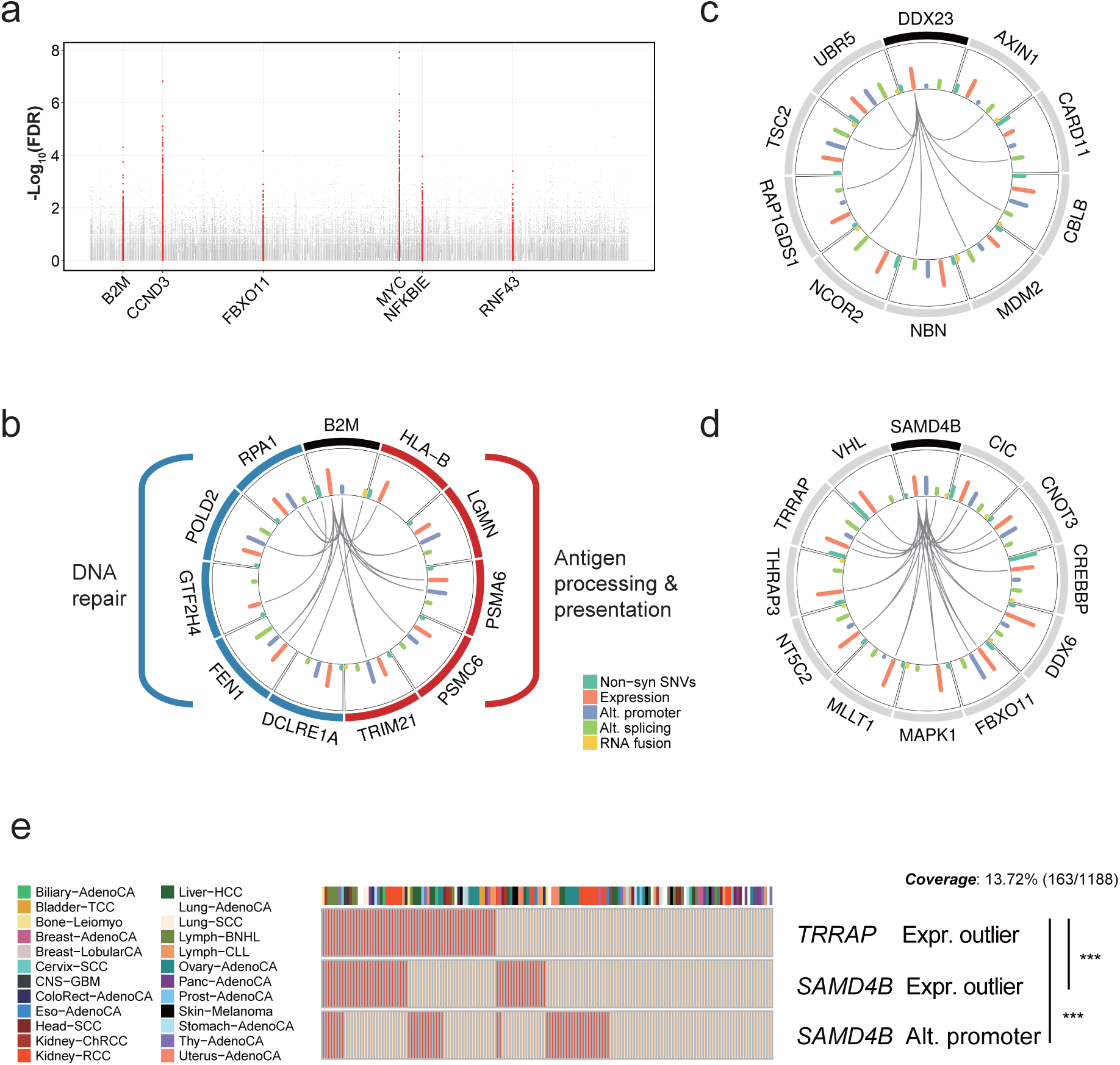
Co-occurrence analysis reveals trans-associations. **a**. Manhattan plot for association of gene expression outliers with cancer gene variants. Each dot represents an alteration pair. The x axis shows all COSMIC genes ordered alphabetically and the y axis represents the FDR-adjusted p values (q values) based on Fisher’s exact tests. COSMIC genes with more than 10 significant associations (FDR ≤< 5%) are colored in red and their names labeled. **b.c.d**. Circos diagrams show the selected genes significantly co-occurred with B2M (b), DDX23 (c) and SAMD4B (d). Connecting lines indicate the specific types of co-occurrences of alteration pairs. The inner histograms indicate the frequencies of incidences of different alteration types shown in different colors. **e**. Heatmap showing the co-occurrence between the TRRAP expression outliers and the SAMD4B transcriptional alterations. Each column indicates a specific tumor with tumor types annotated to the left. *** Adjusted P < 0.001 indicates the significance of the association for the given alteration pairs. Most samples without the listed alterations are not shown for space considerations.

Conversely, we also examined how cancer related genes could be regulated by others through detected co-occurrences. By focusing on genes involved in splicing, we found *PRPF6*, previously reported to preferentially alter splicing of growth regulation genes and thus drive cancer proliferation^92^, to be linked to 16 alternative splicing events of cancer genes (FDR ≤ 5%) (**Extended Data Figure 24e**). Notably, expression outliers of *DDX23* co-occurred with aberrant splicing of a large number of cancer-related genes, including *MDM2* and *TSC2* (**Figure 7c**). *DDX23* encodes a component of the U5 snRNP complex and is involved in nuclear pre-mRNA splicing, suggesting a possible role of *DDX23* in regulating the splicing of cancer-related genes. Similarly, multiple types of alteration of *SAMD4B* are associated with the expression outlier of a panel of cancer-related genes such as *MAPK1* and *VHL* (**Figure 7d**). In particular, *SAMD4B* and the tumor-associated *TRRAP* gene showed co-occurrence for multiple types of alteration (**Figure 7e**). *SAMD4B* is a gene with limited functional information but has been reported to inhibit TP53 and AP-1 activities^93^, so it is reasonable to speculate that *SAMD4B* regulates the expression of key cancer genes. Overall, the *trans*-associations we uncovered add novel insight into the regulatory network in cancer.

### Associations between somatic mutational signatures and gene expression

Global variations in mutational patterns can be quantified using mutational signatures, which tag mutational processes specific to their tissue-of-origin and environmental exposures^94^. However, a pan-cancer analysis of the relationship between genome-wide mutational signatures and gene expression levels has not been explored yet.

We considered 28 mutational signatures derived using non-negative matrix factorization of context-specific mutation frequencies in the PCAWG cohort (PCAWG-7 beta 2 release^95^). We tested for association between signature prevalence in patients and total gene expression, accounting for total mutational burden and other technical and biological confounders (**Methods**). This identified 1,176 genes associated with at least one signature (FDR ≤ 10%, **Extended Data Figure 25**, **Supplementary Table 15**), a markedly different set of genes compared to associations with total mutational burden alone (**Supplementary Table 15**). Lymphoma Signature 9 showed the largest number of associations, followed by the smoking - related Signature 4 (**Figure 8a, Supplementary Table 15**).

**Figure 8:**
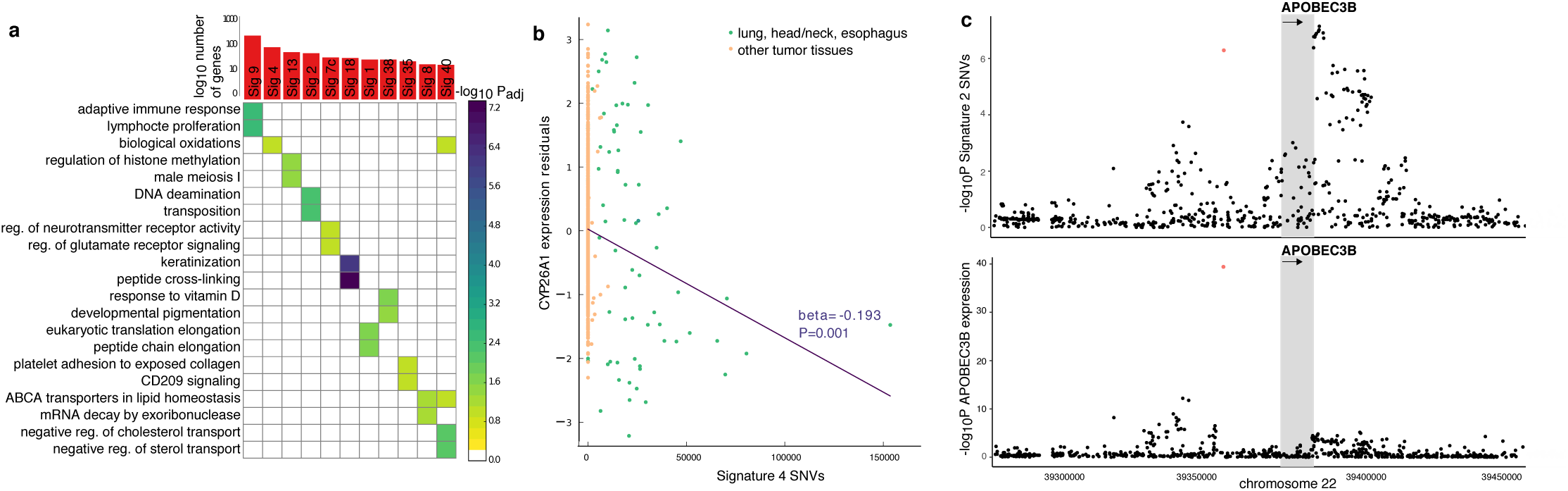
Associations between mutational signatures and gene expression. **a.** Summary of significant associations. Top panel: Total number of associated genes per signature (FDR ≤ 10%). Bottom panel: Enriched GO categories or Reactome pathways for genes associated with each signature (FDR ≤ 10%, significance level encoded in color, −log_10_ P_adj_). **b.** Representative signature-gene association, depicting a negative association between *CYP26A1* expression and Signature 4. **c.** Manhattan plots of associations between *cis* germline variants proximal to *APOBEC3B* (plus or minus 100kb from the gene boundaries) and Signature 2 (top panel) or *APOBEC3B* gene expression level (bottom panel). The gray region denotes the gene body, the orange variant the lead eQTL variant rs12628403.

While some signatures have clear aetiologies, others are not fully characterized. To annotate these signatures *de novo*, we considered 18 signatures with 20 or more associated genes (**Methods, Extended Data Figure 26**) and assessed enrichment using GO categories^96,97^ and Reactome Pathways^96,97^. We found that 11 signatures were enriched for at least one category (FDR ≤ 10%, **Supplementary Table 15**), revealing associations consistent with known aetiologies (**Figure 8a**). For example, Lymphoma Signature 9 was associated with 354 genes enriched for lymphocyte/leukocyte-related processes and immune response, including *TCL1A, LMO2* and *TERT* (P=1.2. ^-10^ .10^-10^, 2.0.10^-09^). The smoking Signature 4 was associated with 119 genes enriched for biological oxidation processes (*e.g.,* benzo[a]pyrene) and including *CYP24A1*, a gene that is known to be down-regulated in tobacco-smoke exposed tissue^98^ (**Figure 8b**). The 70 genes associated with APOBEC Signature 2 were significantly enriched for DNA deaminase pathways.

Among signatures with unknown aetiology, our results link Signature 8, prevalent in medulloblastoma, to 25 genes enriched for ABCA-transporter pathways. Drugs targeting these pathways are currently in clinical trials for treating medulloblastoma^97,99^. Signature 38, which is correlated with the canonical UV Signatures 7 (*e.g.*, 7a: r^2^=0.375, P=5·10^-40^, **Extended Data Figure 26c**), was linked to melanin processes (**Figure 8a**). Melanin synthesis causes oxidative stress to melanocytes^100,101^ and we found Signature 38 associated with the oxidative stress promoting gene *TYR*^*102*^ (P=1.0·10^-4^). A hallmark of Signature 38 are C>A mutations, also a typical product of reactive oxygen species mediated by activity of 8-hydroxy-2′-deoxyguanosine (DFG, 2010). This suggests that Signature 38 may capture DNA damage indirectly caused by UV after direct sun exposure due to oxidative damage^103^, with *TYR* as a possible mediator of the effect.

The cause-and-effect relationship of correlated somatic variations and gene expression changes are not clear *a priori*. We utilized germline eQTL lead variants of signature-associated genes as an anchor to gain directed mechanistic insight by testing for associations between these variants and the signature. This eQTL-based approach entails substantially fewer tests than genome-wide analyses^104,105^. Among 1,176 signature-linked genes, 197 had a germline eQTL, but we found only *APOBEC3A/B* eQTL rs12628403 to be associated with the corresponding Signature 2 (P=5.1·10^-7^, **Figure 8c**, FDR ≤ 10%, multiple testing over 197 tests, **Supplementary Table 15**), confirming it as a risk variant for Signature 2 prevalence^106^. Colocalisation^107^ and mediation^108,109^ analyses confirmed the variant as a plausible genetic determinant of *APOBEC3A/B* expression and Signature 2 prevalence (**Supplementary Table 15**), with a remarkable 87.11% of the genetic effect conferred to the signature by *APOBEC3B* expression (**Extended Data Figure 27**, **Methods**).

In summary, we identified global *trans* effects between mutational signatures and gene expression levels and thereby derived *de novo* annotations of signatures with previously unknown roles.

### Cancer genes are altered in *cis* through heterogeneous mechanisms

In our analyses of *cis*-acting mutations associated with these individual RNA phenotypes, the vast majority were observed rarely in the PCAWG cohort. Many cancer genes (*e.g., MET^4,110,111^*) are known to be somatically altered through heterogeneous mechanisms such as gene fusions, splicing mutations, and nonsynonymous mutations; therefore, looking at genes that are altered through multiple *cis*-acting mechanisms may help to identify novel cancer genes in which an individual alteration type is rare. A total of 5,413 genes were altered through gene expression, allele-specific expression, splicing, and/or gene fusion and had an associated DNA-level mutation in *cis* (**Methods**, **Supplementary Table 16**). PCAWG-defined driver genes tended to have more diverse mechanisms of RNA-level alterations when compared to genes that have not been previously identified as a cancer gene (P < 0.001) (**Figure 9a**). We identified, for instance, a somatic eQTL, a splicing associated variant, and fusions in the known tumor suppressor *NF1* in the MAPK pathway (**Figure 9b**). Another gene with a somatic eQTL, a splicing associated variant, and a fusion event was *PTGFRN*, a gene currently not in the COSMIC cancer gene census (**Figure 9c**). Interestingly, both the fusion event and splicing event preserve the frame of the resulting gene products. Further investigation is necessary to understand the functional impact of these RNA alterations.

**Figure 9:**
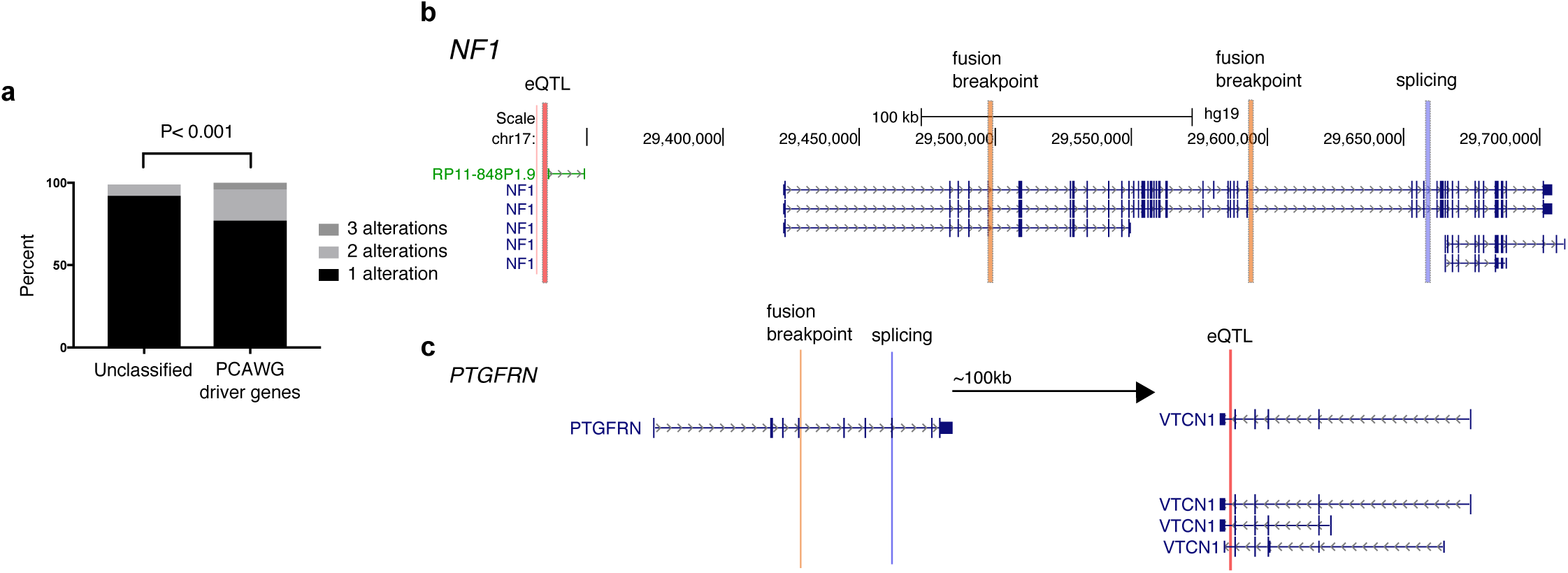
Genes can be altered in cis through multiple mechanisms. **a**. Genes with at least one type of RNA alteration that also has an associated change at the DNA-level in *cis*. Genes are either classified as a PCAWG driver gene or not classified as a driver gene nor a cancer gene from the cancer gene census. Examples of a known cancer gene, **b.** *NF1,* and an unclassified gene, **c.** *PTGFRN,* having heterogeneous mechanisms of alterations.

### Known and novel candidate driver genes are recurrently altered at the RNA-level

Previous pan-cancer analyses have shown that most somatic mutations occur in a small number of genes, and that these alterations are rare^83,112,113^. Due to lack of recurrence of observed alterations, it has been difficult to statistically distinguish functionally relevant, potential driver alterations from passenger alterations. To take advantage of the large variety of DNA and RNA alterations available in this study and motivated by the great diversity of alterations in known driver genes (**Figure 9**), we aimed to identify genes that are both recurrently and heterogeneously altered, under the hypothesis that these genes have increased functional relevance.

This analysis identified 1,012 genes with significant recurrent aberrations (FDR < 5%, **Figure 10a,** permutation-based significance estimation, **Methods**), with the top ranking genes carrying both RNA and DNA aberrations, where RNA alterations account for 13.0-99.4% (mean: 81.4%) of all identified alterations in each gene (**Figure 10a, 10b, Supplementary Table 17**). This ranking is enriched for the union of cancer census genes^79^ (101/603) and PCAWG-defined driver genes (51/157, P=5·10^-26^, enrichment 2.82, **Figure 10c**). *TP53* has nearly the highest proportions of DNA alterations (71.8% DNA alterations, 413/575) and *IGF2* has the highest proportion of RNA alterations (98.5% RNA alterations, 263/267). Furthermore, when we specifically look at the two most frequent alterations for each gene, a majority (75.1%) of the alterations are at the RNA-level (**Figure 10d**). While the total number of RNA alterations does depend on the selected filter parameters, it appears reasonable to conclude that RNA alterations are more likely to occur than DNA alterations for most genes.

**Figure 10:**
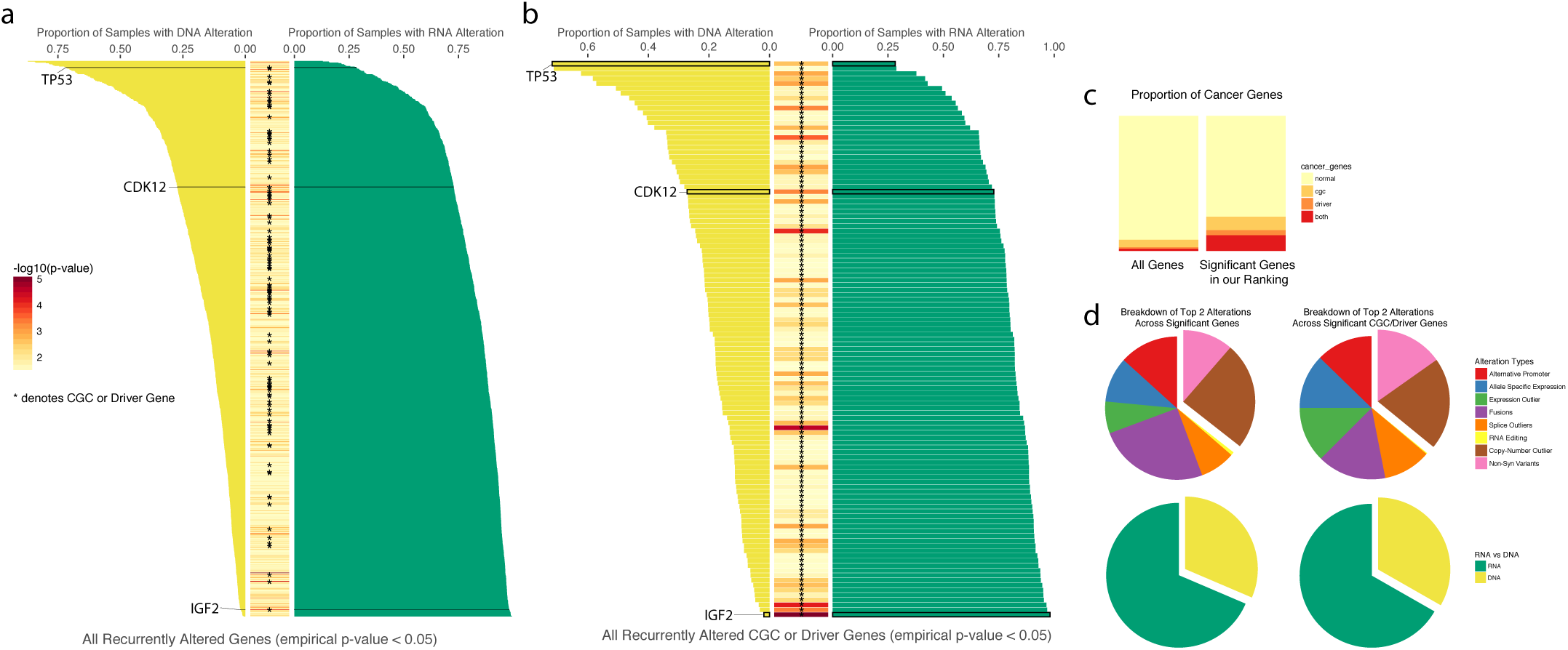
Significantly mutated genes through DNA and RNA alterations. **a.** Full list of 1012 genes that are both frequently and heterogeneously altered across both RNA-and DNA-level alterations. Yellow bars to the left indicate the proportion of samples that had DNA-level alterations while green bars to the right indicate proportion of RNA-level alterations. Middle column is a heatmap corresponding to the −log10(p-value). Asterisks indicate a COSMIC Cancer Gene Census gene (CGC) or PCAWG driver genes. **b.** Same as **a,** but with only CGC or Driver Genes shown. **c.** An enrichment of cancer genes within our list of significantly recurrent genes. **d.** Distribution of alteration types among all significant genes or just CGC or PCAWG driver genes.

Among the top 5% (51/1012) of our ranked genes is *CDK12* (rank 40), which is impacted by multiple but non-overlapping types of alterations. Alterations within its protein kinase domain have been shown to cause dysregulation of DNA repair in cancer^114–117^. Aggregating over all alterations, we find 87 samples that have an alteration in this domain, with 64 (74%) patients having no DNA, but only a RNA alteration in the domain. The most frequent alteration in this gene is an alternative promoter event, where the alternative promoter leads to a truncated transcript of *CDK12*, removing a majority of the kinase domain. Fusion and splice events lead to additional disruptions of the same domain. While we have not selected for mutual exclusivity, we find that the alterations in this gene are marginally mutually exclusive (P=0.07, *WExT*^*118*^). This gene has previously also been classified as cancer census^79^ and PCAWG driver gene^80^. This example illustrates that performing a recurrence analysis over diverse RNA and DNA alterations can help to identify genes known to be important in tumorigenesis.

Our recurrence analysis of heterogeneous RNA alterations also identifies 894 genes that are neither known cancer census nor driver genes. This includes IRF5^119–122^, ZFAT^123^, BCAS3^124,125^, KLF13^126^, TLK2^127–129^, and COL6A3^130,131^, providing new hypotheses for follow-up studies. Those genes may have received less attention because they harbor only rare DNA alterations. The results of our study can also help to understand which parts of the transcripts are altered and functionally affected.

## Discussion

Here we present the most comprehensive catalog of RNA-level alterations in cancer, spanning 27 different tumour types, and provide a harmonised resource of matched transcriptome and whole-genome sequences. In particular, our catalogue includes gene and transcript expression, splicing, alternative promoter, fusion and their association with somatic DNA-level alterations across more than one thousand patients.

We identified 1,012 genes that are recurrently altered by multiple mechanisms (both on DNA or RNA level), which were enriched for known cancer census genes and driver genes as identified in the PCAWG cohort. The list of recurrently altered genes includes genes that are primarily altered at the DNA level (such as *TP53*), but also genes that are most frequently altered on the RNA level (such as *IGF2^132^*). Through the analysis of RNA alterations, we can identify additional gene alterations in samples that otherwise were thought to have no driver alterations based on DNA level analysis alone. In fact, out of 87 samples from the PCAWG study that did not have a driver alteration at the DNA-level ^133^ and also had RNA-Seq data, every sample had an RNA-level gene alteration that was identified through our recurrence analysis. Although cancer is thought to primarily be driven by changes in DNA, such somatic changes may lead to systematic changes on RNA level. Even though, it is also possible that most of the detected RNA alterations, like most somatic mutations, are passenger events, the cancer gene enrichment/pathway analysis suggests that a subset is likely to contribute to cancer. Follow-up validation studies are needed to determine which RNA alterations are functional.

Our work represents comprehensive analysis of gene expression regulatory variation across human cancers, combining eQTL mapping, ASE analysis and expression-linkages with somatic signatures. We identified germline eQTL for around 19% of expressed genes (3,509 out of 18,898), out of which only a small proportion of 426 genes appeared to be specific to cancer (defined as not found in GTEx). This suggests that the germline regulatory variants are largely retained in disease, although some cancer-specific regulatory effects could play important roles. Building on ASE quantifications, we attributed allelic expression dysregulation to distinct classes of genetic variations, showing that the allelic copy number imbalance is a major determinant of allele-specific expression dysregulation in cancer. We have also mapped individual linkages between somatic aberrations in *cis* and individual genes (649 out of 18,898), where 68.4% of associations were between non-coding somatic variants and gene expression (272 intergenic, 172 intronic). In addition to *cis* associations, we have also identified global *trans* effects between somatic signatures and gene expression levels, thereby deriving *de novo* annotations of signatures with previously unknown roles.

We have specifically looked at the effects of somatic mutations in gene promoter regions. Except for the *TERT* promoter, recurrent promoter mutations are rare and we did not observe a strong association with expression nor with alternative promoter usage. The analysis of alternative promoters relies on isoform quantification, which has several limitations and is generally less robust than gene-level analyses. Nevertheless, the recurrence analysis of RNA phenotypes suggests that even without a significant association with promoter mutations, they still contribute to an altered cancer transcriptome.

We identify a broad relationship between somatic alterations and splicing changes looking at mutations further into the intron than was feasible with exome sequencing. We found enrichments of impactful mutations on splicing in extended splice site regions, the polypyrimidine tract, and the branch site position. Overall, we identify 1,872 splicing associated variants (SAVs) that are characteristically rare. We also found 3,941 novel cancer-specific exons that can be partially explained through mutation-driven exonization, including in the tumor suppressor *STK11*. Enrichments of exonization events overlapping to Alu sequences provides a link between genome evolution and cancer evolution and give hints on the exonization mechanism. These insights illustrate the power of whole genome sequencing data studied in this project.

This study, also for the first time, systematically compares and integrates gene fusions with whole genome rearrangements across many tumour types. Most of the novel fusions were found in only one sample and the low sample numbers make it difficult to distinguish passengers from drivers. On the other hand, the promiscuous fusion gene partners were often linked to cancer related pathways, thereby indicating a possible functional role in cancer. While ∼36% of all detected fusion transcripts were predicted to be in-frame, several UTR-mediated fusion transcripts preserve complete coding sequences of one fusion partner. These include a known fusion *TBL1XR1–PIK3CA* in a breast tumour and a notable novel example *CTBP2-CTNNB1* in a gastric tumour. About 82% of the detected fusions can be associated with specific genomic rearrangements. For the remaining fusions, it is entirely possible that the relevant genomic rearrangements have not been detected, or it is also possible that some fusions happen directly on RNA level, as trans-splicing or readthrough events. The availability of whole genome sequences in addition to RNA, allowed us to develop a systematic classification of fusion events, and proposed a novel bridged fusion mechanism to explain how genome rearrangements can lead to a gene fusion.

We show that global differences in RNA expression phenotypes are largely tissue-specific; therefore our ability to associate mutations in *cis* or *trans* are limited by the small sample size within each histotype. Although genetic associations to individual RNA phenotypes were rare, we found that genes altered through heterogeneous mechanisms were enriched for cancer genes, highlighting the importance of examining multiple ways that cancer genes can be altered. We recognize that further work is necessary to investigate additional mechanisms of genome alteration that can lead to changes in RNA such as epigenetic changes ^134^ or enhancer hijacking (e.g., ^135,136^), and our work points to the importance of further investigations. Overall, our analysis shows diverse modes of alteration of cancer genes and pathways at the DNA and RNA level and reveals underappreciated mechanisms of cancer genome alterations confirmed through RNA changes. This demonstrates that RNA analysis reveals cancer-associated pathway alterations that have not yet been detected via exclusive DNA analysis.

## Methods

### RNA-Seq alignment and quality control analysis

Normal and tumour ICGC RNA-seq data, included in the PCAWG cohort^78^, was aligned to the human reference genome (GRCh37.p13) using two read aligners: STAR ^137^ (version 2.4.0i, 2-pass), performed at MSKCC and ETH Zürich, and TopHat2^138^ (version 2.0.12), performed at the European Bioinformatics Institute. Both tools used Gencode (release 19) as the reference gene annotation. For the STAR 2-pass alignment, an initial alignment run was performed on each sample to generate a list of splice junctions derived from the RNA-seq data. These junctions were then used to build an augmented index of the reference genome per sample. In a second pass, the augmented index was used for a more sensitive alignment. Alignment parameters have been fixed to the values reported in (https://github.com/akahles/icgc_rnaseq_align). The TopHat2 alignment strategies also followed the 2-pass alignment principle, but was performed in a single alignment step with the respective parameter set. For the TopHat2 alignments the irap analysis suite ^139^ was used. The full set of parameters is available along with the alignment code in (https://hub.docker.com/r/nunofonseca/irap_pcawg/). For both aligners, the resulting files in BAM format were sorted by alignment position, indexed, and are available for download in the GDC portal (https://portal.gdc.cancer.gov/) and in the Bionimbus Data Cloud (https://bionimbus-pdc.opensciencedatacloud.org/pcawg/). The individual accession numbers and download links can be found in the PCAWG data release table: http://pancancer.info/data_releases/may2016/release_may2016.v1.4.tsv.

Quality control of all datasets was performed at three main levels: i) assessment of initial raw data using FastQC ^140^ (version 0.11.3); ii) assessment of aligned data (percentage of mapped and unmapped reads for both alignment approaches); iii) quantification (by correlating the expression values produced by the STAR and TopHat2 based expression pipelines) (**Extended Data Figure 1**). In total we defined 6 QC criteria to assess the quality of the samples. We marked a sample as a candidate for exclusion if: (1) three out five main FastQC measures (base wise quality, k-mer overrepresentation, guanine-cytosine content, content of N bases and sequence quality) did not pass; (2) more than 50% of reads were unmapped or less than 1M reads could be mapped in total using the STAR pipeline; (3) more than 50% of reads were unmapped or less than 1M reads could be mapped in total using the TopHat2 pipeline; (4) we measured a degradation score ^141^ greater than 10; (5) the fragment count in the aligned sample (averaged over STAR and TopHat2) was less than 5M; (6) the correlation between the expression counts of both pipelines was less than 0.95. If a sample did not pass one of these six criteria it was marked as problematic and were placed on a graylist. If more than two criteria was not passed, we excluded the sample.

A subset of 722 libraries from the projects ESAD-UK, OV-AU, PACA-AU, and STAD-US were identified as technical replicates generated from the same sample aliquot. These libraries were integrated post-alignment for both the STAR and the TopHat2 pipelines using samtools^142^ into combined alignment files. Further analysis was based on these files. Read counts of the individual libraries were integrated to a sample-level count by adding the read counts of the technical replicates.

Initially, a total of 2,217 RNA-seq libraries were fully processed by the pipeline. QC filtering and integration of technical replicates (722 libraries) gave a final number of 1,359 fully processed RNA-seq sample aliquots from 1,188 donors.

### GTEx data analysis

For a panel of normal RNA-Seq data from a variety of tissues, data from 3,274 samples from GTEx (phs000424.v4.p1) were used and analyzed with the same pipeline as PCAWG data for quantifying gene expression. A list of GTEx identifiers are provided at https://www.synapse.org/#!Synapse:syn7596611.

### Quantification and normalization of transcript and gene expression

STAR and TopHat2 alignments were used as input for HTSeq^14^ (version 0.6.1p1) to produce gene expression counts. Gencode v19 was used as the gene annotation reference. Quantification on a per transcript level was performed with Kallisto^143^ (version 0.42.1). This implementation is available as a Docker container at https://hub.docker.com/r/nunofonseca/irap_pcawg. The implementation of the STAR and TopHat2 quantification is available as Docker containers in:

- https://github.com/akahles/icgc_rnaseq_align
- https://hub.docker.com/r/nunofonseca/irap_pcawg/

, respectively. Consensus expression quantification was performed by taking the average expression based on STAR and TopHat2 alignments. Gene counts were normalized by adjusting the counts to fragments per kilobase of million mapped (FPKM)^144^ as well as fragments per kilobase of million mapped with upper quartile normalization (FPKM-UQ) where the total read counts in the FPKM definition has been replaced by the upper quartile of the read count distribution multiplied by the total number of protein-coding genes.

The FPKM and FPKM-UQ calculations were:

- FPKM = (C.10^9^) / (N.L) where N = total fragment count to protein coding genes, L = length of gene, C = fragment count
- FPKM-UQ = (C.10^9^) / (UQ.L.G) where UQ= upper quartile of fragment counts to protein coding genes on autosomes unequal to zero, G = number of protein coding genes on autosomes

### t-SNE analysis

The t-SNE plot in Figure 2D was produced using the RTsne package^145^ (with a perplexity value of 18) based on the Pearson correlation of the aggregated expression (log+1) of the 1,000 most variable genes. FPKM expression values per gene were aggregated (median) by tissue (GTEx) and study (PCAWG). Coefficient of variation for each gene was also computed per tissue (GTEx) and study (PCAWG) to determine the 1,000 most variable genes. Purity values were obtained from syn8272483.

### Associations between genetic variation and gene expression

#### Patient Cohort

We analyse the tumor whole genome sequencing (WGS) and matched RNA-Seq data of 1,188 patients from the Pan-cancer Analysis of Whole Genomes (PCAWG) cohort. Germline genotypes, SNV calls and segmented allele-specific somatic copy-number alteration (SCNA) calls were obtained from^105,146^. We matched 1,188 tumor RNA-seq IDs^147^ to WGS white-list tumor IDs (synapse entry syn10389164). For patients with multiple WGS IDs (2 out of 1,188) or RNA-seq aliquot IDs (17 out of 1,188), we resolved the matching by pairing samples with the same *tumor_wgs_submitter_specimen_id* (**Supplementary Table 1**). The 1,188 patients are spread across 27 cancer types and 29 project codes and include 899 carcinomas; 34 patients are metastatic and 13 recurrent with the remaining patients being primary tumors (**Supplementary Table 1**).

We used the data of these 1,188 patients for performing somatic and germline expression quantitative trait loci (eQTL) mapping, allele-specific expression (ASE) analysis and association studies between gene expression and mutational signatures.

#### Gene expression

Gene expression values (measured in FPKM) were obtained from the PCAWG-3 group (PCAWG-3 Tophat2/Star gene expression, syn5553985). Genes with FPKM ≥ 0.1 in at least 1% of the patients (12 patients) were retained, resulting in 47,730 genes. Only 18,898 protein-coding genes (according to the *gene_type* biotype reported in Gencode v.19) were used for the subsequent QTL analyses. Log2 expression values were subjected to peer analysis ^148^ to account for hidden covariates (syn7850427). In order to balance number of covariates, statistical power and available sample sizes per cancer type, we followed the GTEx protocol and estimated 15, 30 and 35 hidden covariates to be used depending on sample size (N<150, 150≤N<250, N≥250) ^149^. Peer residuals were then rank standardised across patients. The FPKM cutoff and peer correction were also applied to the subset of 899 carcinoma patients, yielding 18,837 protein-coding genes after filtering.

#### Covariates

In all linear models, we accounted for known confounding factors by modeling them as fixed effects. In all association studies, we accounted for sex, project code (describing cancer type and country of origin) and per gene copy-number (CN) status (**Supplementary Table 1** for the list of per patient covariates; syn7253568 and syn7253569 for sex and project codes; syn9661460 for per gene CN). Per gene CN alterations were derived as the average copy number across all copy number aberrations called within the annotated gene boundaries based on syn8042988.

The somatic eQTL, ASE and mutational signature analyses additionally accounted for total somatic mutation burden (number of single nucleotide variants (SNVs) and short insertions and deletions (indels)) and sample purity (**Supplementary Table 1**). Purity was estimated based on CN segmentation. In addition, the somatic eQTL and ASE analyses accounted for local SNV burden calculated in a 1Mb window from the gene coordinates (syn8494689 for *cis* somatic burden).

The germline eQTL analysis additionally modelled the population structure as random effect. The population structure was assessed by a kinship matrix that was calculated based on every 20th germline variant, processed as described below (see Germline variants). The kinship matrix was then calculated as an empirical patient-by-patient covariance matrix.

**Table 1.**
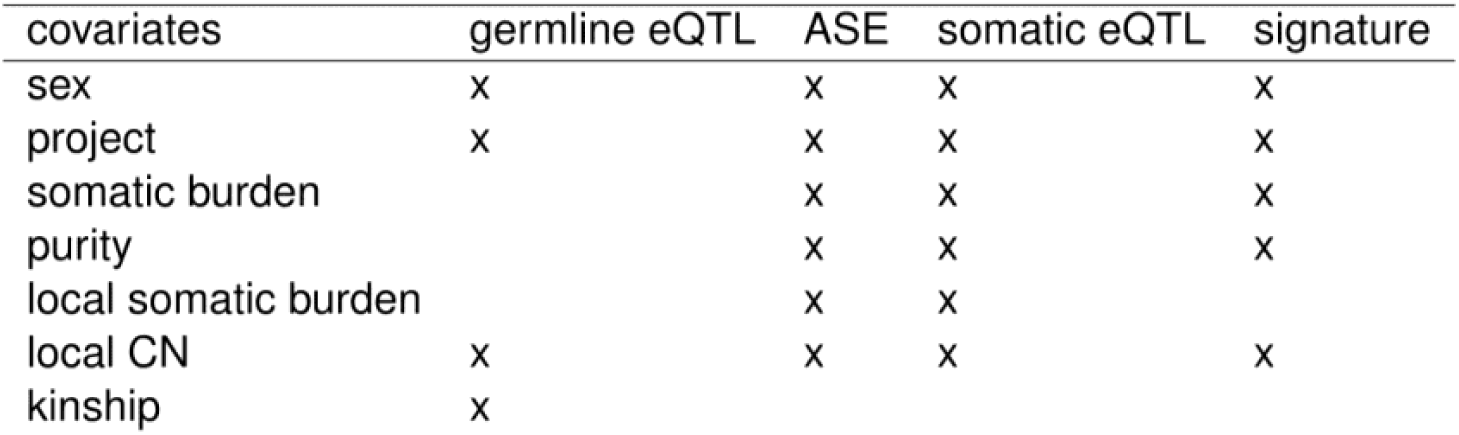
Covariates that were accounted for per analysis method. The project code describes cancer type and country-of-origin. Somatic burden is the total number of SNVs and indels. Purity was estimated based on CN segmentation. Local somatic burden is the number of SNVs in a 1Mb window around the gene coordinates. Local CN was defined as the average CN state across all SCNAs called within the annotated gene boundaries.

#### Gene Ontology and Reactome Pathway enrichment

We performed Gene Ontology^150,151^ and Reactome Pathway^152,153^ enrichment with the Bioconductor packages biomaRt^154,155^, clusterProfiler^156^, and ReactomePA^157^ (FDR ≤ 10%). The number of genes used as background set is described per analysis method.

### Germline eQTL

#### Germline Variants

PCAWG variant calls v0.1^105^ were downloaded from GNOS and processed following the PCAWG-8 protocol:

1. VCF files were indexed and merged using bcftools^158^.
2. All variants were filtered for *PASS* flag.
3. All variants were filtered for quality larger than 20.
4. Only bi-allelic sites were considered.

HDF5 files for each 100kb chunk of the VCF files were generated, assuming additivity that was numerically encoded as 0/1/2 for homozygous reference/heterozygous/homozygous alternative state. For indels, we encoded the presence or absence of the variant as 0/1. Each variant was normalized to mean 0 and standard deviation 1. Missing variants were mean-imputed. To create our eQTL release set v1.0, the resulting HDF5 files were subsequently merged into a global HDF5 file and all variants which follow any of the following conditions were removed:

1. Minor allele frequency (MAF) ≤1%
2. Missing values ≥ 5%

#### Germline eQTL analysis

In the germline eQTL analyses, we used the processed gene expression dataset from 1,178 patients for which germline variant calls (eQTL release set v1.0, see Germline Variants) were available. Linear mixed models were used to model the correlation between germline variants (within 100kb from gene boundaries) and gene expression values (see Gene expression) using the limix package^159^. Known covariates were modeled as fixed effects and population structure as random effect (see Covariates).

A two-step approach was employed to adjust for multiple testing. First, for each gene, we adjusted for the number of independent tests estimated based on local linkage disequilibrium (LD)^160^. Second, we performed a global correction across the lead variants, *i.e.*, the most significant SNPs, per eQTL. Germline eGenes were defined as genes with an eQTL with global FDR ≤ 5%.

#### eQTL release set v0.1

The germline eQTL release set v0.1 is a pilot call set that was generated before the eQTL release set v1.0 and was used in our ASE analysis. The eQTL sets v0.1 and v1.0 differ as follows. In v0.1, SNVs and indels from 1,194 patients from the Annai variant call set (syn4231951 and syn4231952) were used, compared to 1,178 patients in v1.0. As these variants were called on a patient-by-patient basis, they were merged and a high missingness call rate per variant was generated due to low quality variants or to the absence of a call from simply being a reference allele. To solve this issue, we kept variants with missingness ≤ 20% across patients and inferred missing variants as the reference allele. Otherwise, we discarded the whole variant. We filtered variants with MAF ≤ 1%. Multiple testing was firstly applied to all variants tested within a *cis* window of 1Mb up and downstream of gene boundaries and then to all lead variants with a Benjamini-Hochberg procedure. Compared to v1.0, more genes were considered since we retained protein coding and non protein coding genes with FPKM ≥ 0.5 in at least 1% of the cohort (N=37,116). We obtained 5,373 eGenes (FDR ≤ 10%). Results of this pilot germline eQTL analysis are available on synapse (syn6157699).

#### GTEx comparative analysis

The GTEx comparative eQTL analysis was based on the eQTL maps version 6p^161^. We mapped the positions and alleles of our PCAWG-specific eQTL to the eQTL in all GTEx tissues. In order to determine whether a lead eQTL variant is replicated in a given GTEx tissue, we followed the following strategy according to Kilpinen *et al.* (2017)^162^. For each eGene, we considered the eQTL lead variant and assessed the replicability of the signal in the GTEx cohort based on marginal association statistics using matching tissue-of-origin in 23 cases (P < 0.01/23) as well as all 42 GTEx tissues without cell lines (P < 0.01/42). If the lead variant did not replicate or was not tested, we determined replication based on the variant with the smallest P value within the LD block (r^2^ ≥ 0.8 estimated based on UK10k) of the lead variant across 23 (or 42) tissue-matched GTEx analyses. If neither lead nor any variant within the LD block were tested, we determined replication based on the smallest P value of any variant within the 100kb window tested within the GTEx cohort.

To assess if the PCAWG-specific eQTL replicate in testis, we repeated our GTEx comparison while including GTEx testis tissue. We found 37 eQTL that only occur in the PCAWG cohort and in testis, but in no other tissue.

### Allele-specific expression analysis

#### Assembling Phased Germline and Somatic Variants

To understand the precise effect of somatic variations in their genomic context and for subsequent allele-specific analyses, both germline and somatic variants were phased. For assembling phased germline genotypes we used the Sanger 1000G callset (PCAWG-11), and applied IMPUTE2 (Howie et al. 2009) for phasing of heterozygous germline variants. The IMPUTE2 output was corrected using results from the Battenberg CN calling algorithm (Nik-Zainal et al. 2012) to ascertain that no haplotype switches occur within regions of consecutive CN gain. The resulting phased germline genotypes were arranged such that haplotype 1 always corresponded to the amplified alleles in regions with SCNAs (major allele). In cases where both co-occur on the same NGS read (∼10M variants, ∼20% of all SNVs), we phased individual somatic variants to the nearest germline heterozygous site. For downstream analyses, we only considered SNVs that were phased by at least three reads to the respective germline variant (∼6M out of 10M SNVs).

All phased SNVs were aggregated into functional categories based on their genomic regions defined by gene annotations (upstream, downstream, promoter, 5’ UTR, intron, synonymous, missense, stop gain, 3’ UTR) and mapped to the nearest gene within a *cis* window of 100kb using the Variant Effect Predictor (VEP) tool^163^. Promoter variants were defined as 1kb upstream of the transcription start site (TSS). We included flanking regions by using the VEP ‘UpDownDistance’ plugin with a maximum range parameter of 100kb. We divided the upstream and downstream variant categories into disjoint categories using 10kb windows from 10 to 100kb. We integrated ‘splice donor’ and ‘splice acceptor’ variants into the general ‘splice region’ variant category and mapped ‘stop retained’ variants to the ‘synonymous’ variant category. We averaged transcript-level annotations to gene-level annotations to retrieve the expected functional effect of a variant for a given gene. We analysed the relationship between SNV variant allele frequency (VAF) and SCNAs at the same locus to determine whether variants occurred before (‘early’) or after (‘late’) the corresponding SCNA (PCAWG-11). We computed a weighted *cis* mutational burden per category by estimating the cancer cell fraction (CCF) of each SNV and aggregating SNVs to a total localised burden weighted by their respective CCF.

#### ASE read counts

The positional information of the heterozygous germline variants was used together with the RNA-seq BAM files as input to the GATK ASEReadCounter (Castel et al. 2015) algorithms for counting ASE reads. We considered reads with a minimum mapping quality of 20 and a minimum base quality of 10. Only heterozygous variants with a minimum coverage of eight RNAseq reads were considered for all further analyses.

The raw ASE read counts were post-processed as follows: (1) ASE sites were converted to BED files and aligned against the ENCODE 50mer mappability track (wgEncodeCrgMapabilityAlign50mer.bigWig) to extract mappability scores for all sites. All sites with mappability scores unequal to 1 were removed. (2) All sites with allelic read counts less or equal to 1 were removed to prevent genotyping error to influence ASE quantification. (3) All sex chromosomes were dropped for further analysis. (4) We estimated sequencing error per patient as the sum of non-reference and non-alternative bases over the total number of bases. We assessed statistical mono-allelicity through a binomial test using the estimated sequencing error probabilities, corrected using the Benjamini-Hochberg step down procedure. All sites that appeared to be statistically mono-allelic were removed. (5) For each ASE site, CN states were retrieved from the Sanger CN consensus callset (PCAWG-11). Purity estimates for each patients were retrieved from the accompanying purity tables.

To aggregate site-level ASE to a gene-level readout and to allow for estimation of effect directionality, we used the phased germline genotypes. Gene mapping was performed against ENSEMBL release 75 using the pyEnsembl Python library. We retrieved all genes at each ASE site and summed up the read counts on the respective haplotypes to gene-level haplotype-specific read counts. We further averaged haplotype-specific CN states to a mean haplotype-specific CN state per gene and computed the gene-level CN ratio as the major over total ratio of those averages. To allow for a robust assessment of gene-level ASE we only considered genes with at least 15 reads total, yielding 4,379,378 gene-patient pairs of 1,120 patients and 17,009 unique genes across 12,441,502 accessible sites in total. Every remaining gene was tested for allelic expression imbalance (AEI) using a binomial test against an expected read ratio of 0.5 to derive nominal P values, and a binomial test against the expected CN ratio modified by tumor purity to derive CN-corrected P values. Nominal and CN-corrected P values were adjusted separately for multiple testing using the Benjamini-Hochberg procedure. Significant AEI was called at FDR ≤ 5%. We further annotated each gene with the number of ASE sites used for aggregation. For all downstream analyses, we only considered genes annotated as protein coding (ENSEMBL biotype=’protein_coding’).

#### Generalized linear models

Across all 4,379,378 gene-patient pairs, we trained multivariate linear models using (i) logistic regression against a binary indicator of AEI absence or presence in a gene or (ii) standard linear regression against the phased ASE ratio of a gene to assess the directionality of the regulatory change. For (i), haplotype-specific mutations were summed up to a total burden per category, whereas for (ii), we used the difference in burden between the haplotypes 1 and 2. The consistency of the phasing map between somatic variants and ASE sites ensured that model coefficients kept their directionality independent of the arbitrary labelling of haplotypes as 1 or 2. The full set of considered factors is as follows:

- CN ratio at the gene locus (0.5 ≤ x ≤ 1)
- Sample purity (0 < x < 1)
- Natural logarithm of total gene length (x > 0)
- Natural logarithm of the length of the canonical transcript (x > 0)
- Heterozygosity of the lead eQTL variant (x = 0 if homozygous, x = 1 if not homozygous)
- All mutsational burden categories as determined by VEP annotations (upstream in 10kb windows, downstream in 10kb windows, promoter, 5’ UTR, intron, synonymous, missense, stop gain, 3’ UTR; x ≥ 0 for logistic model, x ∈ ℝ for directed model)

To compare global effects and different contributions of SCNA, germline eQTL, coding and non-coding SNVs, a simplified logistic model was trained after accumulating all coding and non-coding variants to separate categories and reporting standardised effect sizes (**Figure 5e**).

#### Cancer Gene Enrichment

Cancer Gene Enrichment was conducted on the COSMIC census (Forbes et al. 2017) using Fisher’s exact test and Gene Set Enrichment Analysis (GSEA) as described in the manuscript. For enrichment, the average score of a gene was computed across the cohort and only genes with at least five replicates in the cohort were kept, yielding a total of 16,078 genes.

#### Chromosomal distribution of ASE

We calculated the recurrence of ASE genes in each tumor type. To examine the chromosomal distribution of ASE genes, we calculated the average recurrence of all genes for every 200-gene window with a 10-gene step, and then subtracted the average ASE occurrence in each tumor type to obtain the peaks of ASE surplus across all chromosomes. The recurrence of CN genes was calculated in an analogous manner.

### Somatic eQTL

#### Somatic calls and mutational burden

We used the set of consensus SNVs somatic calls provided by PCAWG (syn7357330) based on three core caller pipelines and MuSE^164^. On average, we counted 22,144 somatic SNVs per patient, with different median numbers of SNVs per cancer type, ranging from 1,139 in thyroid adenocarcinoma to 72,804 SNVs in skin melanoma (**Extended Data Figure 17a**). Due to the low frequency of somatic SNVs across the cohort (**Extended Data Figure 17b**), we collapsed the variants by genomic regions defined by gene annotations (Gencode v.19^165^). Specifically, we generated a set of disjoint gene exons by collapsing overlapping exon annotations into single features using bedtools^166^. The set of disjoint introns was generated using bedtools by subtracting the collapsed exonic regions from the gene regions. To map local effects of somatic mutations in flanking features outside the gene body, we binned the surrounding regions (plus and minus 1Mb from the gene boundaries) into 2kb windows overlapping by 1kb.

We defined three different types of aggregated somatic burden to assess differences in power in detecting somatic eGenes and P value calibration. The burden in a genomic region was defined as (1) a binary value that indicates presence or absence of SNVs, (2) the aggregated burden as sum of SNVs, or as (3) weighted burden, *i.e.* sum of VAFs of the SNVs (**Extended Data Figure 18 a**). Genotypes were standardised across patients (to mean zero and standard deviation one) and effect sizes of the association study were divided by the corresponding genotype standard deviation.

We assessed calibration of all three analyses with QQ-plots of nominal and permuted P values (permutation of the patients in the gene expression matrix) (**Extended Data Figure 18 b-d**).

#### Somatic eQTL analysis

Linear models were used to model the correlation between recurrent somatic burden and gene expression of up to 18,898 protein-coding genes, using the limix package^167^ (see Gene expression). Gene expression was corrected for 35 hidden peer factors. Known covariates were modeled as fixed effects (see Covariates).

The somatic eQTL analysis was performed on all 1,188 patients and on the subset of 899 carcinoma patients (representing 20 of the 27 cancer types) to replicate the analysis on a more homogeneous set of tumors. A *cis* window of 1Mb from the gene boundaries (Gencode v.19) to find mutated genomic intervals with a burden frequency ≥ 1% in the cohort (at least 12 patients in the full cohort and 9 patients in the carcinoma cohort). Altogether, 18,708 of the genes had at least one mutated interval at that frequency and were included in the analysis.

Bonferroni correction was applied to correct for multiple *cis* windows tested within the same gene. Then, Benjamini-Hochberg correction was applied to adjust the P values of the lead genomic regions across genes. Somatic eGenes were defined as genes with an eQTL at an FDR ≤ 5%.

The analysis of co-localization of SVs and somatic burden was performed per aliquot id, looking for the closest SV to the leading genomic interval identified for each eGene, using the consensus WGS-based somatic structural variants (version 1.6; <u>syn7596712</u>).

#### Functional enrichment in somatic *cis* eQTL

To identify putative regulatory sites enriched for somatic eQTL, we retrieved functional annotations of the lead genomic flanking intervals of the somatic eQTL (638 somatic eQTL). Therefore, we mapped somatic eQTL to 25 Roadmap Epigenomics chromatin marks of 127 different cell types ^168^ and ENCODE transcription factor binding site (TFBS) annotations in 9 cell types (including 8 cancer and one embryonic stem cell lines^169^; **Supplementary Table 5 and 6**). We compared annotations in the significant set of eQTL with a null distribution based on a 1k random samplings of a matched set of genomic intervals. To define the matched sets of genomic intervals, we selected flanking genomic intervals from the whole set of tested genes that showed a similar distance from the gene start (exact distance ±2kb) and that matched the exact burden frequency of the corresponding interval in the significant associations. We then overlapped the 1k matched sets with Roadmap Epigenomics and ENCODE annotations. To avoid ambiguous overlaps (with multiple annotations), we only retained genomic intervals showing a minimum overlap of 10% of their length.

We retrieved an empirical P value of enrichment for each annotation by counting the number of randomly sampled flanking intervals (N) showing greater number of overlaps compared to the eQTL set (P = (N+1)/(1000+1)). Benjamini-Hochberg correction was applied to the empirical P values (over 25 marks in 127 cell lines for Roadmap Epigenomics annotations and over 149 TFBS for 9 ENCODE cell lines). We then computed the fold change per annotation and cell line as ratio of annotated lead flanking intervals and mean number of annotated matched random flanking intervals over the 1k samplings.

#### Variance Component Analysis

Limix was used to perform variance decomposition using the same covariates as in the somatic variant analyses except for local CN state (see Covariates). The random effects were based on the following germline variants and somatic burden (see Somatic calls and mutational burden for detailed description of burden):

- *cis* somatic intronic: weighted burden in introns
- *cis* somatic exonic: weighted burden in exons
- *cis* somatic flanking: weighted burden in 1kb-overlapping regions of 2kb within 1Mb from gene boundaries
- somatic intergenic: weighted burden in 1kb-overlapping regions of 2kb outside the 1Mb window
- *cis* germline: germline variants within 100kb from gene boundaries
- *trans* germline: genome-wide population structure (see Covariates)
- Local CN variation (see Covariates)

All the data was mean-centered and standardised. For each of the random effects, a linear kernel was computed and used as covariance matrix. The resulting variance components were normalized to add up to one.

### Mutational signature associations

We obtained 39 mutational signatures from PCAWG-7 beta 2 release (PCAWG7, 2017) and used linear models to associate the mutational signatures with gene expression of up to 18,898 protein-coding genes across 1,159 patients while accounting for known covariates (see Covariates) (quality control: **Extended Data Figure 25a-e**). The 1,159 patients were a subset of the total 1,188 patients, for which mutational signature profiles were available. Gene expression was corrected for 35 hidden peer factors (see Gene expression).

We retained 18,888 genes that showed a minimum FPKM of 0.1 in at least 1% of 1,159 the patients (see Gene expression). Signatures with zero variance and a prevalence below 1% were filtered. Like this, we obtained 28 signatures. We applied linear models to associate expression of these genes with the signatures across all 1,159 patients, a subset of 877 carcinoma patients or a subset 891 European patients to assess consistency of the associations (**Extended Data Figure 25f-g**).

Across all patients, we found 1,176 significantly associated genes after Benjamini-Hochberg correction (we used an FDR ≤ 10% for enrichment analyses, multiple testing was applied across all signature-gene pairs, **Supplementary Tables 15a-c**). We performed gene enrichment analyses of the significant genes per signature (see Gene Ontology and Reactome Pathway enrichment, here 18,831 background genes, multiple testing correction across all ontologies per signature FDR ≤ 10%, **Supplementary Table 15d**). Whereas most signatures only affected few genes, 18 showed recurrent *trans* effects and affected expression of over 20 genes (**Extended Data Figure 26d, Supplementary Table 15e**). We further found that the vast majority of genes (85.8%) were associated with only one signature (1,009 genes); 129 genes were associated with two, 32 with three, five with four and one with five signatures.

To assess how tissue-specific both mutational signatures and their associations with gene expression are, we analysed the occurrence of each signature in each of the cancer types. We assessed the presence (at least one SNV of a signature in at least one patient with a specific cancer type) and mean prevalence (mean number of SNVs of a certain signature across all patients of a specific cancer type) of the signatures in the cancer types (**Extended Data Figure 28**). We defined cancer type-specific signatures to occur in up to 4 cancer types (signatures 4, 7, 9, 12, 16, 38 and 39) and common signatures to be missing in up to 5 cancer types (signatures 2, 13, 18). For each of these signatures, we performed cancer type-specific analyses, *i.e.*, we assessed the association between the respective signature and gene expression in just the patients that are of a cancer type that shows mutations of the respective signature (**Extended Data Figure 28** left heatmap). We then correlated the P values of these cancer type-specific analyses with the P values of the analysis across all patients and calculated the Pearson correlation coefficients (**Extended Data Figure 29a-k**). We show that the correlation between cancer type-specific and whole-cohort P values is dependent on the sample size of the respective analysis (r^2^=0.671, **Extended Data Figure 29i**).

We further performed principal component analysis (PCA) on the signatures across both, patients (PCA on signature-specific SNVs per patient) and genes (PCA on adjusted P values of signature-gene expression associations) (**Extended Data Figure 26a-b**).

To assess significance of the functional annotation of SNVs by mutational signatures, we also associated gene expression with the total number of SNVs and correlated the P values (-log_10_P) of the associations with the respective signature-specific P values. The absolute Pearson correlation coefficients remain below 0.1 (**Supplementary Table 15f**).

To establish causality of signature-gene expression associations, we included the germline eQTL into the analysis using linear mixed models; 197 of our 1,176 signature-associated genes were also germline eGenes. These 197 associations involved 26 of the 28 mutational signatures. We associated the lead variants of these eGenes with the rank-standardised signature SNVs across 2,507 patients. We used the subset of the 2,818 WGS patients for which mutational signature profiles and all known covariates were available. We accounted for the same fixed covariates as in the mutational signature-gene expression association studies and, in addition, for kinship as a random effect (see Covariates).

We then performed proportional colocalisation analysis with Bayesian Model Averaging (BMA) using the R package coloc ^170^ to test whether gene expression and mutational signatures share common causal genetic variants in a given gene region. A proportional colocalisation analysis tests the null hypothesis of colocalisation by assuming that two phenotypes that share causal variants will have proportional regression coefficients for either phenotype with any variant selection in the vicinity of the causal variant. We applied the BMA approach, with each tested model consisting of a selection of two variants. The P values are then averaged over all models to generate posterior predictive P values^170^. We filtered variants so that no pair of variants showed r^2^>0.95 and each variant’s marginal posterior probability of inclusion with one of the phenotypes was greater than 0.01. The nominal P values of rejecting the null hypothesis of colocalisation are listed in **Supplementary Table 15e**.

We then performed mediation analysis^108,109^ to assess directionality of the effect between germline eQTL, gene expression and mutational signature. First, Causal Mediation Analysis was applied to each of the triples of eQTL lead variant, gene and mutational signature using a structural equation model from the R package lavaan^171^. Then, we employed the R package mediation^172^ to assess significance of mediation and estimate the proportion of mediated effect by nonparametric bootstrap confidence intervals (1000 simulations).

### Estimation of alternative promoter activity

We estimated promoter activities using RNA-seq data and Gencode (release 19) annotations for 70,937 promoter in 20,738 genes. We grouped transcripts with overlapping first exons under the assumption that they are regulated by the same promoter ^173^. We quantified the expression of each transcript from the RNA-Seq data using Kallisto ^143^ and calculated the sum of expression of the transcripts initiated at each promoter to obtain an estimate of promoter activity. To obtain the relative activity for each promoter, we normalized each promoter’s activity by the overall gene’s expression. More details can be found in ^174^.

### Identification of RNA fusions

Gene fusions between any two genes were identified based on two different gene fusions detection pipelines: FusionMap (version 2015-03-31) pipeline ^175^ and FusionCatcher (version 0.99.6a)/STAR-Fusion (version 0.8.0) pipeline ^176^. The detailed procedure is described in ^177^. ChimerDB 3.0 was used as a reference of previously reported gene fusions. The database contains 32,949 fusion genes splitted into three groups:

- KB : 1,067 fusion genes manually curated based on public resources of fusion genes with experimental evidences;
- - Pub : 2,770 fusion genes obtained from text mining of PubMed abstracts.
- - Seq : archive with 30,001 fusion gene candidates from deep sequencing data. This set includes fusions found by re-analysing the RNA-seq data of the TCGA project encompassing 4,569 patients from 23 cancer types.

Briefly, FusionMap was applied to all unaligned reads from the PCAWG aligned TopHat2 RNA-seq BAM files for each aliquot to detect gene fusions. In the FusionCatcher/STAR-Fusion pipeline, for each aliquot with paired-end RNA-seq reads FusionCatcher was applied to the raw reads, with the genome reference. To reduce the number of false positive fusions, the two sets of fusions were filtered to exclude fusions based on the number of supporting junction reads, sequence homology, occurrence in normal samples (from the GTEx and the PCAWG cohort). To get a high-confident consensus fusion call set from these two pipelines, a fusion to be included in the final set of fusions had to: i) be detected by both fusion detection tools in at least one sample; and/or ii) be detected by one of the methods and have a matched SV in at least one sample. The consensus WGS-based somatic structural variants (version 1.6) were obtained from the PCAWG repository in Synapse (https://www.synapse.org/#!Synapse:syn7596712). Finally, 3,540 fusion events were included as the consensus fusion call set, from these 2,268 were detected by both FusionCatcher/STAR-Fusion and FusionMap (from these 1821 had SV support) and 1,112 were detected by only one method and had SV support. All fusions are available in Synapse (https://www.synapse.org/#!Synapse:syn10003873).

### Identification of alternative splicing

We used the alignments based on the STAR pipeline to collect and quantify alternative splicing events with SplAdder^178^. The software has been run with its default parameters with confidence level 3. We generated individual splicing graphs for each RNA-Seq sample for both tumor as well as matched normal samples (when available). All graphs were then integrated into a merged graph to comprehensively reflect all splice junctions observed in all samples together. Based on this combined graph, SplAdder was used to extract alternative splicing events of the following types: alternative 3’ splice site, alternative 5’ splice site, cassette exon, intron retention, mutually exclusive exons, coordinated exon skip (see Supplemental Figure 3 in ^178^). Each identified event was then quantified in all samples by counting split alignments for each splice junction in any previously identified event and the average read coverage of each exonic segment involved in the event was determined. We then computed a percent spliced in (PSI) value for each event that were then used for further analysis. We further generated different subsets of events, filtered at different levels of confidence, where confidence is defined by the SplAdder confidence level (generally 2), the number of aligned reads supporting each event, the number of samples that were found to support the event by SplAdder, and the number of samples that passed the minimum aligned read threshold.

### Identification of RNA editing events

We used an RNA editing events calling pipeline, which is an improved version of a previously published one ^179^. Firstly, we summarized the base calls of pre-processed aligned RNA-reads to the human reference in pileup format. Secondly, the initially identified editing sites were then filtered by the following quality-aware steps: (1) The depth of candidate editing site, base quality, mapping quality and the frequency of variation were taken into account to do a basic filter: the candidate variant sites should be with base-quality ≥ 20, mapping-quality ≥ 50, mapped reads≥ 4, variant-supporting reads ≥3, and mismatch frequencies (variant-supporting-reads/mapped-reads) ≥ 0.1 (2) Statistical tests based on the binomial distribution B(*n, p*) were used to distinguish true variants from sequencing errors on every mismatch site ^180^, where *p* denotes the background mismatch rate of each transcriptome sequencing, and *n* denotes sequencing depth on this site. (3) Discard the sites present in combined DNA SNP datasets (dbSNP v.138, 1000 Genome SNP Phase 3, human populations of Dutch^181^, and BGI in-house data. (Combined datasets deposited at: ftp://ftp.genomics.org.cn/pub/icgc-pcawg3) (4) Estimate strand bias and filter out variants with strand bias based on two-tailed Fisher’s exact test. (5) Estimate and filter out variants with position bias, such as sites only found at the 3’-end or at 5’- end of a read. (6) Discard the variation site in simple repeat region or homopolymer region or <5 bp from splicing site. (7) To reduce false positives introduced by misalignment of reads to highly similar regions of the reference genome, we performed a realignment filtering. Specifically, we extracted variant-supporting reads on candidate variant sites and realign them against a combination reference (hg19 genome + Ensembl transcript reference v75) by bwa0.5.9-r16. We retain a candidate variant site if at least 90 % of its variant-supporting reads are realigned to this site. Finally, all high confident RNA editing sites were annotated by ANNOVAR^182^. (8) To remove the possibility of an RNA editing variant being a somatic variant, the variant sites are positionally filtered against PCAWG WGS somatic variant calls. The calls were obtained from the PCAWG repository in Synapse (https://www.synapse.org/#!Synapse:syn7364923). (9) The final two steps of filtering are designed to enrich the number of functional RNA editing sites. Firstly, we only keep events that occur more than two times in at least one cancer type. Secondly, we only keep events that occur in exonic regions with a predicted function of either missense, nonsense, or stop-loss.

### Gene-Centric table creation

To perform the outlier recurrence analysis, each alteration type was condensed into a binary gene-centric format. Since alterations can occur at many different levels (nucleotide, exonic, gene, or transcript), to make them comparable we projected each alteration type onto the gene body. We summarised each alteration type by its presence or absence within a single gene, yielding a binary value per type for each gene-sample pair. The events we included in this analysis were: RNA editing, non-synonymous variants, expression, splicing alterations, copy-number alterations, fusions, and alternative promoters. Each alteration type was summarised differently due to their inherent differences. RNA editing events and non-synonymous variants can occur several times within a single gene body, so these events were denoted as 1 if they occurred at least once within a gene-sample pair.

For copy-number, to obtain a single numerical value per gene-sample pair the copy-number alteration was averaged over the gene body. Since we do not have matched normal samples against which to compare, we instead consider outlying events within each histotype as significant. Thus, a value of 1 was given to average copy-number alterations larger than 6 or smaller than 1.Similarly to non-synonymous variants, multiple splice events can occur within a gene body. The event with the most extreme percent-spliced in (PSI) within the gene-body is selected as the candidate event for the gene. The candidate’s PSI value for a gene is compared over all samples within a histotype and it is set to 1 (*i.e.*, significant) only if it is in the top or bottom 1% within that histotype. We used the same binarization method for expression outliers. Alternative promoter outliers were calculated based on relative promoter activity within each cancer type. To binarize the promoter activity, a z-score cut-off of two over the relative expression distribution within each cancer type was used.

For allele specific expression outliers, only genes significant allelic imbalance (FDR ≤ 5% and allelic imbalance > 0.2, binomial test) were denoted as 1.

### Identifying genes with heterogeneous mechanisms of alterations in *cis*

Genes with multiple heterogeneous mechanisms of RNA alteration were identified from associations of cis-variants with gene expression, allele-specific expression, fusions, and splicing. For gene expression, genes associated with somatic eQTL with FDR < 5% were selected. For allele-specific expression, the top 5% of genes ranked by the predicted contribution of somatic variants on ASE. For fusions, all RNA fusions with structural variant support were selected. For splicing, genes having somatic mutations within 10bp of an annotated splice site or 3bp of a branchpoint and associated splicing were selected. These associated splicing events also had to have a |Z-score| greater than or equal to 3 and the difference of percent spliced in the outlier event was greater than or equal to 10%.

### Recurrence analysis

The recurrence analysis was performed on the binarized gene-centric table for all nine alteration types. The recurrence analysis was performed in three main steps: 1) Aggregate within each alteration type across all samples. This results in a sum for each gene-alteration pair. 2) Convert the counts to ranks within each alteration. The smallest rank goes to the most frequently altered genes. Ranks are split evenly across ties. 3) To generate a single score for each gene, the second smallest rank across alterations is used as the score. To identify a score cutoff for significantly altered genes, a null distribution was generated through permutation. The permutations were performed over the samples within each gene-alteration pair, this was done over all genes and samples 1000 times, concatenating together all observations, results in 16.8M permuted scores. A P value less than 0.05 as derived from the null distribution was defined as significant, resulting in a score greater than or equal to 706.5 considered as significant.

### Co-occurrence analysis

The co-occurrence analysis was also performed on the aforementioned binarized gene-centric table, but only including variants, expression outliers, alternative promoters, alternative splicing and fusions. SCNA and ASE are excluded due to a large number of anticipated co-occurrence. In this analysis, we required at least one gene of a given alteration pair to be a COSMIC gene.

For each alteration pair, based on the number of donors with both alterations, one alteration only and neither alterations in a set of cancer samples, we performed Fisher’s exact test to determine whether the alteration pair was independent of each other. Such tests were followed by Benjamini-Hochberg multiple testing correction to obtain the FDR (or q-values). To rule out the potential false positive association caused by tissue specific alterations, we performed the same analysis for each of the tumor types with at least 50 patients, and only retained those alteration pairs which were significantly associated in both the pan-cancer analysis and in at least one specific cancer indication. Among the significantly associated alteration pairs, the co-occurred pairs were those with odds ratio greater than 1. Pathway enrichment and visualization^96,183^ was conducted using the R package ReactomePA^184^. The circos plots were generated using the R package circlize^185^. The splicing related genes were derived from the genes annotated as “REACTOME_MRNA_SPLICING” or “REACTOME_MRNA_SPLICING_MINOR_PATHWAY” in the Molecular Signatures Database (MSigDB)^186^.

## Extended data figures

**Extended Data Figure 1.**
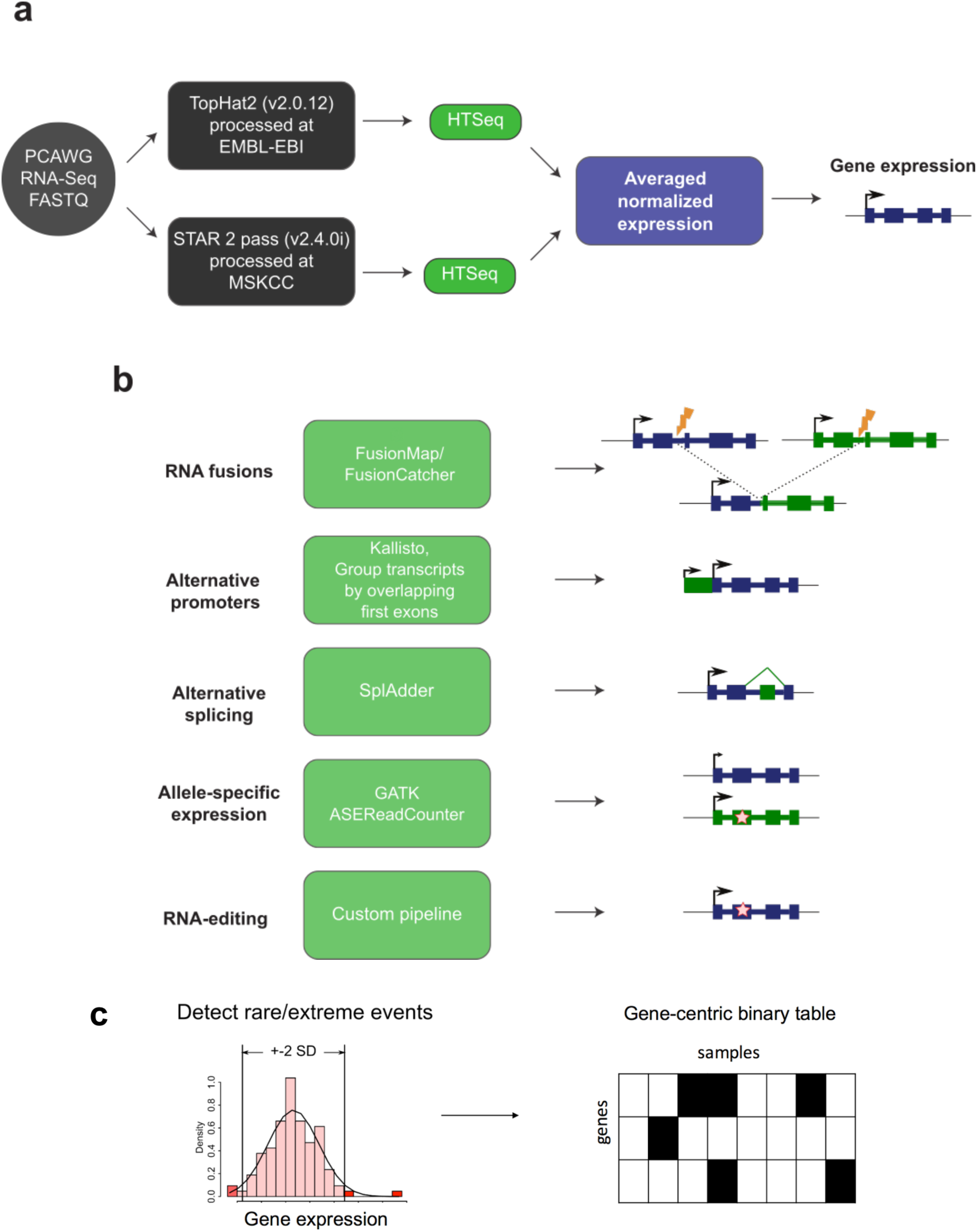
Unified RNA-Seq analysis to identify RNA-level alterations. **a**. Workflow of RNA-Seq alignment and quantification of gene expression. **b.** Computational methods used to detect additional types of RNA alterations including RNA fusions, alternative promoters, alternative splicing, allele-specific expression, and RNA editing. **c.** To unify analysis of alterations across all RNA phenotypes, a gene-centric binary table was created for each RNA phenotype indicating if a sample had an alteration in a given gene for a given sample. For quantitative RNA phenotypes (gene expression, alternative promoters, alternative polyadenylation, alternative splicing, and allele-specific expression), for a given gene, samples with extreme values, when compared to the samples in the same histotype, were considered altered.

**Extended Data Figure 2.**
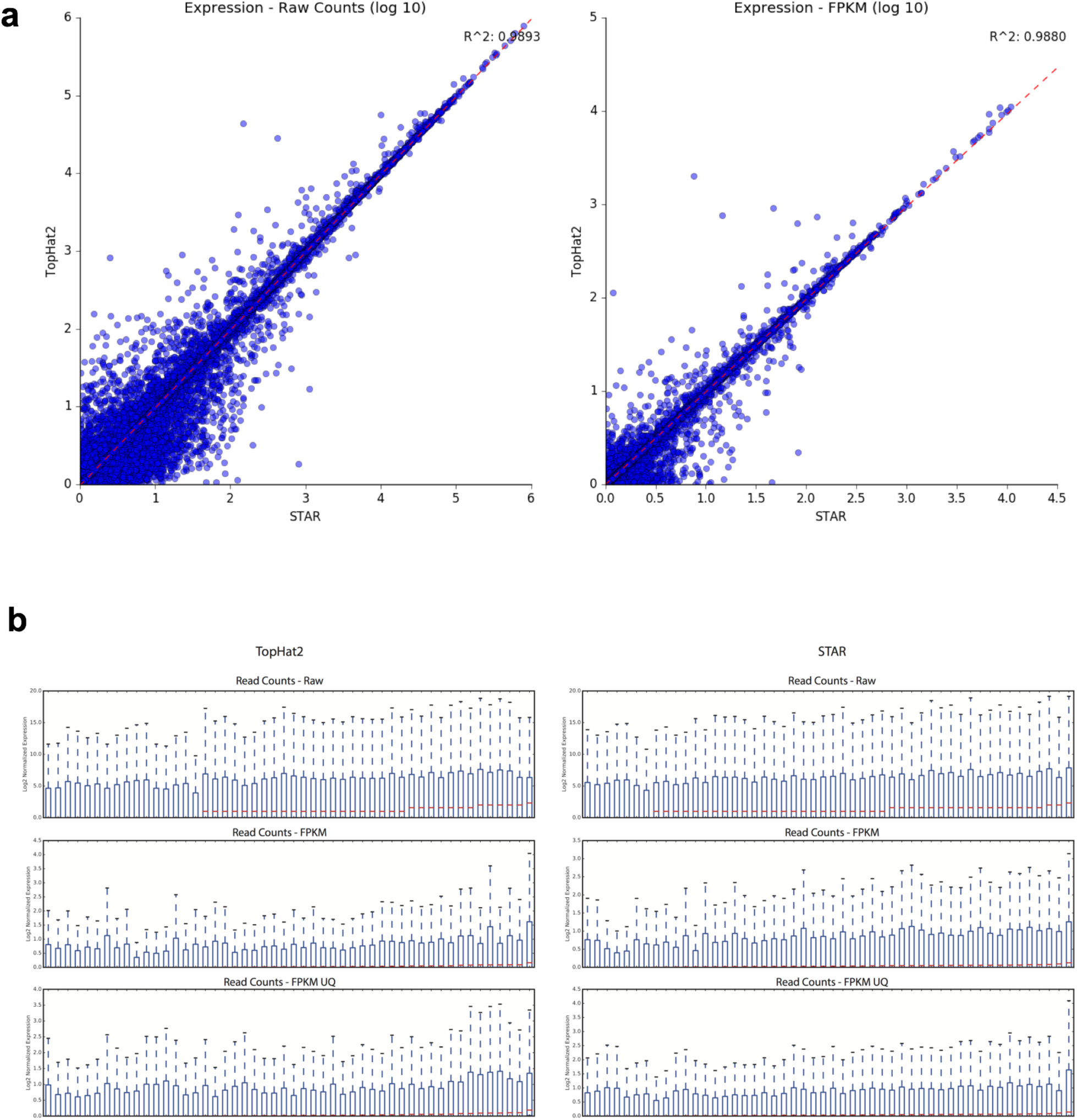
Consensus gene expression quantification and upper-quartile normalization. a. Correlation of STAR vs TopHat2 HTSeq counts for both raw counts (left) and FPKM-UQ normalized counts (right). b. Boxplots of raw (top), FPKM (middle) and upper-quartile normalized FPKM values (FPKM-UQ, bottom) for the same random subset of 50 samples taken from the cohort.

**Extended Data Figure 3.**
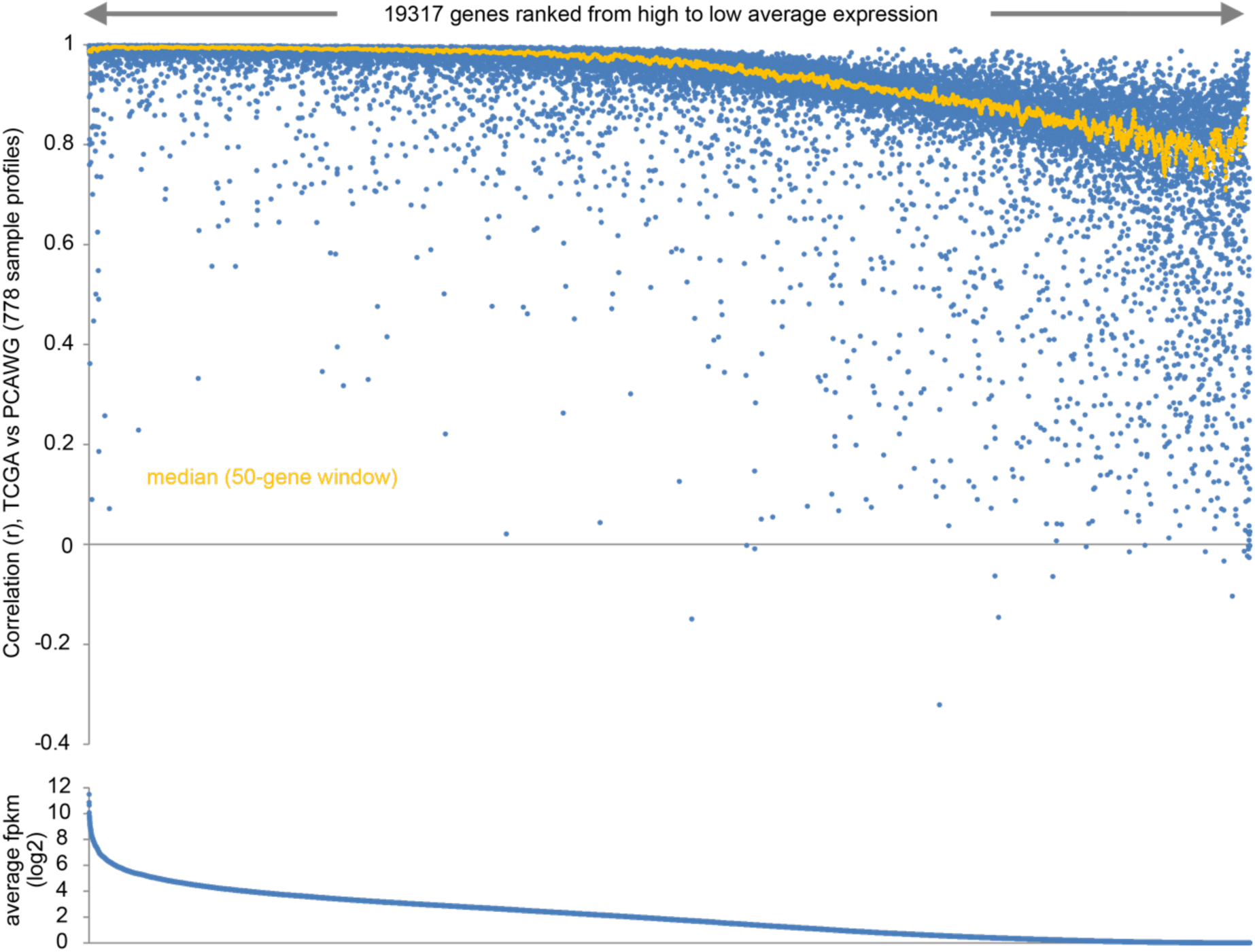
Comparison of TCGA RSEM and FPKM-UQ quantification. For 778 tumor expression profiles represented in both PCAWG and TCGA datasets, gene-level correlations (Pearson’s using log-transformed values) between the two datasets were computed (top scatter plot). Genes represented in both datasets are ranked from high to low average log2 fpkm values (PCAWG dataset).

**Extended Data Figure 4.**
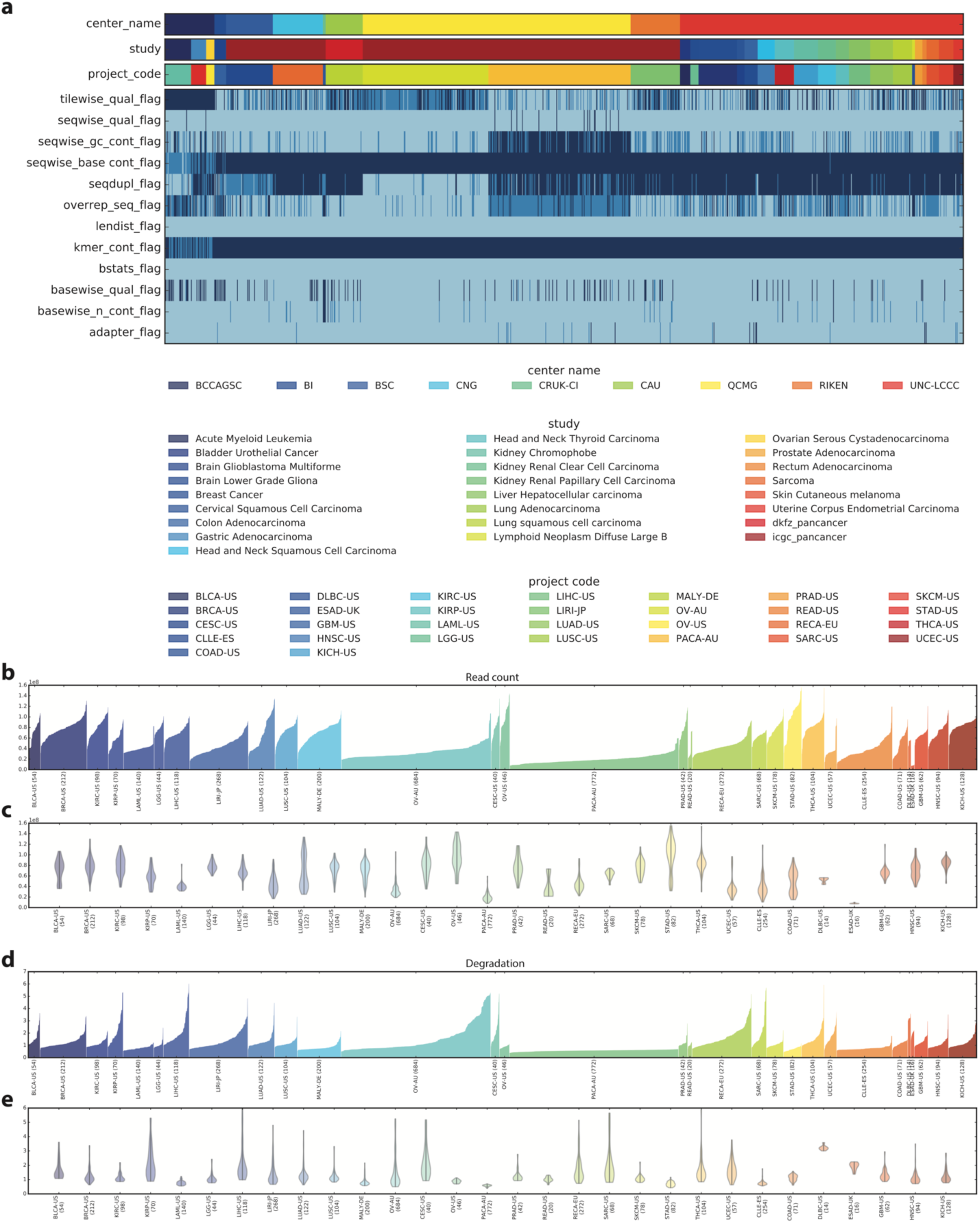
Quality control analysis of RNA-Seq data. **a.** Overview of QC measures collected on the full dataset based on output of the FastQC tool. Top bar encodes sequencing center of the library, middle top bard encodes the study metadata as used for tracking, lower top bar labels the project code of a library. **b.** Total read count per library shown as histogram. Libraries are colored by project code. **c.** Total read count per library shown as comparative distributions. Libraries are colored by project code. **d.** Sample degradation scores (3’->5’ bias) per library shown as histogram. Libraries are colored by project code. **e.**Sample degradation scores (3’->5’ bias) per library shown as comparative distributions. Libraries are colored by project code.

**Extended Data Figure 5.**
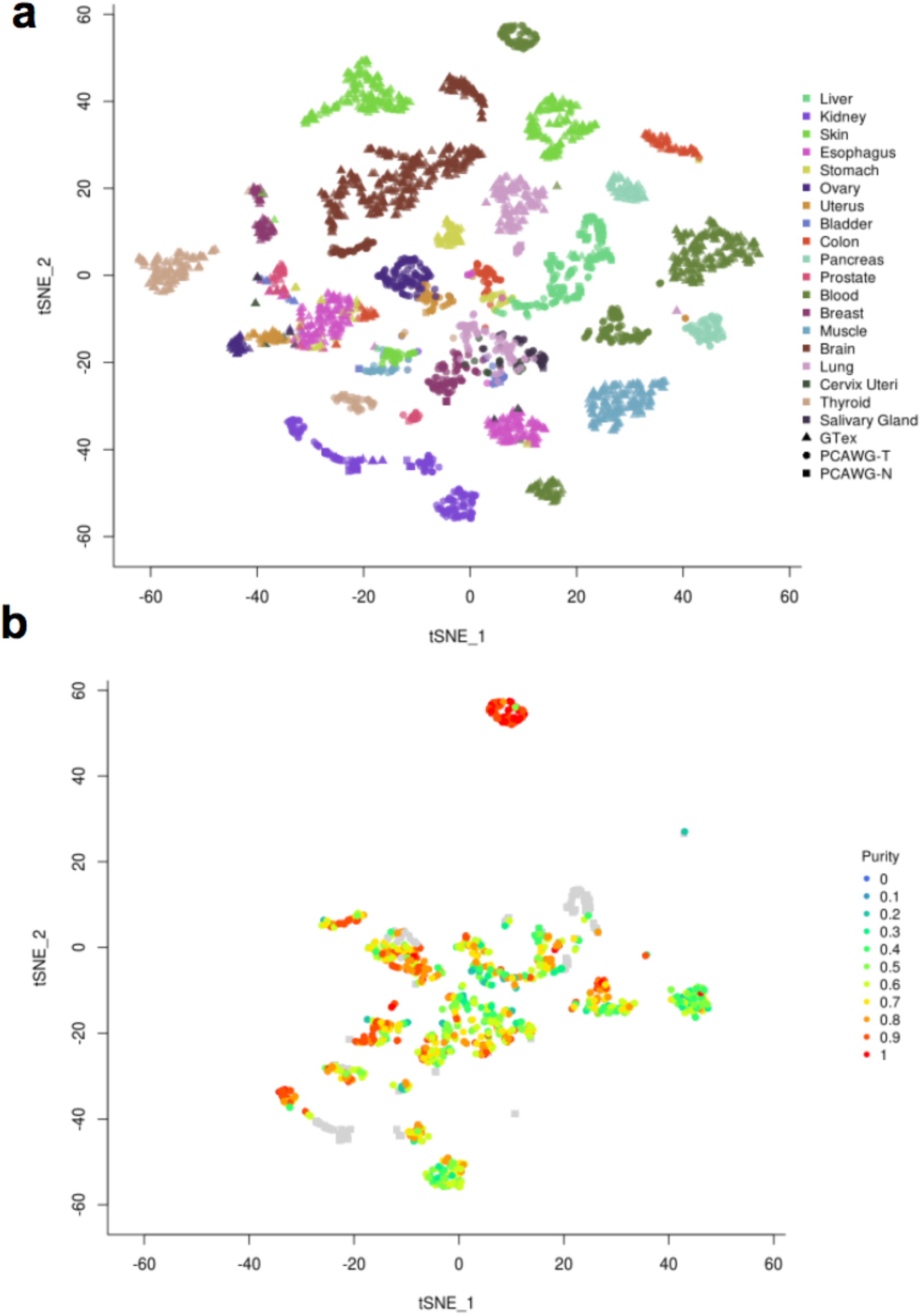
t-SNE analysis of gene expression. a) t-SNE plot based on gene expression from samples from GTEx (normal samples) and PCAWG (normal and tumor samples) coloured by tissue.b) t-SNE plot (same as in panel a) with the PCAWG samples coloured based on the estimated tumor purity.

**Extended Data Figure 6.**
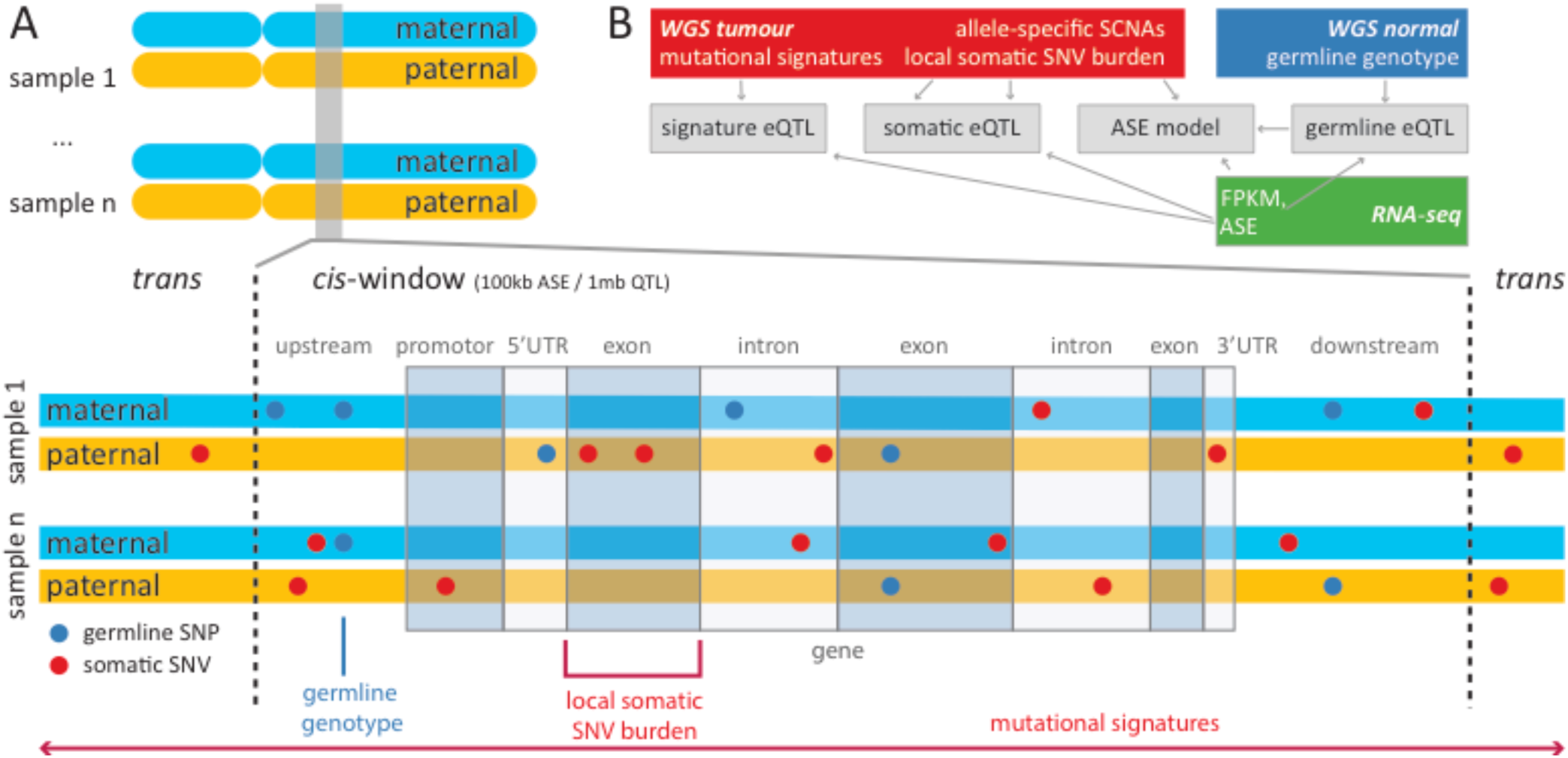
Integrative regulatory variant approach. Overview of the different sources of genetic variation considered in the analysis. Germline variants (blue) were individually tested for association with total gene expression using standard eQTL approaches. Cis somatic variants were aggregated in burden categories and tested for global association with allele-specific expression, as well as total expression on a per-gene level using eQTL analyses. Trans effects were estimated by testing total gene expression for association with mutational and epigenetic signatures. Window sizes were 1Mb for all cis eQTL analyses and 100kb for allele-specific expression.

**Extended Data Figure 7.**
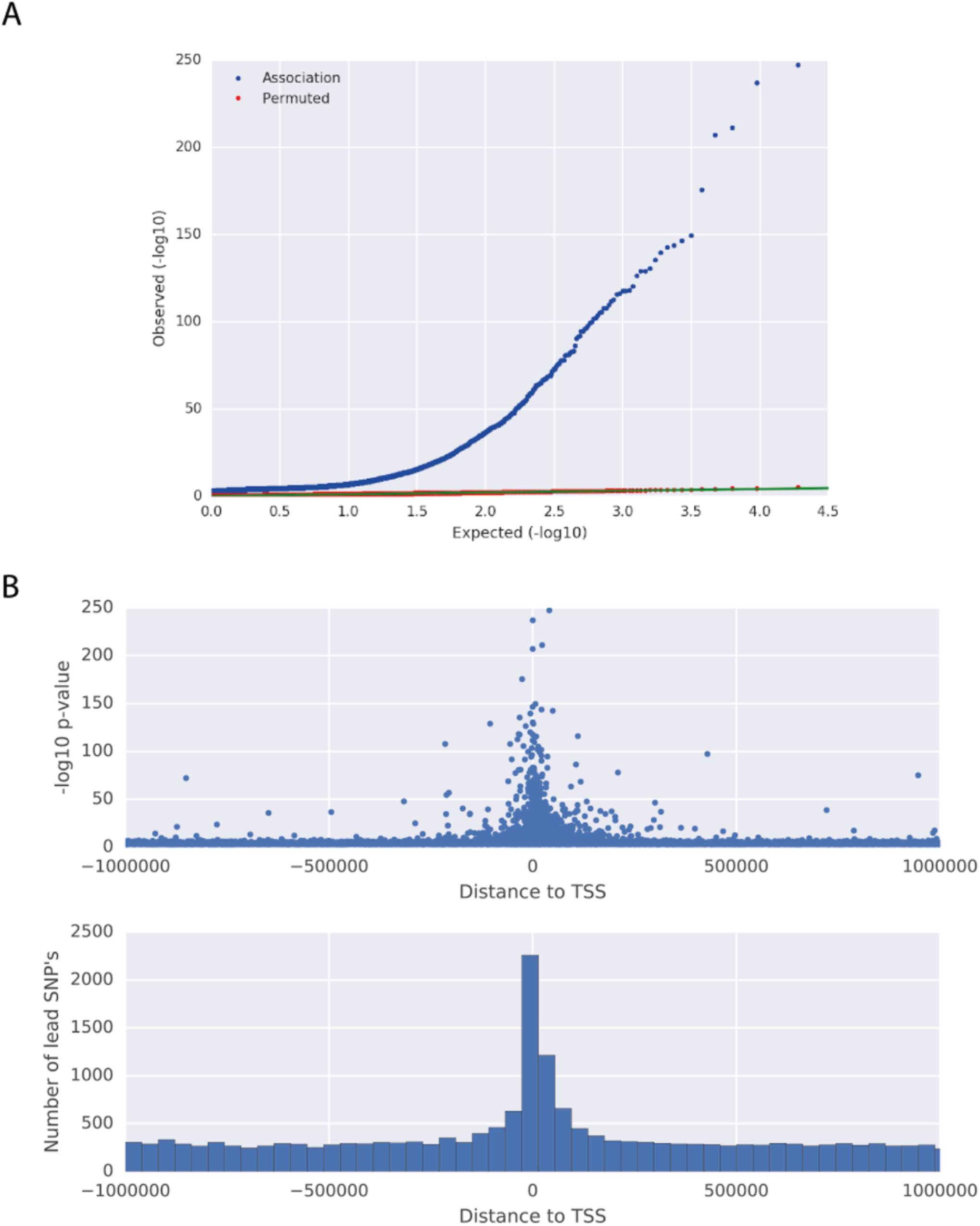
Germline eQTL lead variants. **A)** QQ plot of p-values of germline eQTL lead variants in the pan-cancer analysis (FDR ≤ 5%, blue) and p-values of the same analysis after permutation (random permutation of patients, red). **B)** Distribution of distance to the respective transcription start site of all germline eQTL lead variants.

**Extended Data Figure 8.**
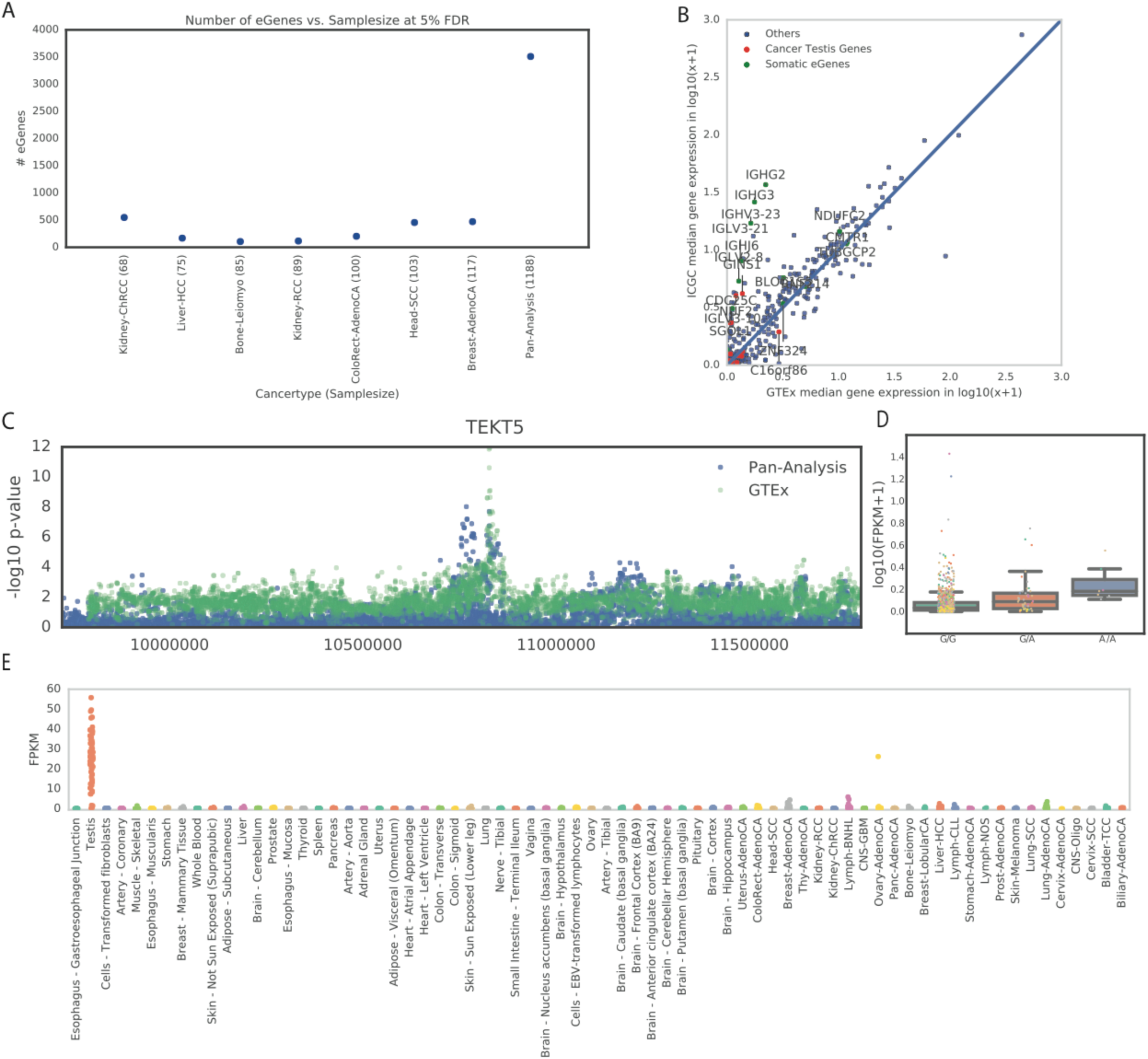
Germline eGenes. **A)** Number of germline eGenes (genes with at least one germline eQTL, FDR ≤ 5%) per cancer type, sorted by sample size (in parenthesis). **B)** Median gene expression across the ICGC and GTEx cohort for 411 ICGC-specific eQTL (green: somatic eGenes, red: cancer testis genes)**. C**) Manhattan plot for *TEKT5*, showing associations in the ICGC cohort (blue) and in GTEx (green, minimum p-values across all GTEx tissues). Two independent eQTL were identified. **D**) Boxplot of log_10_FPKM of the ICGC-specific eQTL lead variant of *TEKT5*. **E**) *TEKT5* gene expression (FPKM) in GTEx tissues and ICGC cancer types.

**Extended Data Figure 9.**
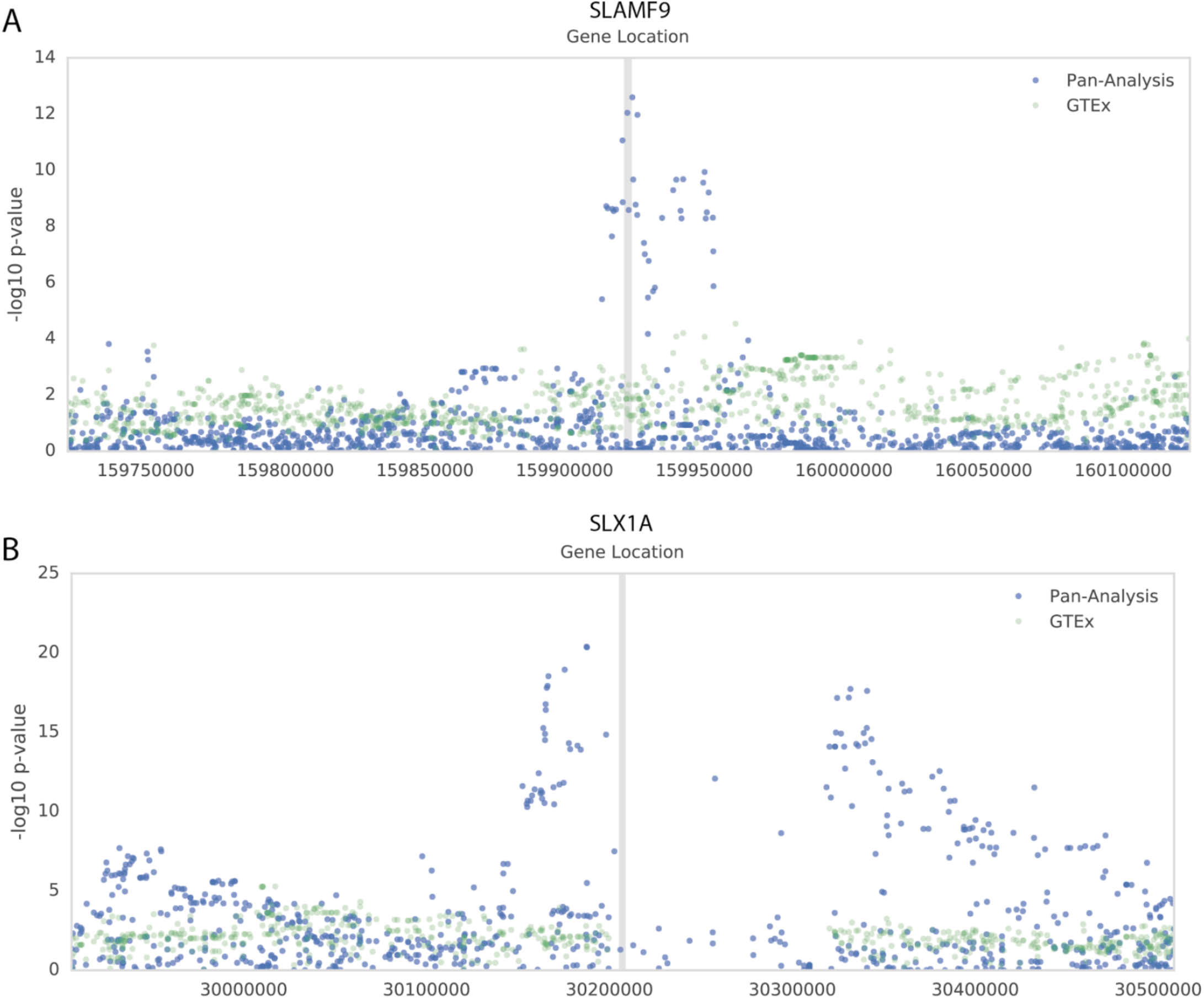
Mahattan plot GTEx comparison. Manhattan plots showing associations of the pan-analysis in the ICGC cohort (blue) and in GTEx (green). **A)** Manhattan plot of *SLAMF9* as an example of an pan-analysis specific eQTL. **B)** Manhattan plot of *SLX1A* as another example of an pan-analysis specific eQTL.

**Extended Data Figure 10.**
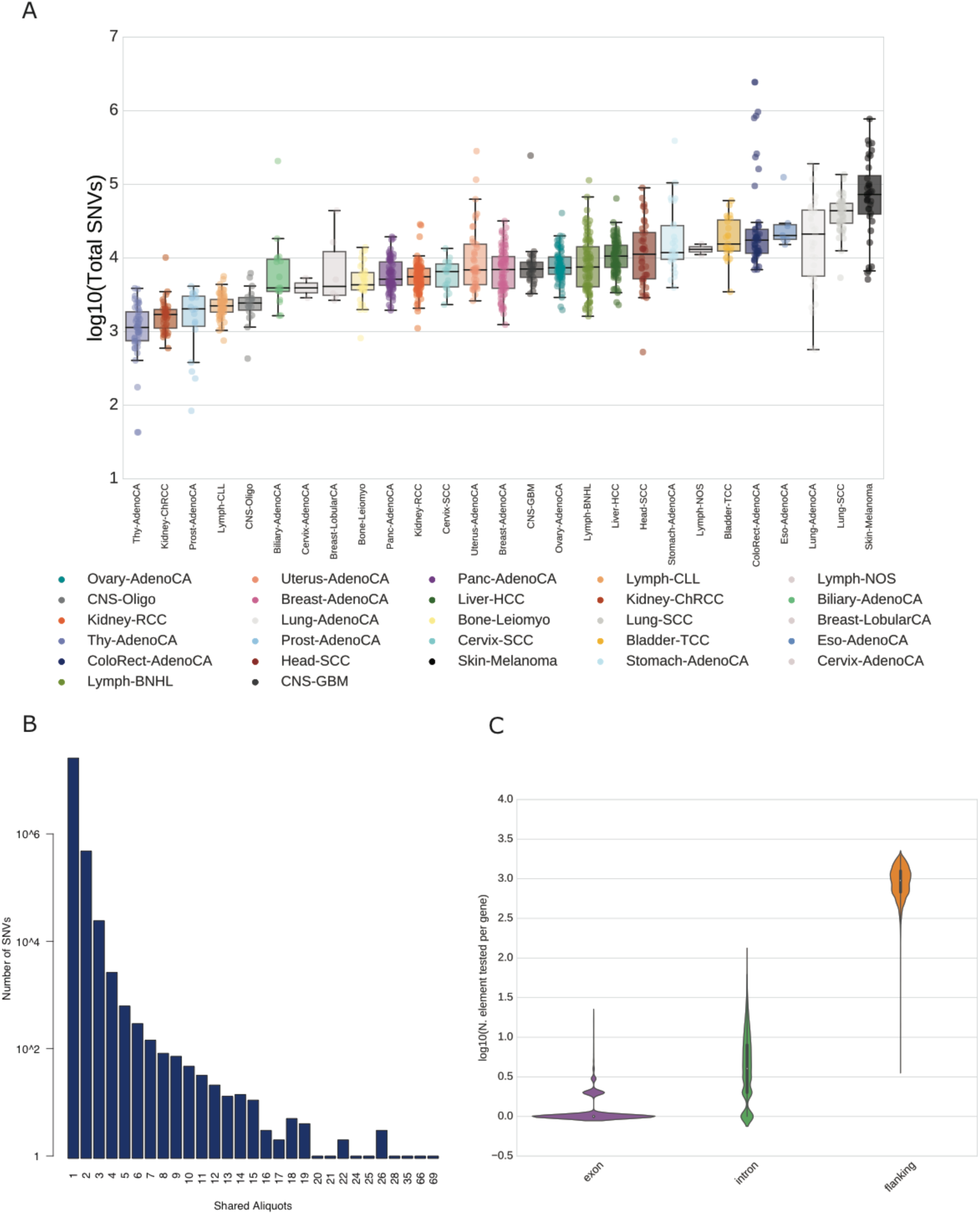
*Cis* mutational somatic burden. A) Total somatic mutational load per cancer type. Median numbers of SNVs ranges from 1,139 in thyroid adenocarcinoma to 72,804 in skin melanoma. B) Number of somatic SNVs shared by patients. A small fraction of 86 SNVs is shared by more than 1% of the cohort (12 patients). C) Number of mutated regions with somatic burden frequency ≥1% tested per gene, stratified according to their genomic position.

**Extended Data Figure 11.**
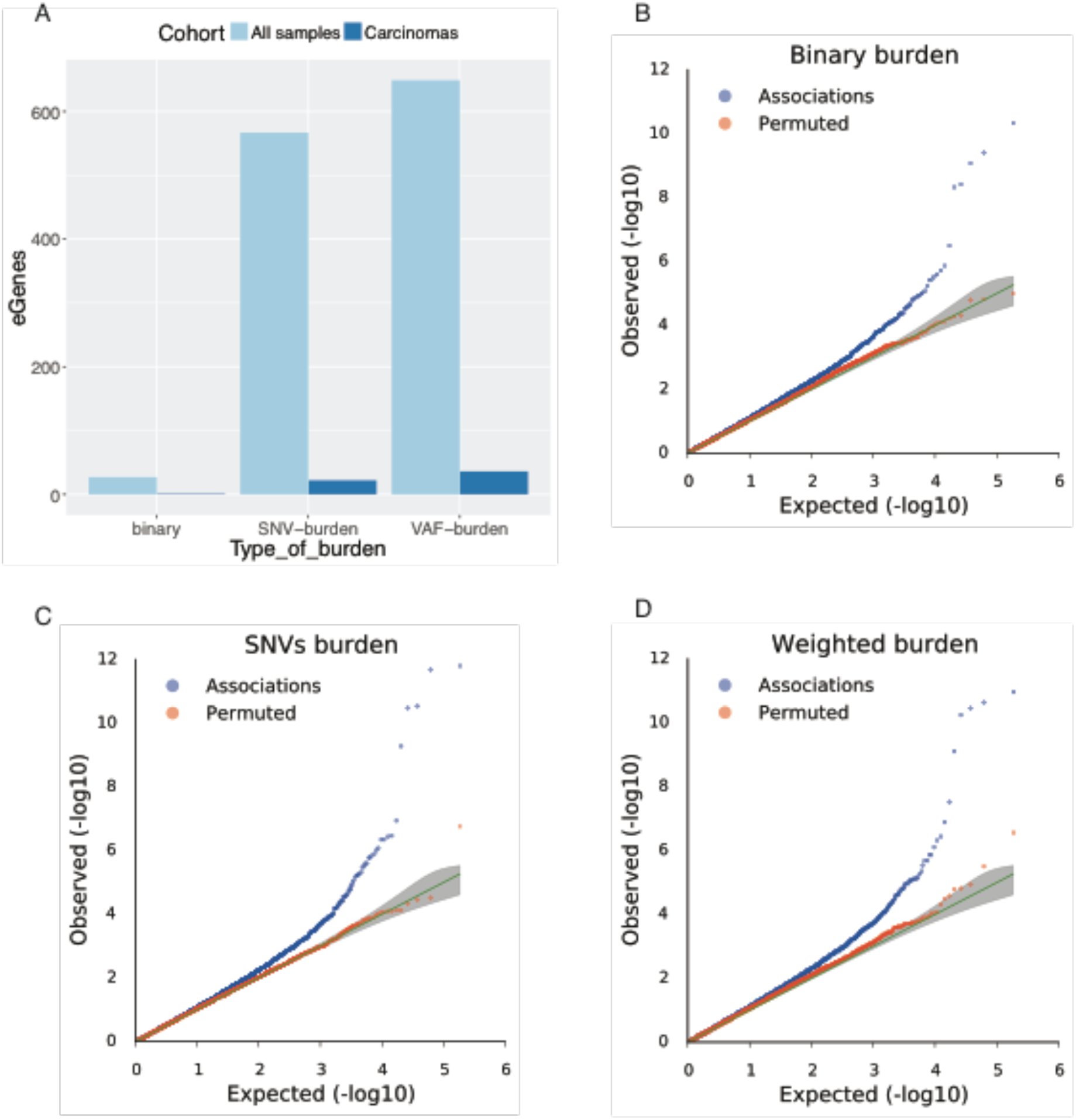
Power of different strategies for estimating somatic mutational burden for eQTL analysis. A) Number of significant somatic eQTL (FDR ≤ 5%) identified with different mutational burden estimates using all 1,188 patients and a subset of 899 carcinomas patients. B-D) QQ plots of the p-values of the somatic eQTL analysis. Considered were mutational burden calculated as B) binary burden (presence or absence of at least one somatic mutation), C) total SNVs load (number of somatic mutations per element) or D) weighted burden (sum of variant allele frequencies over the genomic region tested, Methods).

**Extended Data Figure 12.**
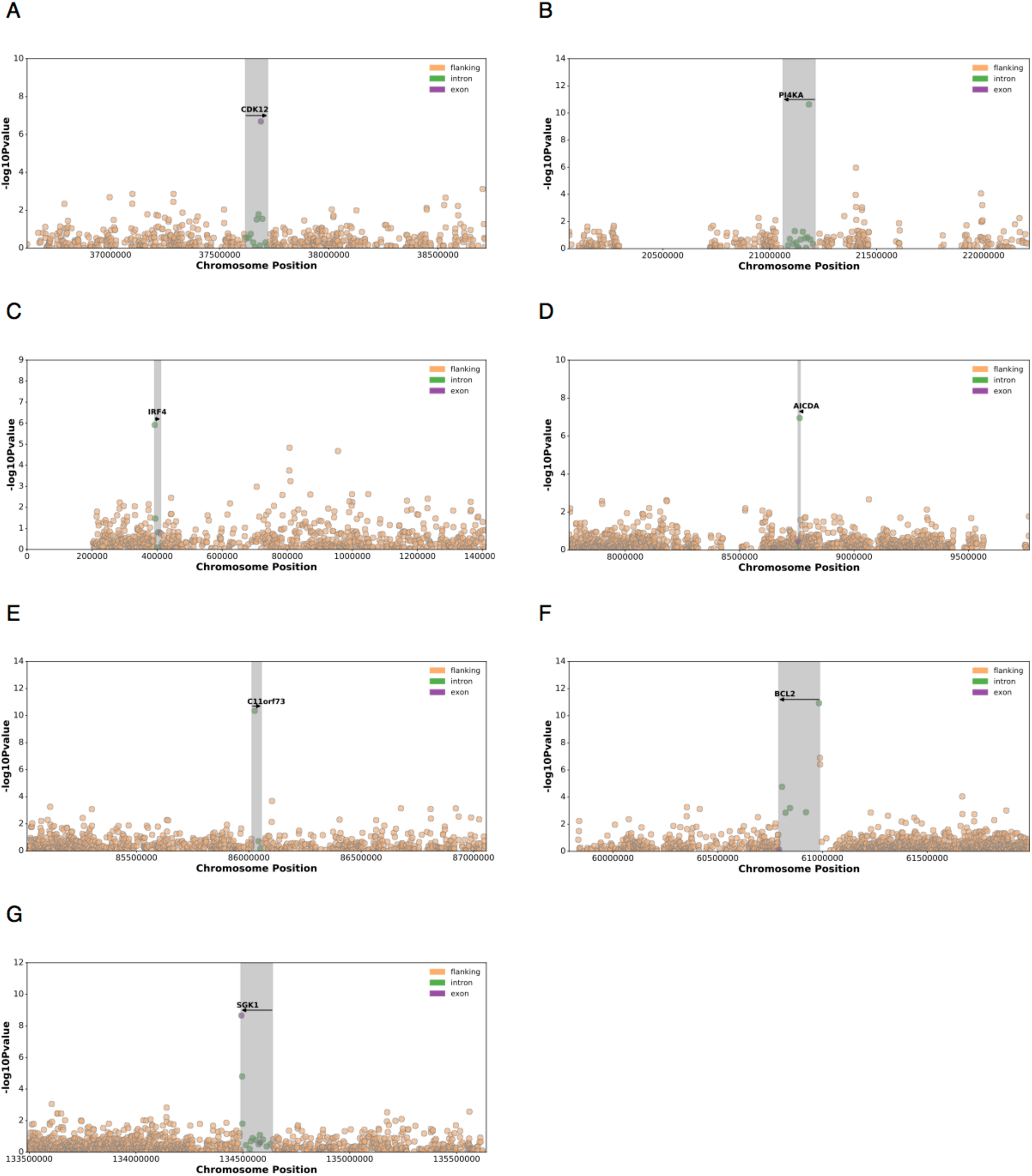
Manhattan plots of seven somatic eGenes associated to genic lead burden. Altogether, eleven genic somatic eQTL were detected to show significant gene expression changes associated to somatic burdens within the gene boundaries (intronic or exonic). The seven genes shown here are known to be important in the pathogenesis of specific cancers. **A)** *CDK12*. **B)** *PI4KA.* **C)** *IRF4*. **D)** *AICDA*. **E)** *C11orf73.* **F)** *BCL2*. **G)** *SGK1*.

**Extended Data Figure 13.**
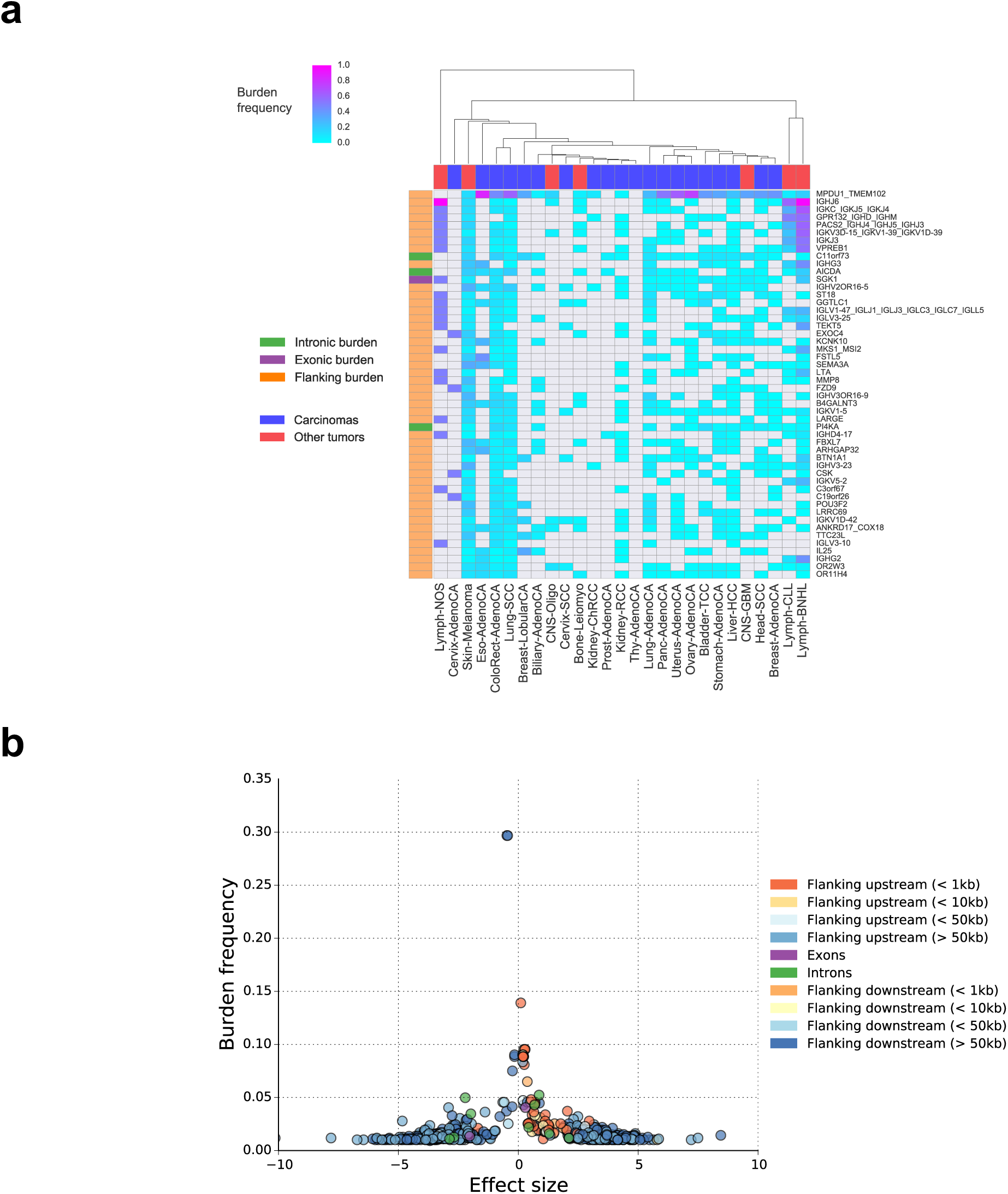
Somatic burden prevalence in the cohort. **a.**Clustering of somatic *cis* eQTL (FDR ≤ 5%) by mean burden frequency estimated in each cancer type. The heatmap shows the first top 50 associations, sorted by mean burden frequency of the lead element across all cancer types. Row labels describe the HGCN names of the eGenes associated to leading somatic burden. Multiple eGenes associated to the same genomic interval are joined by an underscore. Row colors indicate the genomic region of the burden (flanking, intronic or exonic). Column colors distinguish the two main tumor types in the cohort, namely carcinomas and other tumors (lymphomas, skin melanoma and glial tumors). **b.** Burden frequency versus effect size for somatic associations.

**Extended Data Figure 14.**
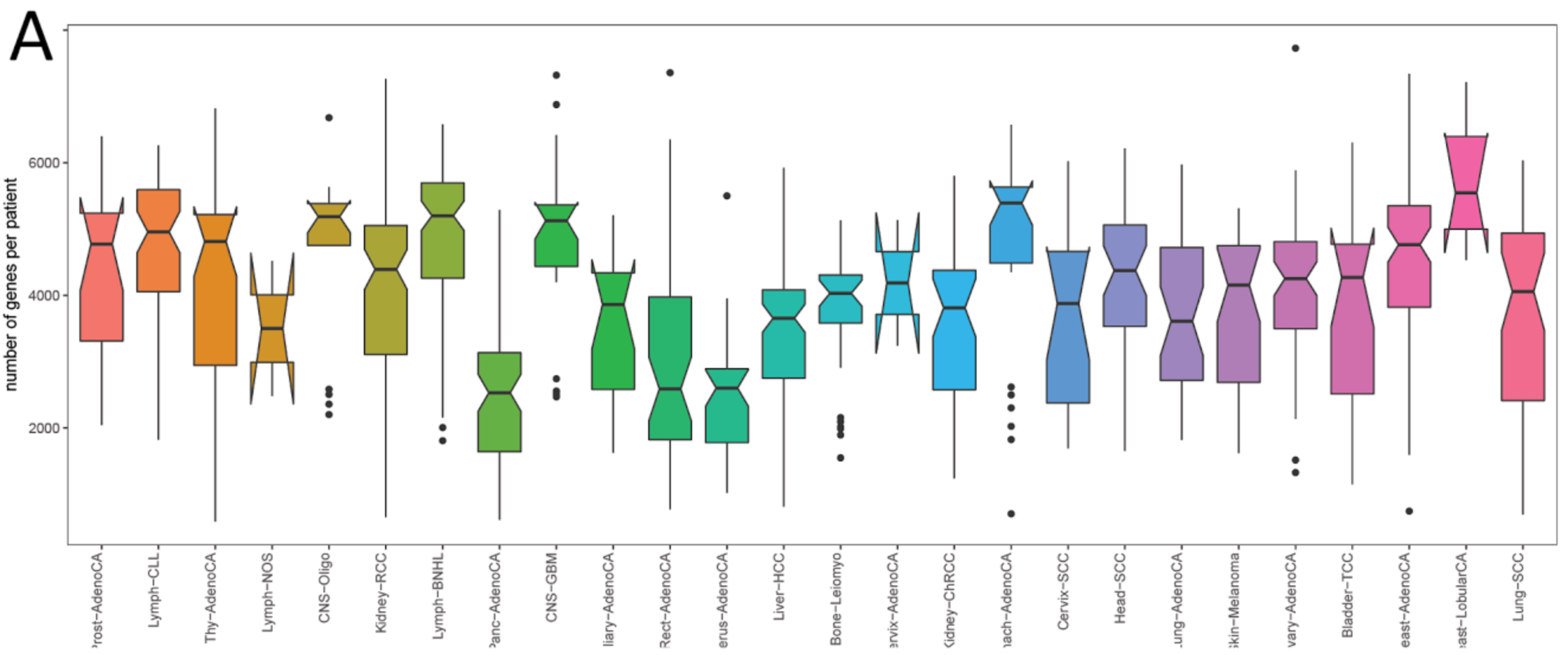
ASE analysis. All cancer types are ordered by average AEI frequency. Number of genes per patient for which ASE could be quantified, stratified according to cancer type, resulting in between 588 and 7,728 genes per patient.

**Extended Data Figure 15.**
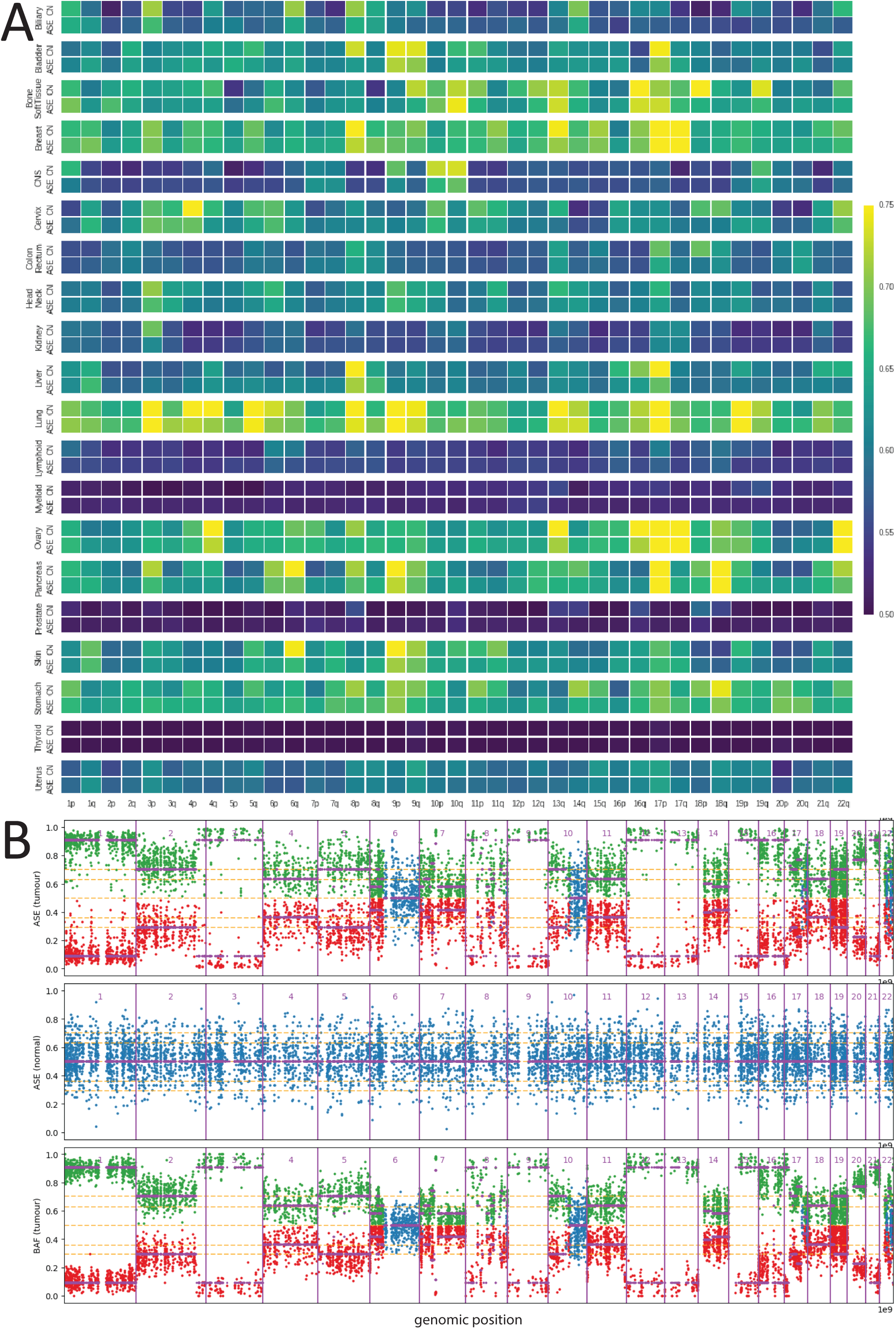
SCNAs as major driver for allelic dysregulation in cancer. A) Absolute allelic expression imbalance closely follows allelic imbalance on the genomic level. Values of 0.5 (blue) denote equal number of reads from both alleles. Values of 1 (yellow) reflect monoallelic expression or regions with loss of heterozygosity. B) Comparison between B-allele frequency (BAF) and ASE ratios from a single lung cancer patient (LUAD-US) with profound chromosomal instability shows strong correlation between allelic imbalance on expression and genomic levels.

**Extended Data Figure 16.**
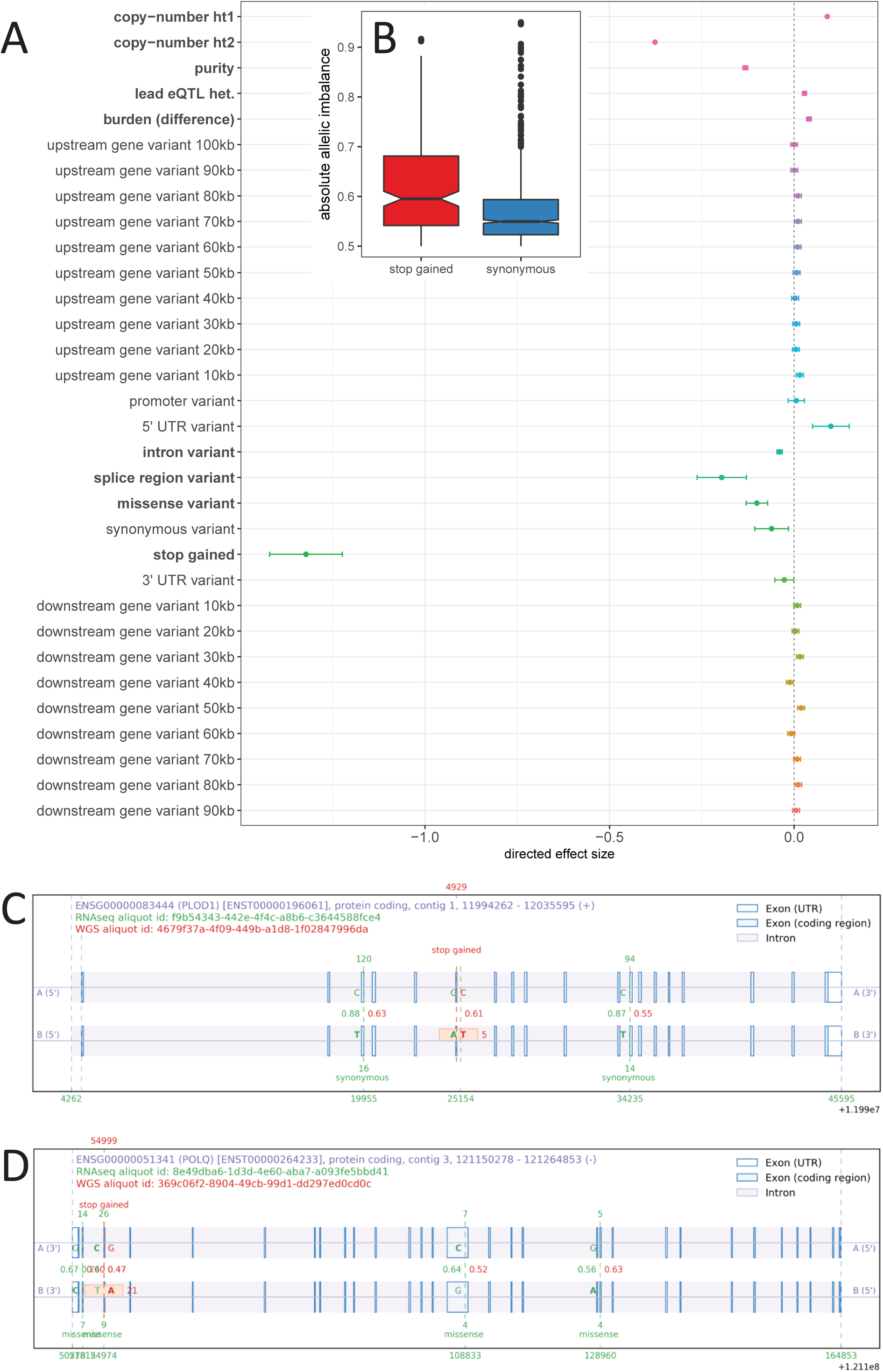
Results of a directed joint multivariate model that links individual somatic events to the ASE ratio. **A)** Relevance of individual somatic mutation types (‘copy-number ht1’ and ‘copy-number ht2’ as local allele-specific SCNAs of haplotypes 1 and 2), germline eQTL and other co-variates for ASE ratio. Significant covariates (FDR ≤ 5%) highlighted in bold. **B)** Comparison of the effect of protein truncating variants (‘stop-gained’) and synonymous variants on ASE ratio. **C6D)** Examples of ASE ratio dysregulation.

**Extended Data Figure 17.**
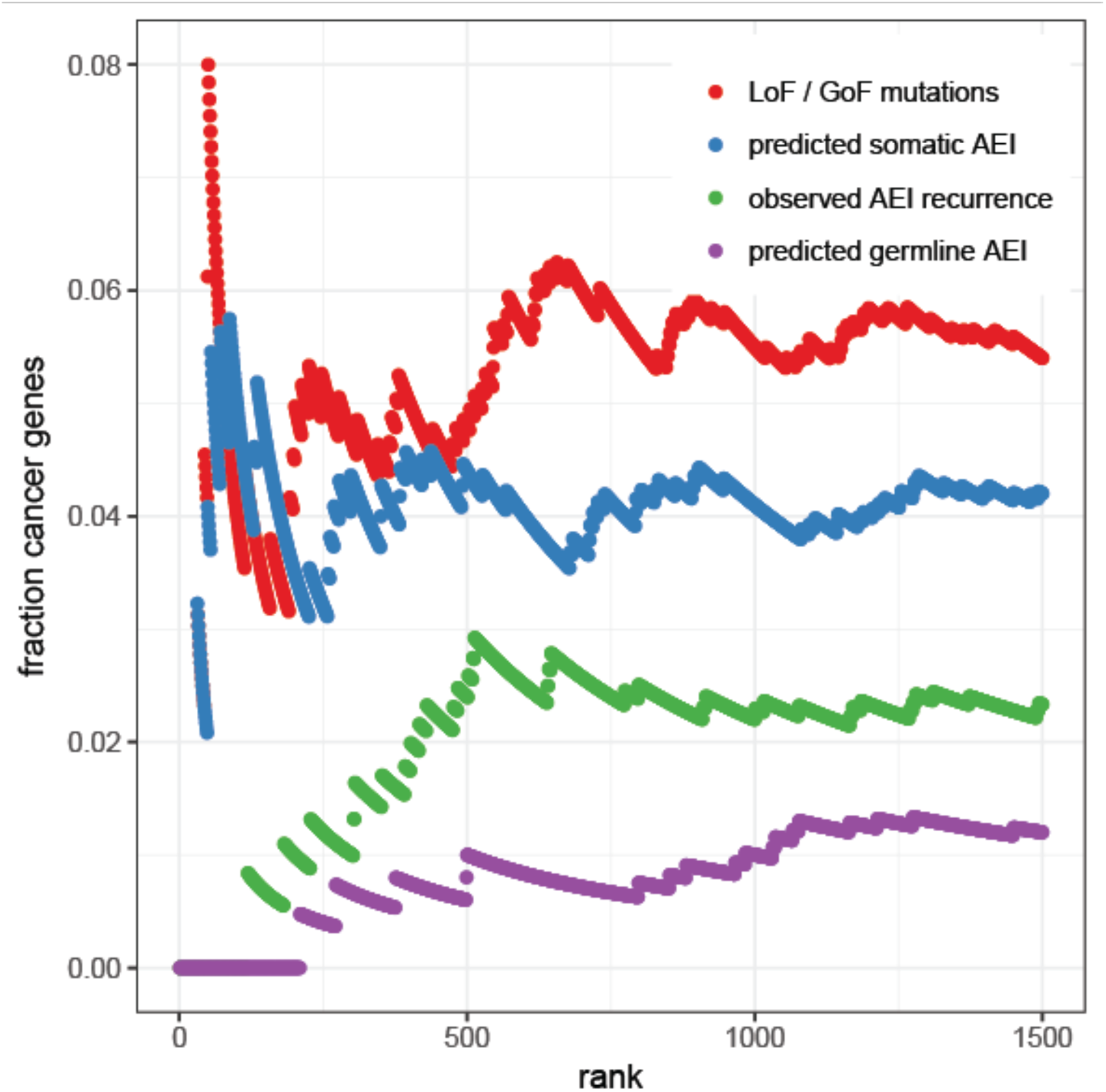
Somatic allelic imbalance predicts cancer relevant genes.

**Extended Data Figure 18.**
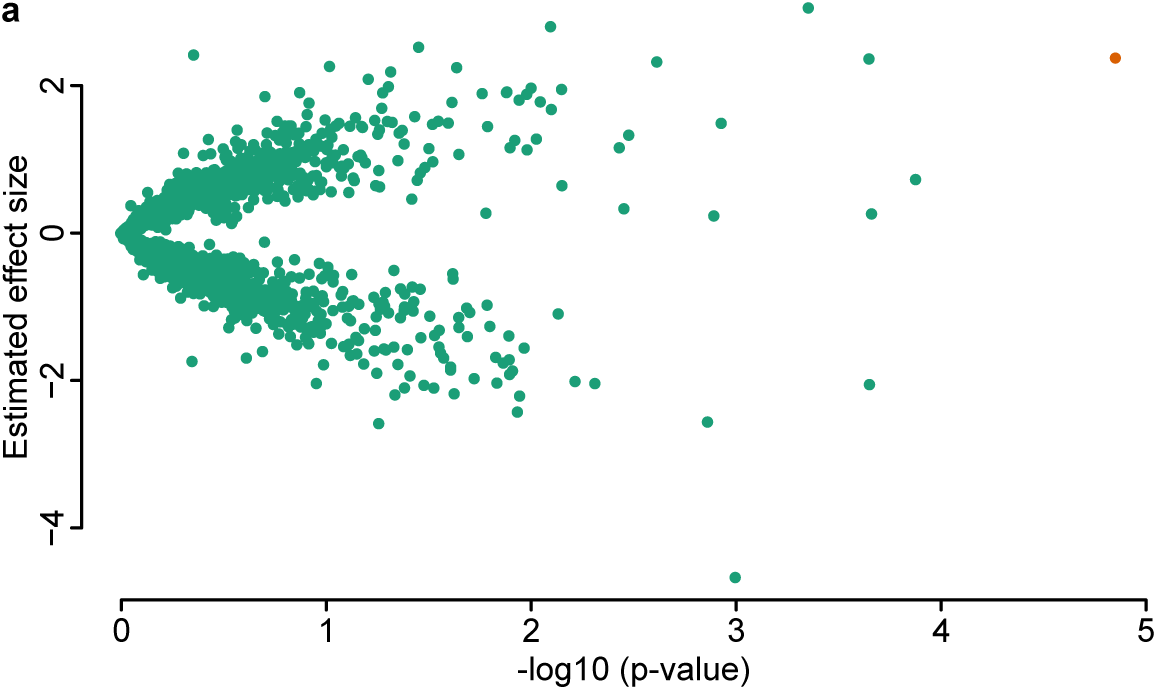
Association of promoter activity with promoter mutations across all samples. The orange colour denotes the significant associations.

**Extended Data Figure 19.**
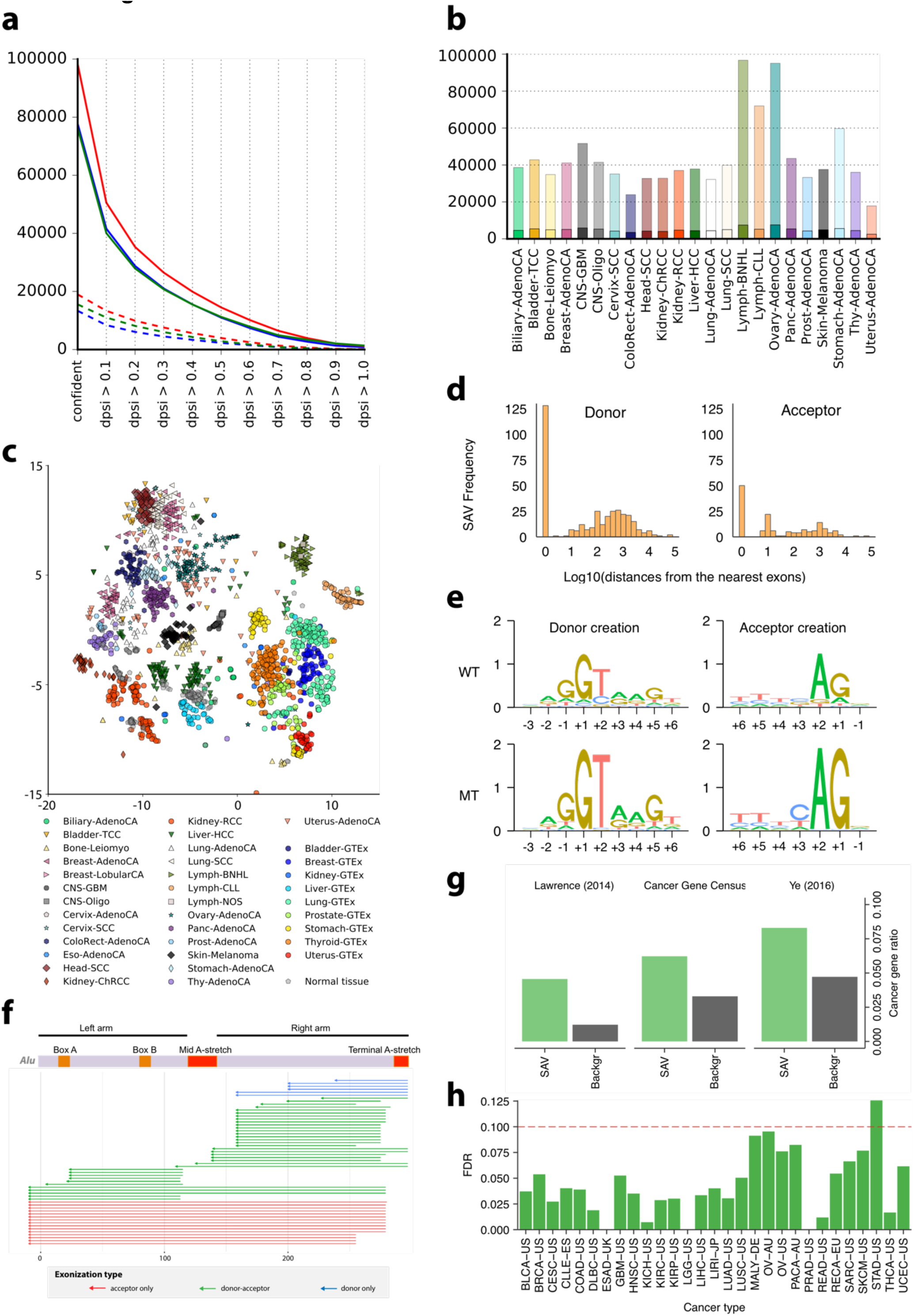
Alternative splicing and association with somatic mutations. **a)** Number of exon skipping events confirmed at different delta PSI thresholds in tumor (red), normal (green) and GTEx (blue) samples for Liver tissue. Dashed lines show the subset of exon skippings that only contain annotated introns. **b)** Number of exon skipping events confirmed at a delta PSI level greater than 0.3 for the individual histotypes. Transparent section of bars represents fraction of novel events, containing at least one un-annotated intron. **c)** Splicing landscape for exon skipping events. t-SNE embedding based on exon skip PSI values for all ICGC tumor and normal samples together with tissue matched GTEx samples. **d)** Permutation-based FDR values for SAV detection based on the different cancer types. **e)** Cancer gene set enrichment for SAV sets, shown for cancer census gene set (middle) and sets determined by Lawrence et al. (left) and Ye et al. (right). **f)** Positional distributions (logarithms of distance from the nearest exons) of somatic variant creating novel splicing donors and acceptors. **g)** Sequence motif logos around somatic mutation creating novel splicing motifs. **h)** SAVs aligned to overlapping ALUs aligned to the ALU reference, showing exonizations creating a novel acceptor (red), a novel donor (green) or both (blue).

**Extended Data Figure 20.**
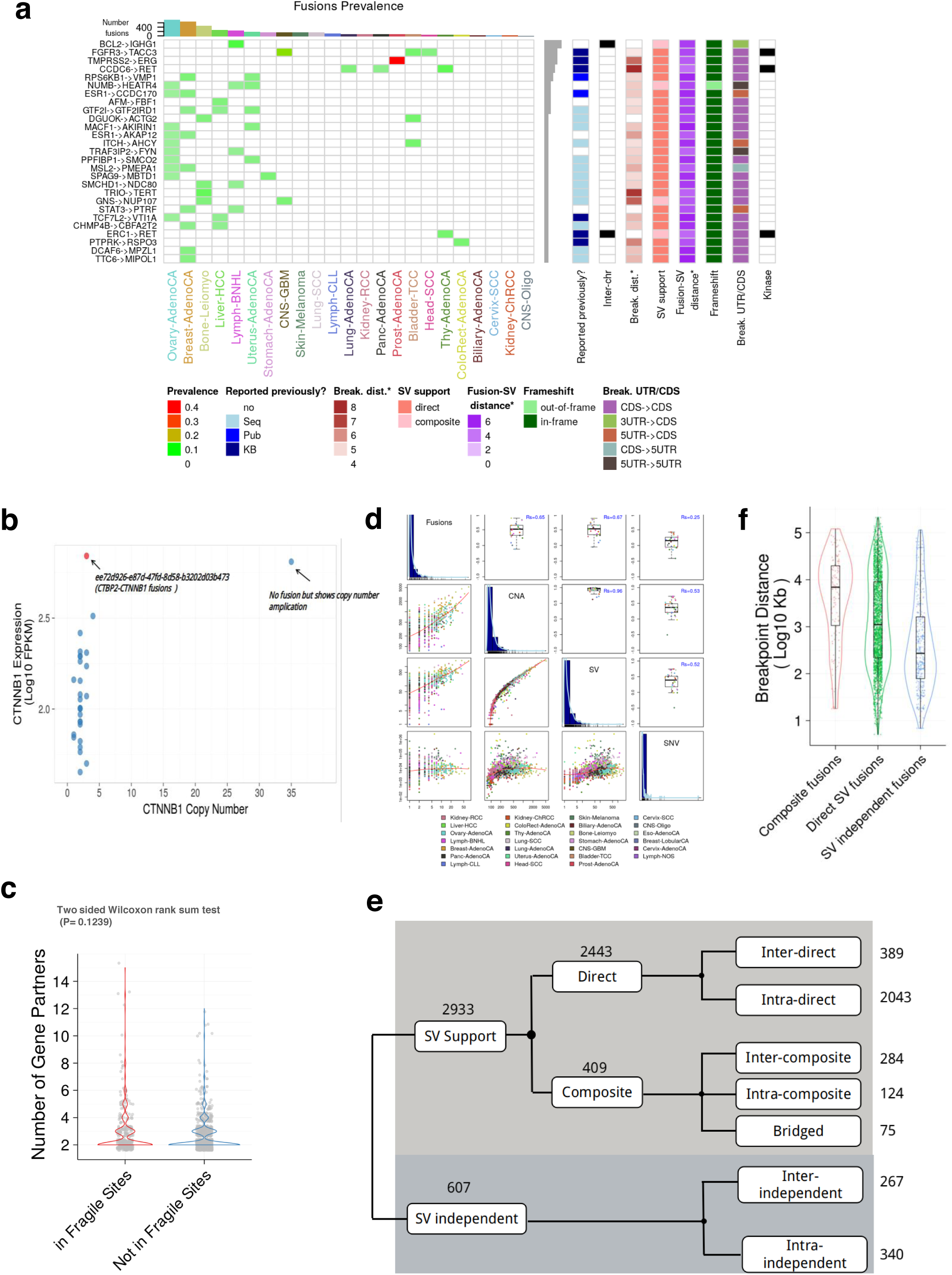
Structural rearrangements associated with RNA fusions. **A).**Features of the 27 most recurrent in-frame or ORF-retaining fusions. Kinase column indicates whether one of the gene partners is a kinase gene **B)**.*CTBP2@CTNNB1* as an example of “Retained ORF” fusion. A scatter plot of *CTNNB1* DNA copy number versus mRNA expression across all ICGC gastric cancer samples **C)**.Fusion genes with promiscuous gene partners overlapped with human common fragile sites do not show different number of gene partners. **D)**.Number of gene fusions per sample and respective number of fusions, structural variants (SV), copy number alterations (CNA), and single nucleotide variants (SNV). The diagonal histograms shows the distribution of the number of alterations per sample. The upper triangle presents the Spearman correlation between two types of alterations per histological type (dot) and together with the overall spearman correlation (in blue). The bottom triangle contains scatter plots contrasting the number of alterations for each sample (dot) **E)**.Systematic classification scheme of all gene fusions based on underlying SVs. Numbers of fusion events of different classes are shown to the right. **F)**. The distribution of distances of fusion breakpoints among intrachromosomal composite, direct SV-support and SV-independent fusions

**Extended Data Figure 21.**
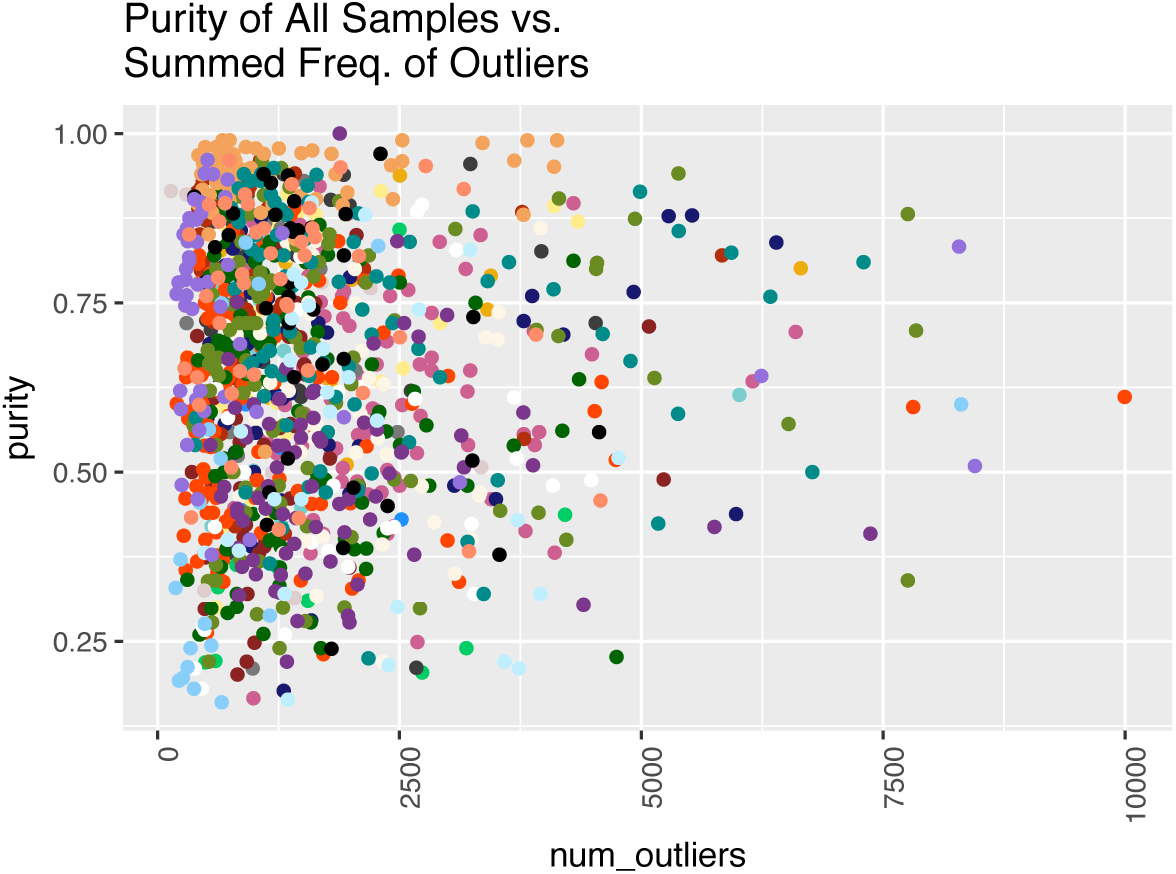
Correlation of Purity and Alteration Frequencies. When estimating the relationship between purity and frequency of outliers, and using histotype as a confounding variable, we find that there is no significant correlation (t-test P=0.36).

**Extended Data Figure 22.**
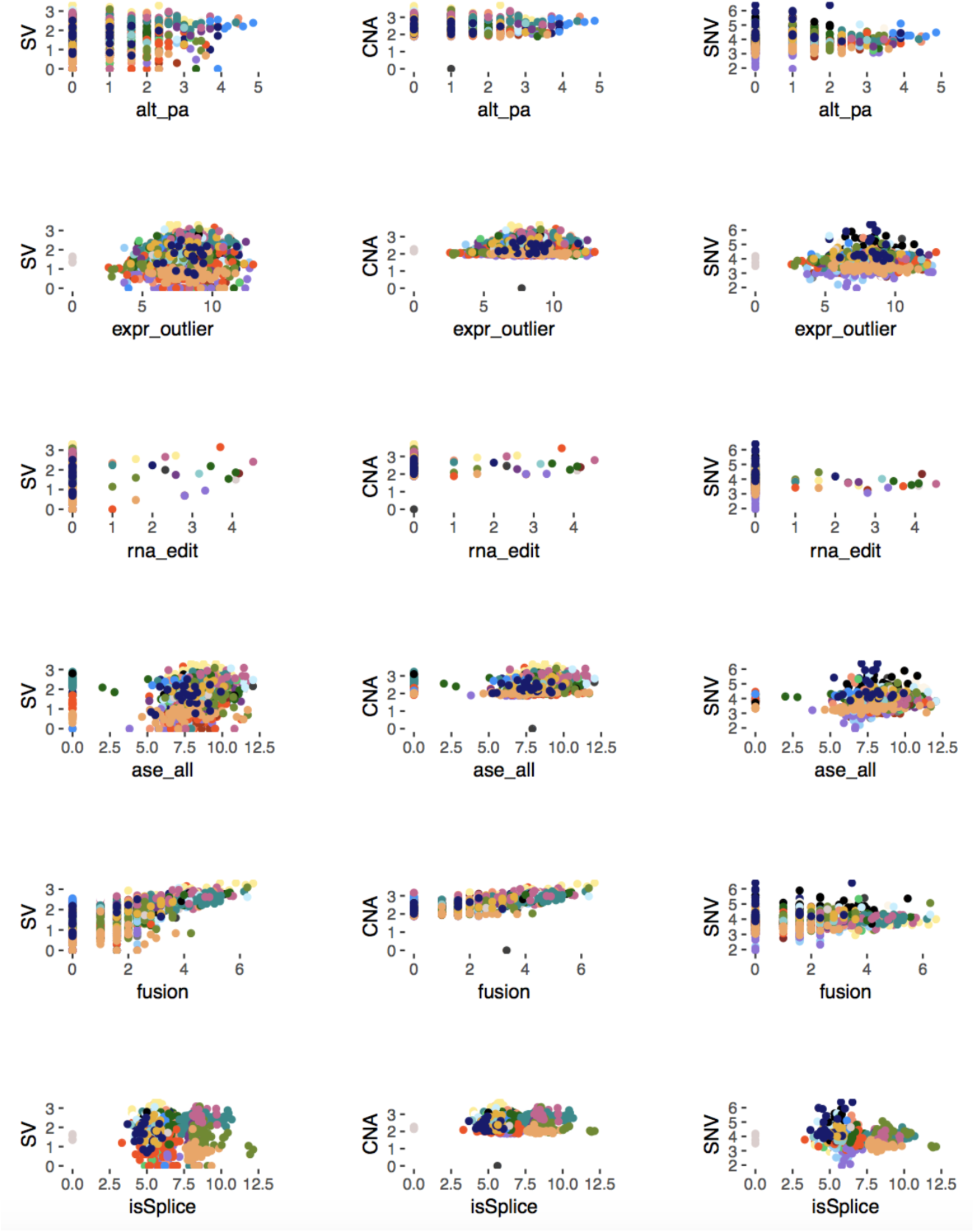
Correlation of the number of somatic genomic alterations with RNA alterations. Shown here are scatter plots of DNA alterations versus RNA alterations, each row is a DNA alteration in the following order: structural variants, copy-number aberrations, and non-synonymous variants. Each row is a RNA alteration in the following order: alternative polyadenylation, expression outliers, rnaediting, allele specific expression, fusions, and splicing. Each point is a sample colored by histotype, and its position is the log(number of aberrations) found in each sample.

**Extended Data Figure 23.**
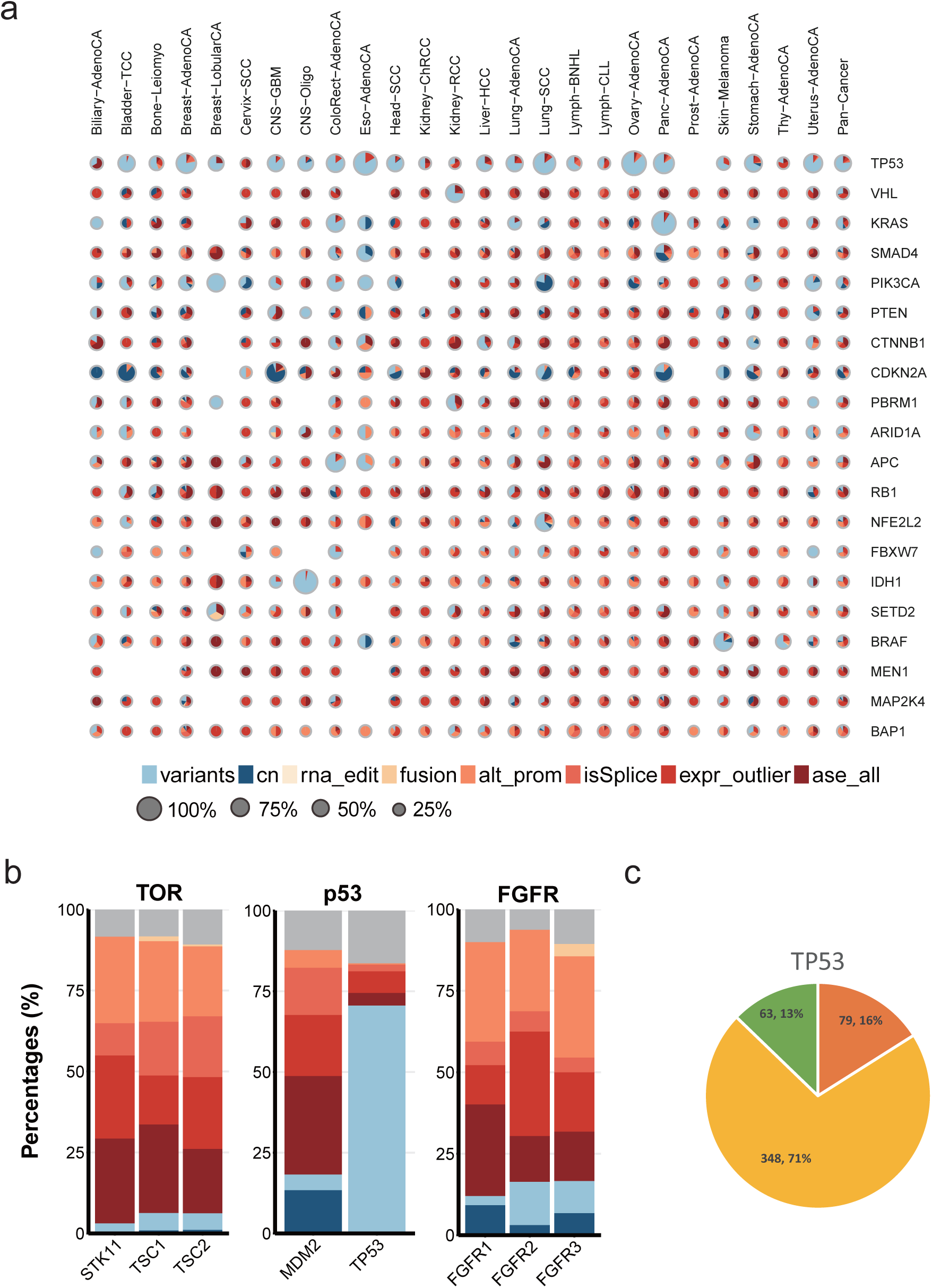
Breakdown of DNA and RNA alterations of cancer genes. **A.** Composite pie charts showing percentages of DNA and RNA alterations for top cancer driver genes. The 20 most significant cancer driver genes identified by PCAWG group in pan-cancer level are depicted, with sizes of pie charts indicating the percentages of patients carrying alterations in the given driver gene. The areas represent the relative percentages of patients exhibiting different alterations depicted by corresponding colors. When multiple types of alterations in one pathway affect the same patient, only a fraction is counted towards each type of the alterations. **B.** Proportional bar plots showing the distribution of gene alterations for genes in the TGF-Beta, apoptosis and FGFR pathways. **C.** Pie chart showing the breakdown of alteration types impacting TP53.

**Extended Data Figure 24.**
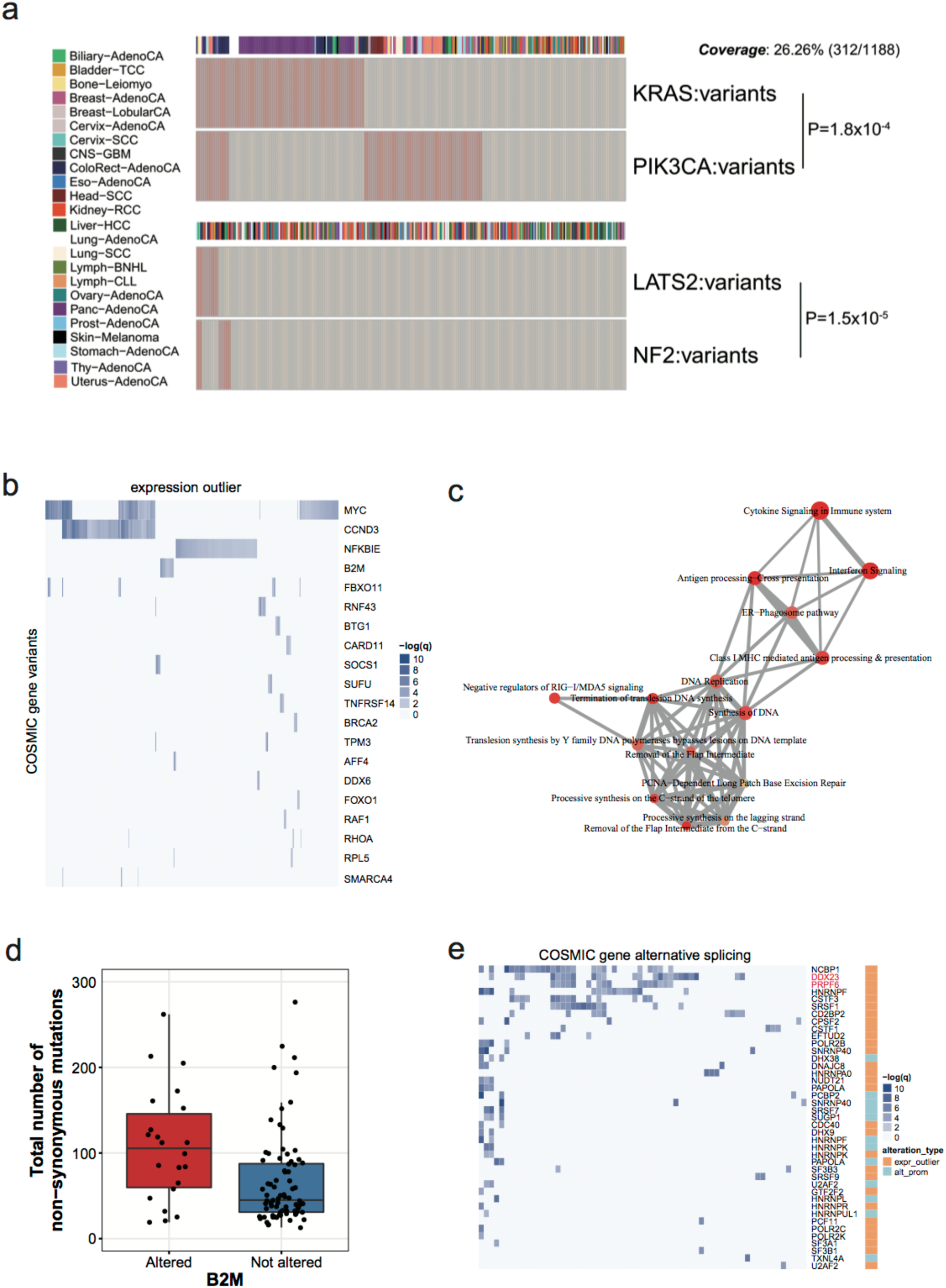
Trans-associations found by co-occurrence analyses. **A.** Heatmap showing the known co-occurrence between mutations of KRAS and PIK3CA, and those between LATS2 and NF2. Each column indicates a specific tumor with tumor types annotated to the left. Most samples without the listed alterations are not shown for space considerations. **B**. Heatmap showing the extent of associations between COSMIC gene somatic mutations and expression outliers of all genes. Each column indicates one gene, and the color intensity shows the significance of trans-association. COSMIC genes labeled to the right are ordered by the number of significant associations. Only the top 20 genes are shown. **C**. Enrichment map showing the significant (FDR ≤ 0.01) pathways based on the top 100 significant genes associated with B2M alterations.Color intensity represents enrichment significance, node sizes the number of analysed genes belonging to the given pathway, and edge sizes the degree of overlap between two gene-sets. **D**. Boxplots showing the total number of non-synonymous mutations acquired for patients with B2M alterations versus those without. **E**. Heatmap showing the extent of associations between alterations of known splicing-related genes and the alternative splicing of COSMIC genes. Each column indicates one COSMIC gene, and the color intensity shows the significance of trans-association. Splicing related genes labeled to the right are ordered by the number of significant associations.

**Extended Data Figure 25.**
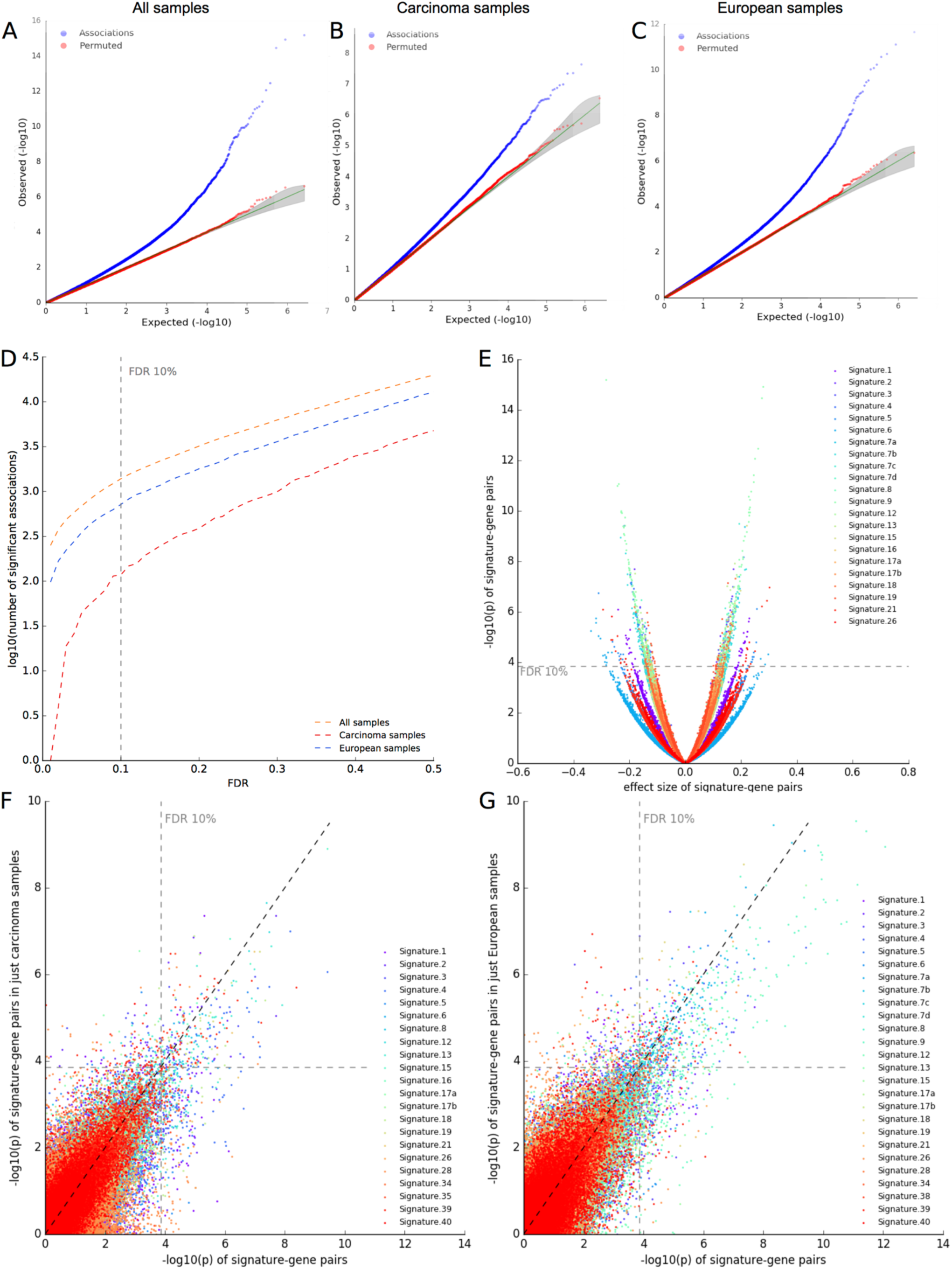
Quality control of the gene expression-mutational signature association studies. **A-C)** QQ plots of the p-values of the linear model associating expression of 18,831 genes with 28 signatures across **A)** all 1,159 patients, **B)** 877 carcinoma patients, or **C)** 891 European patients. **D)** Number of significant associations (log_10_) at different FDR thresholds (across all, carcinoma and European patients). **E)** Volcano plot of directionality of effects in the analysis of all patients. **F-G)** Comparison of analyses between all patients and **F)** carcinoma, **G)** European patients, respectively. The −log_10_P per signature-gene pair are correlated (r=.763 and r=.789, Pearson correlation coefficient), especially above an FDR threshold of 10%.

**Extended Data Figure 26.**
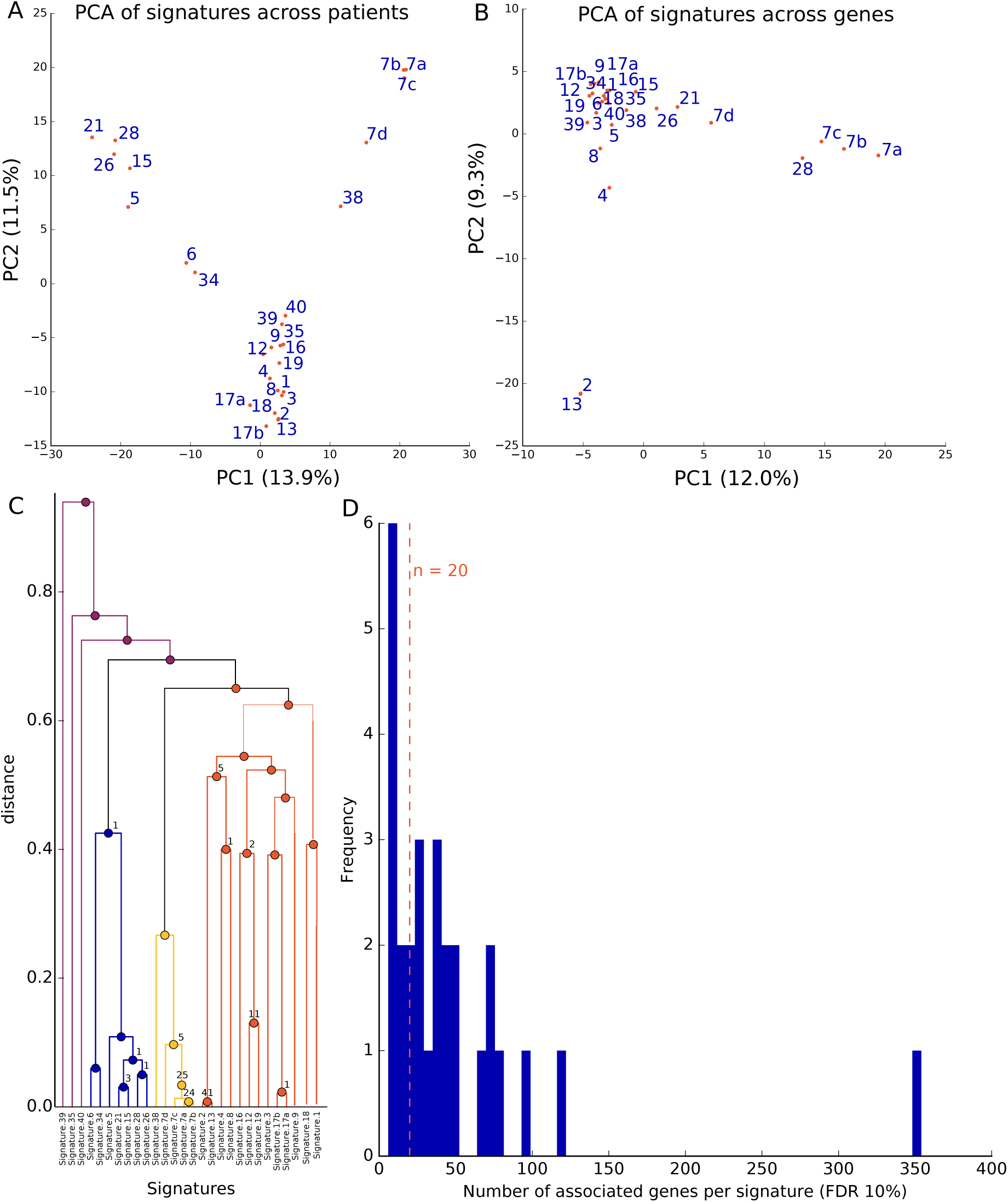
Relationship between mutational signatures and gene expression patterns. **A-B)** PCA of **A)** signatures across 1,159 patients (PCA on signature-specific SNVs per patient) and of **B)** signature-gene expression associations across 18,831 genes (PCA on adjusted p-values of signature-gene expression associations). The PCA on the SNVs recapitulates known interdependencies, e.g. between Signatures 7, whereas the PCA on the signature-gene association studies additionally emphasizes functional relatedness, e.g. between Signatures 2 and 13. **C)** Hierarchical clustering of signatures. The numbers at the nodes indicate the number of genes commonly associated with two to four respective signatures. The dendrogram shows that genes are associated with more than one signature mostly due to similar SNV patterns of these signatures across patients. **D)** Frequency of number of significantly associated genes per signature (FDR ≤ 10%). While many signatures are significantly associated with a few genes, 18 signatures are associated with more than 20 genes. Signature 9 is associated with more than 350 genes. Vice versa, 1009 genes are associated with only one signature, 129 with two, 32 with three, 5 with four and 1 with five signatures.

**Extended Data Figure 27.**
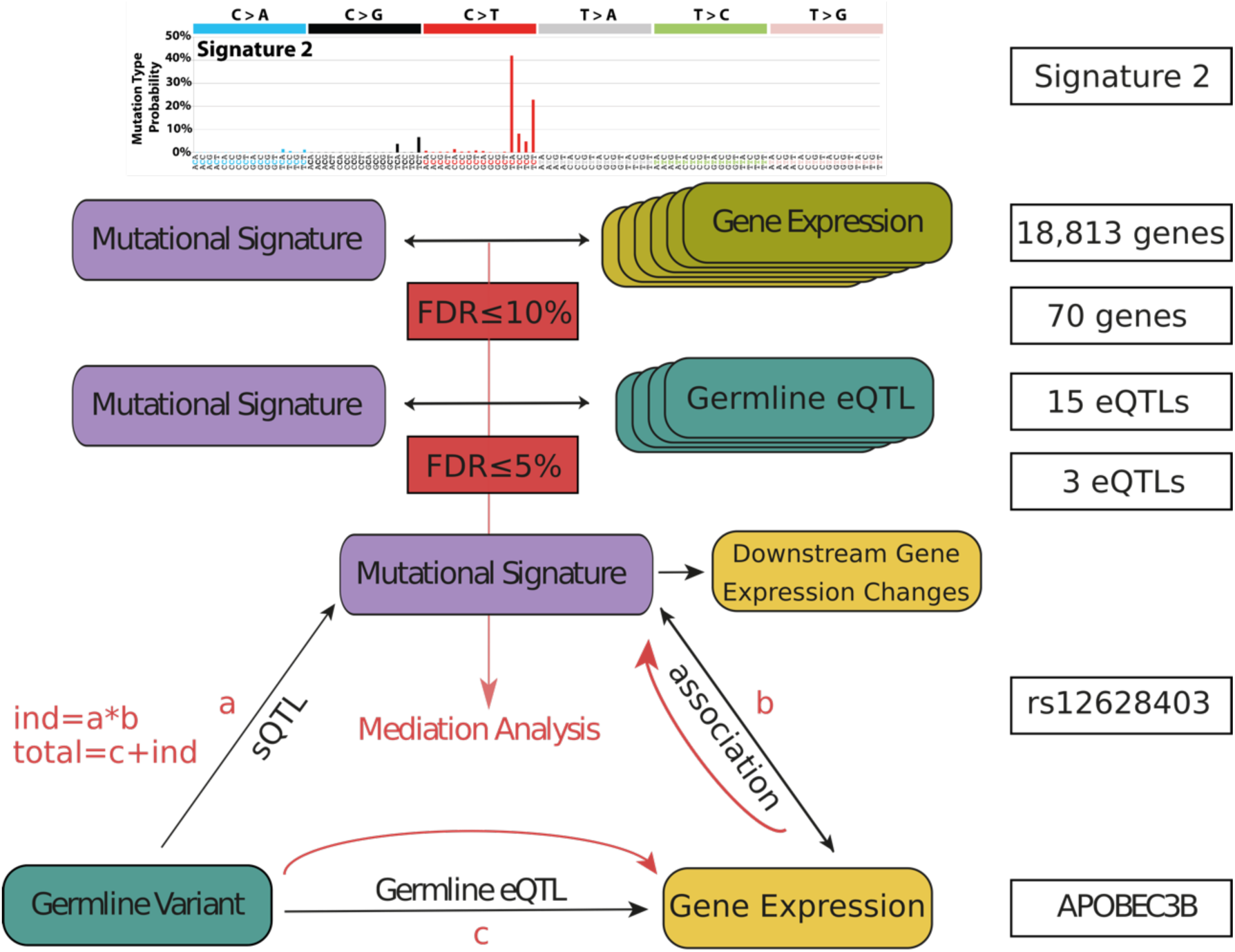
Workflow of gene expression-mutational signature association studies with subsequent mediation analysis including germline eQTL of associated genes. Genes involved in significant mutational signatures and gene expression associations are queried for germline eQTL. If the germline eQTL lead variant is significantly associated with the mutational signature, mediation analysis is applied to each potential triple of germline eQTL lead variant, gene expression and associated mutational signature. Here, the mediating effect of the mutational signature is assessed by comparing the indirect (ind) and total effect of the germline variant onto gene expression (the same analysis has been conducted for gene expression as mediator). a, b and c denote the effect sizes of the individual associations. The boxes on the right show the numbers of genes and eQTL for the *APOBEC3B* case.

**Extended Data Figure 28.**
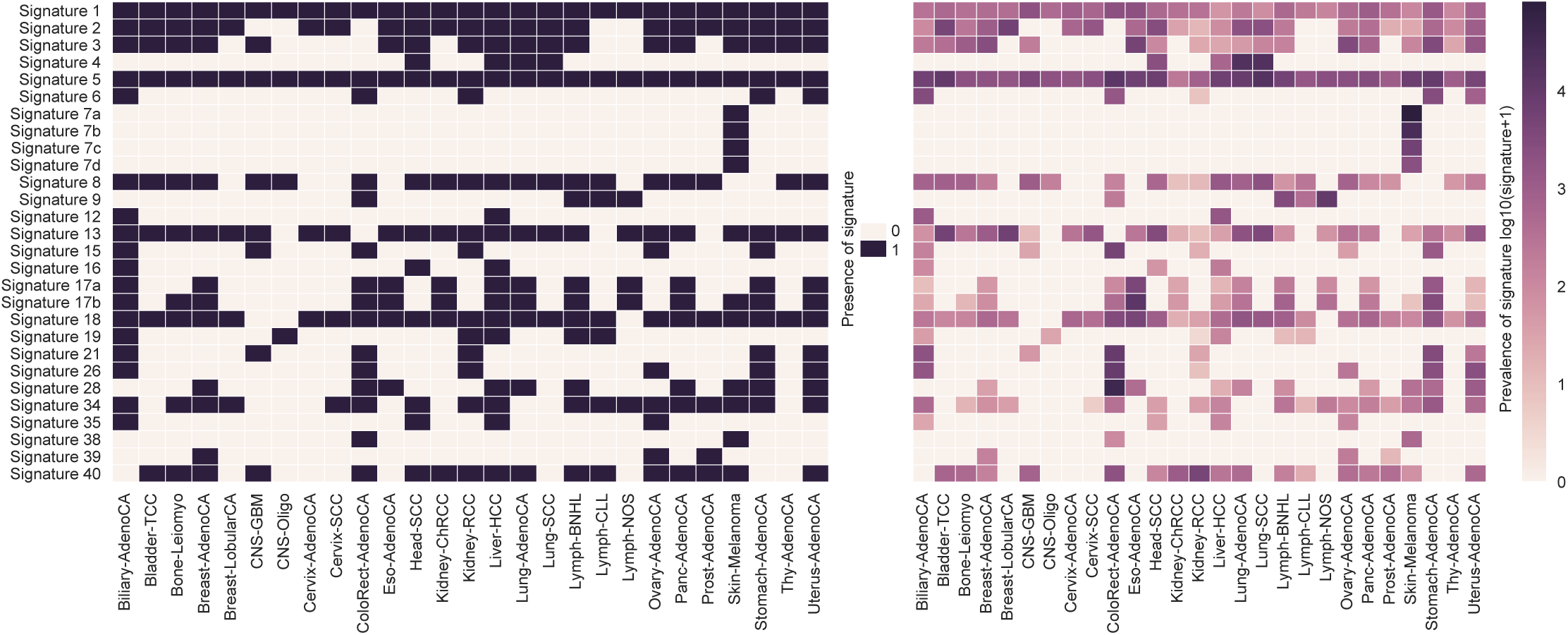
Prevalence of mutational signatures across cancer types. The left heatmap shows the presence of each signature in a specific cancer type (at least one mutation of the respective signature occurs in at least one patient with the specific cancer type). Signatures 1 and 5 occur in all cancer types, signatures 2, 13 and 18 are common signatures and signatures 4, 7, 12, 16, 38 and 39 occur in specific cancer types. The right heatmap shows the prevalence of each signature, i.e. the mean signature count (log_10_(count + 1)) across all patients of one cancer type.

**Extended Data Figure 29.**
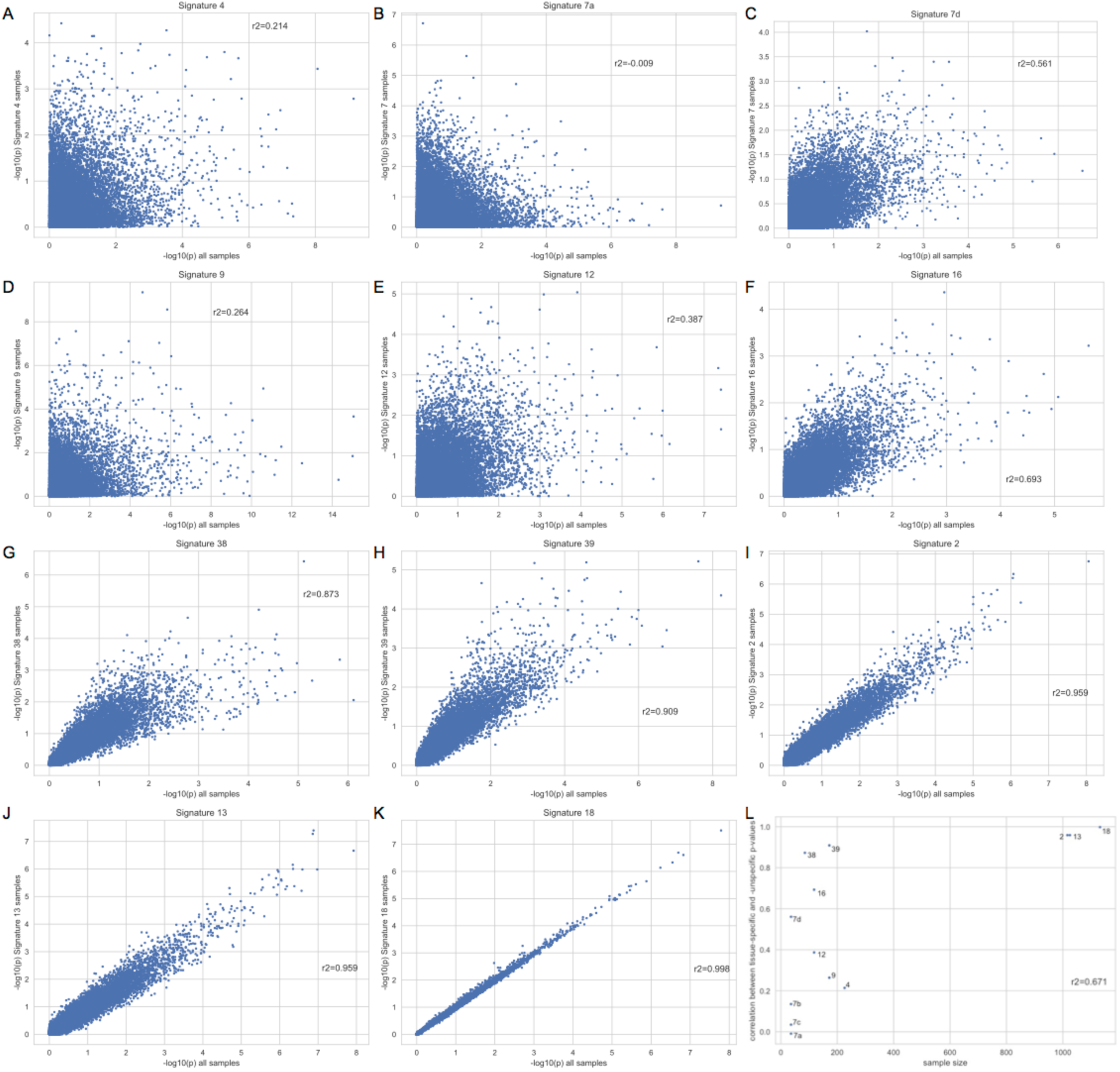
Comparison of the analysis of the whole cohort with cancer type-specific analyses. A-K) The p-values (-log_10_P) of cancer type-specific analyses are compared against the p-values of the analysis applied to the whole cohort and the Pearson correlation coefficient (r^2^) is calculated. Per signature, all cancer types are taken into account that show presence of the specific signature (see Fig. S16). The presented signatures are A-H) cancer type-specific signatures that occur in up to 4 cancer types and I-K) common signatures that are not present in up to 5 cancer types. L) Correlations between cancer type-specific and whole-cohort p-values (r^2^) are plotted over the sample size of the respective cancer types.

## Author Information

**Working group leaders**

Alvis Brazma, Gunnar Rätsch, Angela N. Brooks

**PCAWG Transcriptome Core Group (equal contribution)**

Claudia Calabrese, Natalie R. Davidson, Nuno A. Fonseca, Yao He, André Kahles, Kjong-Van Lehmann, Fenglin Liu, Yuichi Shiraishi, Cameron M. Soulette, Lara Urban

**Data coordination group**

Nuno A. Fonseca (Co-first author), André Kahles (Co-first author), Kjong-Van Lehmann (Co-first author), Marc D. Perry (Co-first author), Linda Xiang, Christina Yung, Junjun Zhang, Katherine A. Hoadley, Peter Bailey, Reiner Siebert, B.F. Francis Ouellette (Principal Investigator), Alvis Brazma (Principal Investigator), Gunnar Rätsch (Principal Investigator), Angela N. Brooks (Principal Investigator)

**RNA-Seq processing group**

Nuno Fonseca (Co-first author), André Kahles (Co-first author), Kjong-Van Lehmann (Co-first author), Chad J. Creighton, Stefan G. Stark, Angela N. Brooks (Principal Investigator), Alvis Brazma (Principal Investigator), Gunnar Rätsch (Principal Investigator)

**eQTL analysis group**

Claudia Calabrese (Co-first author), Kjong-Van Lehmann (Co-first author), Nuno A. Fonseca, André Kahles, Lara Urban, Helena Kilpinen, Sebastian Waszak, Jan Korbel, Alvis Brazma (Principal Investigator), Roland F. Schwarz (Principal Investigator), Gunnar Rätsch (Principal Investigator), Oliver Stegle (Principal Investigator)

**ASE analysis group**

Lara Urban (Co-first author), Fenglin Liu (Co-first author), Helena Kilpinen, Julia Markowski, Serap Erkek, Zemin Zhang (Principal Investigator), Oliver Stegle (Principal Investigator), Roland F. Schwarz (Principal Investigator)

**Alternative splicing analysis group**

André Kahles (Co-first author), Yuichi Shiraishi (Co-first author), Cameron M. Soulette (Co-first author), Kjong-Van Lehmann, Stefan G. Stark, Maximillian G. Marin, Gunnar Rätsch (Principal Investigator), Angela N. Brooks (Principal Investigator)

**Alternative promoter analysis group**

Deniz Demircioglu (First author), Tannistha Nandi, Claudia Calabrese, Kjong-Van Lehmann, Patrick Tan, Jonathan Göke (Principal Investigator)

**Fusion analysis group**

Nuno A. Fonseca (Co-first author), Yao He (Co-first author), Liliana Greger, Alvis Brazma (Principal Investigator), Zemin Zhang (Principal Investigator)

**RNA editing analysis group**

Dongbing Liu (Co-first author), Hong Su, Siliang Li (Co-first author), Yong Hou, Shida Zhu, Qiang Pan-Hammarström, Huangming Yang (Principal Investigator), Kui Wu (Principal Investigator)

**Mutational signature analysis group**

Lara Urban (First author), Sebastian Waszak, Kjong-Van Lehmann, Roland F. Schwarz (Principal Investigator), Oliver Stegle (Principal Investigator)

**Transcriptome meta-analysis group**

Natalie R. Davidson (First author), Fenglin Liu, Kjong-Van Lehmann, Fan Zhang, Deniz Demircioglu, Nuno A. Fonseca, André Kahles, Siliang Li, Roland F. Schwarz, Hong Su, Reiner Siebert, Yao He, Stefan G. Stark, Alvis Brazma (Principal Investigator), Angela N. Brooks (Principal Investigator), Zemin Zhang (Principal Investigator), Gunnar Rätsch (Principal Investigator)

**PCAWG Transcriptome Working Group**

Nuno A. Fonseca (Co-first author), André Kahles (Co-first author), Kjong-Van Lehmann (Co-first author), Claudia Calabrese, Aurélien Chateigner, Natalie R. Davidson, Deniz Demircioglu, Liliana Greger, Yao He, Fabien C. Lamaze, Siliang Li, Dongbing Liu, Fenglin Liu, Marc D. Perry, Yuichi Shiraishi, Cameron M. Soulette, Lara Urban, Linda Xiang, Fan Zhang, Junjun Zhang, Samirkumar B. Amin, Peter Bailey, Isidro Cortés-Ciriano, Brian Craft, Serap Erkek, Milana Frenkel-Morgenstern, Mary Goldman, Katherine A. Hoadley, Yong Hou, Ekta Khurana, Helena Kilpinen, Jan O. Korbel, Chang Li, Xiaobo Li, Xinyue Li, Xingmin Liu, Maximillian G. Marin, Julia Markowski, Tannistha Nandi, Morten M. Nielsen, Akinyemi I. Ojesina, Qiang Pan-Hammarström, Peter J. Park, Chandra Sekhar Pedamallu, Jakob S. Pedersen, Reiner Siebert, Stefan G. Stark, Hong Su, Patrick Tan, Bin Tean Teh, Jian Wang, Sebastian M. Waszak, Heng Xiong, Sergei Yakneen, Chen Ye, Christina Yung, Xiuqing Zhang, Liangtao Zheng, Jingchun Zhu, Shida Zhu, Philip Awadalla, Chad J. Creighton, Matthew Meyerson, B.F. Francis Ouellette, Kui Wu, Huangming Yang, Jonathan Göke, Roland F. Schwarz, Oliver Stegle, Zemin Zhang, Alvis Brazma (Working Group Co-Leader), Gunnar Rätsch (Working Group Co-Leader), Angela N. Brooks (Working Group Co-Leader)

**Affiliations (listed in alphabetical order)**

Aarhus University, Aarhus, DK-8200, Denmark

Morten M. Nielsen, Jakob S. Pedersen

BGI-Shenzhen, Shenzhen, 518083, China

Yong Hou, Chang Li, Siliang Li, Xiaobo Li, Xinyue Li, Dongbing Liu, Xingmin Liu, Qiang Pan-Hammarström, Hong Su, Jian Wang, Kui Wu, Heng Xiong, Huangming Yang, Chen Ye, Xiuqing Zhang, Shida Zhu

The Azrieli Faculty of Medicine, Bar-Ilan University, Safed, 13195, Israel

Milana Frenkel-Morgenstern

Baylor College of Medicine, Houston, 77030, USA

Chad J. Creighton

Berlin Institute for Medical Systems Biology, Max Delbruck Center for Molecular Medicine, Berlin, 13125, Germany

Julia Markowski, Roland F. Schwarz (Senior Author)

Broad Institute, Cambridge, 02142, USA

Angela N. Brooks (Senior Author) (Working Group Leader), Matthew Meyerson, Chandra Sekhar Pedamallu

China National GeneBank-Shenzhen, Shenzhen, 518083, China

Yong Hou, Chang Li, Siliang Li, Xiaobo Li, Dongbing Liu, Xingmin Liu, Hong Su, Kui Wu, Heng Xiong, Chen Ye, Shida Zhu

Dana-Farber Cancer Institute, Boston, 02215, USA

Angela N. Brooks (Senior Author) (Working Group Leader), Matthew Meyerson

Duke-NUS Graduate Medical School, Singapore, 169857, Singapore

Patrick Tan

ETH Zurich, Zurich, 8092, Switzerland

Natalie R. Davidson (Core Group), André Kahles (Core Group), Kjong-Van Lehmann (Core Group), Gunnar Rätsch (Senior Author) (Working Group Leader), Stefan G. Stark

European Molecular Biology Laboratory, Heidelberg, 69117, Germany

Serap Erkek, Jan O. Korbel, Oliver Stegle (Senior Author), Sebastian M. Waszak, Sergei Yakneen

European Molecular Biology Laboratory, Hinxton, CB10 1SD, UK

Alvis Brazma (Senior Author) (Working Group Leader), Claudia Calabrese (Core Group), Nuno A. Fonseca (Core Group), Liliana Greger, Roland F. Schwarz (Senior Author), Oliver Stegle (Senior Author), Lara Urban (Core Group)

Genome Institute of Singapore, Singapore, 138672, Singapore

Deniz Demircioğlu, Jonathan Göke (Senior Author), Tannistha Nandi, Patrick Tan

Harvard Medical School, Boston, 02115, USA

Isidro Cortés-Ciriano, Matthew Meyerson, Peter J. Park

HudsonAlpha Institute for Biotechnology, Birmingham, 35294, USA

Akinyemi I. Ojesina

Karolinska Institutet, Stockholm, 14186, Sweden

Qiang Pan-Hammarström

Ludwig Center at Harvard, Boston, 02215, USA

Isidro Cortés-Ciriano, Peter J. Park

Memorial Sloan Kettering Cancer Center, New York, 10065, USA

Natalie R. Davidson (Core Group), André Kahles (Core Group), Kjong-Van Lehmann (Core Group), Gunnar Rätsch (Senior Author) (Working Group Leader)

National Cancer Centre Singapore, Singapore, 169610, Singapore

Bin Tean Teh, Jonathan Göke

National University of Singapore, Singapore, 117417, Singapore

Deniz Demircioglu

Ontario Institute for Cancer Research, Toronto, M5G 0A3, Canada

Philip Awadalla, Aurélien Chateigner, Fabien C. Lamaze, Marc D. Perry, Linda Xiang, Christina Yung, Junjun Zhang

Peking University, Beijing, 100871, China

Yao He (Core Group), Fenglin Liu (Core Group), Fan Zhang, Zemin Zhang (Senior Author), Liangtao Zheng

SIB Swiss Institute of Bioinformatics, Lausanne, 1015, Switzerland

The UT MD Anderson Cancer Center, Houston, 6032, USA

Samirkumar B. Amin

The University of North Carolina at Chapel Hill, Chapel Hill, 27599, USA

Katherine A. Hoadley

The University of Tokyo, Minato-ku, 108-8639, Japan

Yuichi Shiraishi (Core Group)

Ulm University & Ulm University Medical Center, Ulm, 89081, Germany

Reiner Siebert

University College London, London, WC1N 1EH, UK

Helena Kilpinen

University Hospital Zurich, Zurich, 8091, Switzerland

Gunnar Rätsch (Senior Author) (Working Group Leader)

University of Alabama at Birmingham, Birmingham, 35294, USA

Akinyemi I. Ojesina

University of California, San Francisco, San Francisco, 94143, USA

Marc D. Perry

University of California, Santa Cruz, Santa Cruz, 95064, USA

Angela N. Brooks (Senior Author) (Working Group Leader), Brian Craft, Mary Goldman, Maximillian G. Marin, Cameron M. Soulette (Core Group), Jingchun Zhu

University of Cambridge, Cambridge, CB2 1EW, UK

Isidro Cortés-Ciriano

University of Glasgow, Glasgow, G61 1BD, UK

Peter Bailey

University of Toronto, Toronto, M8G 0A8, Canada

Philip Awadalla, B.F. Francis Ouellette

Weill Cornell Medical College, New York, 10065, USA

Natalie R. Davidson (Core Group), Ekta Khurana, Gunnar Rätsch (Senior Author) (Working Group Leader)

## References

1. Faderl, S. et al. The biology of chronic myeloid leukemia. N. Engl. J. Med. 341, 164–172 (1999).

2. Owens, M. A., Horten, B. C. & Da Silva, M. M. HER2 amplification ratios by fluorescence in situ hybridization and correlation with immunohistochemistry in a cohort of 6556 breast cancer tissues. Clin. Breast Cancer 5, 63–69 (2004).

3. Cancer Genome Atlas Research Network et al. Genomic and epigenomic landscapes of adult de novo acute myeloid leukemia. N. Engl. J. Med. 368, 2059–2074 (2013).

4. Cancer Genome Atlas Research Network. Comprehensive molecular profiling of lung adenocarcinoma. Nature 511, 543–550 (2014).

5. Climente-González, H., Porta-Pardo, E., Godzik, A. & Eyras, E. The Functional Impact of Alternative Splicing in Cancer. Cell Rep. 20, 2215–2226 (2017).

6. Muratani, M. et al. Nanoscale chromatin profiling of gastric adenocarcinoma reveals cancer-associated cryptic promoters and somatically acquired regulatory elements. Nat. Commun. 5, 4361 (2014).

7. Douillard, J.-Y. et al. Relationship between EGFR expression, EGFR mutation status, and the efficacy of chemotherapy plus cetuximab in FLEX study patients with advanced non-small-cell lung cancer. J. Thorac. Oncol. 9, 717–724 (2014).

8. Uhlen, M. et al. A pathology atlas of the human cancer transcriptome. Science 357, (2017).

9. Rheinbay, E. et al. Recurrent and functional regulatory mutations in breast cancer. Nature 547, 55–60 (2017).

10. Weinhold, N., Jacobsen, A., Schultz, N., Sander, C. & Lee, W. Genome-wide analysis of noncoding regulatory mutations in cancer. Nat. Genet. 46, 1160–1165 (2014).

11. PCAWG Consortium et al. PCAWG Marker paper. In preparation (2018).

12. Dobin, A. et al. STAR: ultrafast universal RNA-seq aligner. Bioinformatics 29, 15–21 (2013).

13. Kim, D. et al. TopHat2: accurate alignment of transcriptomes in the presence of insertions, deletions and gene fusions. Genome Biol. 14, R36 (2013).

14. Anders, S., Pyl, P. T. & Huber, W. HTSeq--a Python framework to work with high-throughput sequencing data. Bioinformatics 31, 166–169 (2015).

15. Bullard, J. H., Purdom, E., Hansen, K. D. & Dudoit, S. Evaluation of statistical methods for normalization and differential expression in mRNA-Seq experiments. BMC Bioinformatics 11, 94 (2010).

16. Dillies, M.-A. et al. A comprehensive evaluation of normalization methods for Illumina high-throughput RNA sequencing data analysis. Brief. Bioinform. 14, 671–683 (2013).

17. Li, B. & Dewey, C. N. RSEM: accurate transcript quantification from RNA-Seq data with or without a reference genome. BMC Bioinformatics 12, 323 (2011).

18. Bray, N. L., Pimentel, H., Melsted, P. & Pachter, L. Near-optimal probabilistic RNA-seq quantification. Nat. Biotechnol. 34, 525–527 (2016).

19. Stegle, O., Parts, L., Piipari, M., Winn, J. & Durbin, R. Using probabilistic estimation of expression residuals (PEER) to obtain increased power and interpretability of gene expression analyses. Nat. Protoc. 7, 500–507 (2012).

20. Bryois, J. et al. Cis and trans effects of human genomic variants on gene expression. PLoS Genet. 10, e1004461 (2014).

21. GTEx Consortium et al. Genetic effects on gene expression across human tissues. Nature 550, 204–213 (2017).

22. Kilpinen, H. et al. Common genetic variation drives molecular heterogeneity in human iPSCs. Nature 546, 370–375 (2017).

23. Zhang, X. A., Lane, W. S., Charrin, S., Rubinstein, E. & Liu, L. EWI2/PGRL Associates with the Metastasis Suppressor KAI1/CD82 and Inhibits the Migration of Prostate Cancer Cells. Cancer Res. 63, 2665–2674 (2003).

24. Saito, T. T., Mohideen, F., Meyer, K., Harper, J. W. & Colaiácovo, M. P. SLX-1 is required for maintaining genomic integrity and promoting meiotic noncrossovers in the Caenorhabditis elegans germline. PLoS Genet. 8, e1002888 (2012).

25. Medves, S. et al. A high rate of telomeric sister chromatid exchange occurs in chronic lymphocytic leukaemia B-cells. Br. J. Haematol. 174, 57–70 (2016).

26. Wang, R.-F. & Wang, H. Y. Immune targets and neoantigens for cancer immunotherapy and precision medicine. Cell Res. 27, 11–37 (2017).

27. Scanlan, M. J., Gure, A. O., Jungbluth, A. A., Old, L. J. & Chen, Y.-T. Cancer/testis antigens: an expanding family of targets for cancer immunotherapy. Immunol. Rev. 188, 22–32 (2002).

28. Simpson, A. J. G., Caballero, O. L., Jungbluth, A., Chen, Y.-T. & Old, L. J. Cancer/testis antigens, gametogenesis and cancer. Nat. Rev. Cancer 5, 615–625 (2005).

29. Bajrami, I. et al. Genome-wide Profiling of Genetic Synthetic Lethality Identifies CDK12 as a Novel Determinant of PARP1/2 Inhibitor Sensitivity. Cancer Res. 74, 287–297 (2013).

30. Ekumi, K. M. et al. Ovarian carcinoma CDK12 mutations misregulate expression of DNA repair genes via deficient formation and function of the Cdk12/CycK complex. Nucleic Acids Res. 43, 2575–2589 (2015).

31. Ilboudo, A. et al. Overexpression of phosphatidylinositol 4-kinase type IIIα is associated with undifferentiated status and poor prognosis of human hepatocellular carcinoma. BMC Cancer 14, 7 (2014).

32. Havelange, V. et al. IRF4 mutations in chronic lymphocytic leukemia. Blood 118, 2827– 2829 (2011).

33. Nonaka, T. et al. Involvement of activation-induced cytidine deaminase in skin cancer development. J. Clin. Invest. 126, 1367–1382 (2016).

34. Bhalla, S. et al. Gene expression-based biomarkers for discriminating early and late stage of clear cell renal cancer. Sci. Rep. 7, 44997 (2017).

35. Hartmann, S. et al. Highly recurrent mutations of SGK1, DUSP2 and JUNB in nodular lymphocyte predominant Hodgkin lymphoma. Leukemia 30, 844–853 (2016).

36. Scholtysik, R. et al. Detection of genomic aberrations in molecularly defined Burkitt’s lymphoma by array-based, high resolution, single nucleotide polymorphism analysis. Haematologica 95, 2047–2055 (2010).

37. Boerma, E. G., Siebert, R., Kluin, P. M. & Baudis, M. Translocations involving 8q24 in Burkitt lymphoma and other malignant lymphomas: a historical review of cytogenetics in the light of todays knowledge. Leukemia 23, 225–234 (2009).

38. Lesch, B. J. & Page, D. C. Poised chromatin in the mammalian germ line. Development 141, 3619–3626 (2014).

39. Bernhart, S. H. et al. Changes of bivalent chromatin coincide with increased expression of developmental genes in cancer. Sci. Rep. 6, 37393 (2016).

40. Hanafusa, T., Mohamed, A. E. A., Domae, S., Nakayama, E. & Ono, T. Serological identification of Tektin5 as a cancer/testis antigen and its immunogenicity. BMC Cancer 12, 520 (2012).

41. Castel, S. E., Levy-Moonshine, A., Mohammadi, P., Banks, E. & Lappalainen, T. Tools and best practices for data processing in allelic expression analysis. Genome Biol. 16, (2015).

42. Castel, S. E., Levy-Moonshine, A., Mohammadi, P., Banks, E. & Lappalainen, T. Tools and best practices for data processing in allelic expression analysis. Genome Biol. 16, 195 (2015).

43. Ha, G. et al. Integrative analysis of genome-wide loss of heterozygosity and monoallelic expression at nucleotide resolution reveals disrupted pathways in triple-negative breast cancer. Genome Res. 22, 1995–2007 (2012).

44. Lindeboom, R. G. H., Supek, F. & Lehner, B. The rules and impact of nonsense-mediated mRNA decay in human cancers. Nat. Genet. 48, 1112–1118 (2016).

45. Gerstung, M. et al. The evolutionary history of 2,658 cancers. bioRxiv 161562 (2017). doi:10.1101/161562

46. PCAWG Group et al. PCAWG 11 Marker Paper. In preparation (2018).

47. Pecci, A., Viegas, L. R., Barañao, J. L. & Beato, M. Promoter Choice Influences Alternative Splicing and Determines the Balance of Isoforms Expressed from the Mousebcl-X Gene. J. Biol. Chem. 276, 21062–21069 (2001).

48. Xin, D., Hu, L. & Kong, X. Alternative Promoters Influence Alternative Splicing at the Genomic Level. PLoS One 3, e2377 (2008).

49. Sabarinathan, R., Mularoni, L., Deu-Pons, J., Gonzalez-Perez, A. & López-Bigas, N. Nucleotide excision repair is impaired by binding of transcription factors to DNA. Nature 532, 264–267 (2016).

50. Perera, D. et al. Differential DNA repair underlies mutation hotspots at active promoters in cancer genomes. Nature 532, 259–263 (2016).

51. Huang, F. W. et al. Highly recurrent TERT promoter mutations in human melanoma. Science 339, 957–959 (2013).

52. Horn, S. et al. TERT promoter mutations in familial and sporadic melanoma. Science 339, 959–961 (2013).

53. Rheinbay, E., Nielsen, M. M., Abascal, F. & Tiao, G. Discovery and characterization of coding and non-coding driver mutations in more than 2,500 whole cancer genomes. BioRxiv (2017).

54. Kahles, A., Ong, C. S., Zhong, Y. & Rätsch, G. SplAdder: identification, quantification and testing of alternative splicing events from RNA-Seq data. Bioinformatics 32, 1840–1847 (2016).

55. Jung, H. et al. Intron retention is a widespread mechanism of tumor-suppressor inactivation. Nat. Genet. 47, 1242–1248 (2015).

56. Wang, L. et al. Transcriptomic Characterization of SF3B1 Mutation Reveals Its Pleiotropic Effects in Chronic Lymphocytic Leukemia. Cancer Cel l 30, 750–763 (2016).

57. Signal, B., Gloss, B. S., Dinger, M. E. & Mercer, T. R. Machine-learning annotation of human splicing branchpoints. bioRxiv 094003 (2016). doi:10.1101/094003

58. Mercer, T. R. et al. Genome-wide discovery of human splicing branchpoints. Genome Res. 25, 290–303 (2015).

59. Shiraishi, Y. et al. A comprehensive characterization of cis-acting splicing-associated variants in human cancer. bioRxiv (2017).

60. Schwartz, S. et al. Alu exonization events reveal features required for precise recognition of exons by the splicing machinery. PLoS Comput. Biol. 5, e1000300 (2009).

61. Sorek, R. The birth of new exons: mechanisms and evolutionary consequences. rnajournal.cshlp.org (2007).

62. Mertens, F., Johansson, B., Fioretos, T. & Mitelman, F. The emerging complexity of gene fusions in cancer. Nat. Rev. Cancer 15, 371–381 (2015).

63. Melé, M. et al. Human genomics. The human transcriptome across tissues and individuals. Science 348, 660–665 (2015).

64. Yoshihara, K. et al. The landscape and therapeutic relevance of cancer-associated transcript fusions. Oncogene 34, 4845–4854 (2015).

65. Matsubara, D. et al. Identification of CCDC6-RET fusion in the human lung adenocarcinoma cell line, LC-2/ad. J. Thorac. Oncol. 7, 1872–1876 (2012).

66. Carneiro, B. A. et al. FGFR3-TACC3: A novel gene fusion in cervical cancer. Gynecol Oncol Rep 13, 53–56 (2015).

67. Lee, M. et al. ChimerDB 3.0: an enhanced database for fusion genes from cancer transcriptome and literature data mining. Nucleic Acids Res. 45, D784–D789 (2017).

68. Xu, W. S., Liang, R. H. & Srivastava, G. Identification and characterization of BCL6 translocation partner genes in primary gastric high-grade B-cell lymphoma: heat shock protein 89 alpha is a novel fusion partner gene of BCL6. Genes Chromosomes Cancer 27, 69–75 (2000).

69. Le Tallec, B. et al. Common fragile site profiling in epithelial and erythroid cells reveals that most recurrent cancer deletions lie in fragile sites hosting large genes. Cell Rep. 4, 420–428 (2013).

70. Knezevich, S. R., McFadden, D. E., Tao, W., Lim, J. F. & Sorensen, P. H. A novel ETV6-NTRK3 gene fusion in congenital fibrosarcoma. Nat. Genet. 18, 184–187 (1998).

71. Nacu, S. et al. Deep RNA sequencing analysis of readthrough gene fusions in human prostate adenocarcinoma and reference samples. BMC Med. Genomics 4, 11 (2011).

72. Jia, Y., Xie, Z. & Li, H. Intergenically Spliced Chimeric RNAs in Cancer. Trends Cancer Res. 2, 475–484 (2016).

73. Greger, L. et al. Tandem RNA chimeras contribute to transcriptome diversity in human population and are associated with intronic genetic variants. PLoS One 9, e104567 (2014).

74. Dallery, E. et al. TTF, a gene encoding a novel small G protein, fuses to the lymphoma-associated LAZ3 gene by t(3;4) chromosomal translocation. Oncogene 10, 2171–2178 (1995).

75. Tomlins, S. A. Recurrent Fusion of TMPRSS2 and ETS Transcription Factor Genes in Prostate Cancer. Science 310, 644–648 (2005).

76. Stenson, P. D. et al. The Human Gene Mutation Database: towards a comprehensive repository of inherited mutation data for medical research, genetic diagnosis and next-generation sequencing studies. Hum. Genet. 136, 665–677 (2017).

77. Koboldt, D. C. et al. VarScan 2: somatic mutation and copy number alteration discovery in cancer by exome sequencing. Genome Res. 22, 568–576 (2012).

78. Campbell, P. J. et al. Pan-cancer analysis of whole genomes. bioRxiv 162784 (2017). doi:10.1101/162784

79. Forbes, S. A. et al. COSMIC: somatic cancer genetics at high-resolution. Nucleic Acids Res. 45, D777–D783 (2017).

80. PCAWG-2-5-9-14. PCAWG-2-5-9-14 Marker Paper (Drivers).

81. Lawrence, M. S. et al. Mutational heterogeneity in cancer and the search for new cancer-associated genes. Nature 499, 214–218 (2013).

82. Klijn, C. et al. A comprehensive transcriptional portrait of human cancer cell lines. Nat. Biotechnol. 33, 306–312 (2015).

83. Garraway, L. A. & Lander, E. S. Lessons from the cancer genome. Cell 153, 17–37 (2013).

84. Thomas, R. K. et al. High-throughput oncogene mutation profiling in human cancer. Nat.Genet. 39, 347–351 (2007).

85. Tranchant, R. et al. Co-occurring Mutations of Tumor Suppressor Genes, LATS2 and NF2, in Malignant Pleural Mesothelioma. Clin. Cancer Res. 23, 3191–3202 (2017).

86. Li, Z. et al. A global transcriptional regulatory role for c-Myc in Burkitt’s lymphoma cells. Proc. Natl. Acad. Sci. U. S. A. 100, 8164–8169 (2003).

87. Kress, T. R., Sabò, A. & Amati, B. MYC: connecting selective transcriptional control to global RNA production. Nat. Rev. Cancer 15, 593–607 (2015).

88. Li, Z. & Nabel, G. J. A new member of the I kappaB protein family, I kappaB epsilon, inhibits RelA (p65)-mediated NF-kappaB transcription. Mol. Cell. Biol. 17, 6184–6190 (1997).

89. Mansouri, L. et al. Functional loss of IκBε leads to NF-κB deregulation in aggressive chronic lymphocytic leukemia. J. Exp. Med. 212, 833–843 (2015).

90. Ausserlechner, M. J., Obexer, P., Böck, G., Geley, S. & Kofler, R. Cyclin D3 and c-MYC control glucocorticoid-induced cell cycle arrest but not apoptosis in lymphoblastic leukemia cells. Cell Death Differ. 11, 165–174 (2004).

91. del Campo, A. B. et al. Immune escape of cancer cells with beta2-microglobulin loss over the course of metastatic melanoma. Int. J. Cancer 134, 102–113 (2014).

92. Adler, A. S. et al. An integrative analysis of colon cancer identifies an essential function for PRPF6 in tumor growth. Genes Dev. 28, 1068–1084 (2014).

93. Luo, N. et al. SAMD4B, a novel SAM-containing protein, inhibits AP-1-, p53- and p21-mediated transcriptional activity. BMB Rep. 43, 355–361 (2010).

94. Alexandrov, L. B. et al. Signatures of mutational processes in human cancer. Nature 500, 415–421 (2013).

95. PCAWG Group et al. PCAWG-7 Marker Paper. In preparation (X) (2018).

96. Fabregat, A. et al. The Reactome pathway Knowledgebase. Nucleic Acids Res. 44, D481–7 (2016).

97. Milacic, M. et al. Annotating cancer variants and anti-cancer therapeutics in reactome. Cancers 4, 1180–1211 (2012).

98. Woenckhaus, M. et al. Smoking and cancer-related gene expression in bronchial epithelium and non-small-cell lung cancers. J. Pathol. 210, 192–204 (2006).

99. Ingram, W. J. et al. ABC transporter activity linked to radiation resistance and molecular subtype in pediatric medulloblastoma. Exp. Hematol. Oncol. 2, 26 (2013).

100. Kvam, E. & Tyrrell, R. M. The role of melanin in the induction of oxidative DNA base damage by ultraviolet A irradiation of DNA or melanoma cells. J. Invest. Dermatol. 113, 209–213 (1999).

101. Denat, L., Kadekaro, A. L., Marrot, L., Leachman, S. A. & Abdel-Malek, Z. A. Melanocytes as instigators and victims of oxidative stress. J. Invest. Dermatol. 134, 1512–1518 (2014).

102. Jimbow, K., Chen, H., Park, J. S. & Thomas, P. D. Increased sensitivity of melanocytes to oxidative stress and abnormal expression of tyrosinase-related protein in vitiligo. Br. J. Dermatol. 144, 55–65 (2001).

103. Premi, S. & Brash, D. E. Unanticipated role of melanin in causing carcinogenic cyclobutane pyrimidine dimmers. Mol Cell Onco l 3, e1033588 (2016).

104. Waszak, S. M. et al. Germline determinants of the somatic mutation landscape in 2,642 cancer genomes. bioRxiv 208330 (2017). doi:10.1101/208330

105. PCAWG Group 8. PCAWG-8 Marker Paper. Nature (2017).

106. Middlebrooks, C. D. et al. Association of germline variants in the APOBEC3 region with cancer risk and enrichment with APOBEC-signature mutations in tumors. Nat. Genet. 48, 1330–1338 (2016).

107. Wallace, C. Statistical testing of shared genetic control for potentially related traits. Genet. Epidemiol. 37, 802–813 (2013).

108. Baron, R. M. & Kenny, D. A. The moderator-mediator variable distinction in social psychological research: conceptual, strategic, and statistical considerations. J. Pers. Soc. Psychol. 51, 1173–1182 (1986).

109. Preacher, K. J. & Hayes, A. F. SPSS and SAS procedures for estimating indirect effects in simple mediation models. Behav. Res. Methods Instrum. Comput. 36, 717–731 (2004).

110. Stransky, N., Cerami, E., Schalm, S., Kim, J. L. & Lengauer, C. The landscape of kinase fusions in cancer. Nat. Commun. 5, 4846 (2014).

111. Cancer Genome Atlas Research Network et al. Comprehensive Molecular Characterization of Papillary Renal-Cell Carcinoma. N. Engl. J. Med. 374, 135–145 (2016).

112. Vogelstein, B. et al. Cancer genome landscapes. Science 339, 1546–1558 (2013).

113. Leiserson, M. D. M. et al. Pan-cancer network analysis identifies combinations of rare somatic mutations across pathways and protein complexes. Nat. Genet. 47, 106–114 (2015).

114. Popova, T. et al. Ovarian Cancers Harboring Inactivating Mutations in CDK12 Display a Distinct Genomic Instability Pattern Characterized by Large Tandem Duplications. Cancer Res. 76, 1882–1891 (2016).

115. Joshi, P. M., Sutor, S. L., Huntoon, C. J. & Karnitz, L. M. Ovarian cancer-associated mutations disable catalytic activity of CDK12, a kinase that promotes homologous recombination repair and resistance to cisplatin and poly(ADP-ribose) polymerase inhibitors. J. Biol. Chem. 289, 9247–9253 (2014).

116. Tien, J. F. et al. CDK12 regulates alternative last exon mRNA splicing and promotes breast cancer cell invasion. Nucleic Acids Res. 45, 6698–6716 (2017).

117. Blazek, D. et al. The Cyclin K/Cdk12 complex maintains genomic stability via regulation of expression of DNA damage response genes. Genes Dev. 25, 2158–2172 (2011).

118. Leiserson, M. D. M., Reyna, M. A. & Raphael, B. J. A weighted exact test for mutually exclusive mutations in cancer. Bioinformatics 32, i736–i745 (2016).

119. Garaud, S. & Willard-Gallo, K. IRF5: a rheostat for tumor-infiltrating lymphocyte trafficking in breast cancer? Immunol. Cell Biol. 93, 425–426 (2015).

120. Hu, G. & Barnes, B. J. IRF-5 is a mediator of the death receptor-induced apoptotic signaling pathway. J. Biol. Chem. 284, 2767–2777 (2009).

121. Hu, G., Mancl, M. E. & Barnes, B. J. Signaling through IFN regulatory factor-5 sensitizes p53-deficient tumors to DNA damage-induced apoptosis and cell death. Cancer Res. 65, 7403–7412 (2005).

122. Yanai, H. et al. Role of IFN regulatory factor 5 transcription factor in antiviral immunity and tumor suppression. Proc. Natl. Acad. Sci. U. S. A. 104, 3402–3407 (2007).

123. Tsunoda, T. & Shirasawa, S. Roles of ZFAT in haematopoiesis, angiogenesis and cancer development. Anticancer Res. 33, 2833–2837 (2013).

124. Bärlund, M. et al. Cloning of BCAS3 (17q23) and BCAS4 (20q13) genes that undergo amplification, overexpression, and fusion in breast cancer. Genes Chromosomes Cancer 35, 311–317 (2002).

125. O’Malley, B. W. & Kumar, R. Nuclear receptor coregulators in cancer biology. Cancer Res. 69, 8217–8222 (2009).

126. Henson, B. J. & Gollin, S. M. Overexpression of KLF13 and FGFR3 in oral cancer cells. Cytogenet. Genome Res. 128, 192–198 (2010).

127. Kim, J.-A. et al. Comprehensive functional analysis of the tousled-like kinase 2 frequently amplified in aggressive luminal breast cancers. Nat. Commun. 7, 12991 (2016).

128. Zarow, C. & Victoroff, J. Increased apolipoprotein E mRNA in the hippocampus in Alzheimer disease and in rats after entorhinal cortex lesioning. Exp. Neurol. 149, 79–86 (1998).

129. Mertins, P. et al. Proteogenomics connects somatic mutations to signalling in breast cancer. Nature 534, 55–62 (2016).

130. Arafat, H. et al. Tumor-specific expression and alternative splicing of the COL-A3 gene in pancreatic cancer. Surgery 150, 306–315 (2011).

131. Xie, X., Liu, X., Zhang, Q. & Yu, J. Overexpression of collagen VI α3 in gastric cancer. Oncol. Lett. 7, 1537–1543 (2014).

132. Livingstone, C. IGF2 and cancer. Endocr. Relat. Cancer 20, R321–39 (2013).

133. Sabarinathan, R., Pich, O., Martincorena, I. & Rubio-Perez, C. The whole-genome panorama of cancer drivers. bioRxiv (2017).

134. Dawson, M. A. & Kouzarides, T. Cancer epigenetics: from mechanism to therapy. Cell 150, 12–27 (2012).

135. Weischenfeldt, J. et al. Pan-cancer analysis of somatic copy-number alterations implicates IRS4 and IGF2 in enhancer hijacking. Nat. Genet. 49, 65–74 (2017).

136. Zhang, X. et al. Identification of focally amplified lineage-specific super-enhancers in human epithelial cancers. Nat. Genet. 48, 176–182 (2016).

137. Dobin, A. et al. STAR: ultrafast universal RNA-seq aligner. Bioinformatics 29, 15–21 (2013).

138. Kim, D. et al. TopHat2: accurate alignment of transcriptomes in the presence of insertions, deletions and gene fusions. Genome Biol. 14, R36 (2013).

139. Fonseca, N. A., Petryszak, R., Marioni, J. & Brazma, A. iRAP - an integrated RNA-seq Analysis Pipeline. (2014). doi:10.1101/005991

140. Bioinformatics, B. FastQC: a quality control tool for high throughput sequence data. Cambridge, UK: Babraham Institute (2011).

141. Cancer Genome Atlas Research Network. The Molecular Taxonomy of Primary Prostate Cancer. Cell 163, 1011–1025 (2015).

142. Li, H. et al. The Sequence Alignment/Map format and SAMtools. Bioinformatics 25, 2078– 2079 (2009).

143. Bray, N. L., Pimentel, H., Melsted, P. & Pachter, L. Near-optimal probabilistic RNA-seq quantification. Nat. Biotechnol. 34, 525–527 (2016).

144. Mortazavi, A., Williams, B. A., McCue, K., Schaeffer, L. & Wold, B. Mapping and quantifying mammalian transcriptomes by RNA-Seq. Nat. Methods 5, 621–628 (2008).

145. Krijthe, J. H. Rtsne: T-Distributed Stochastic Neighbor Embedding using Barnes-Hut Implementation. (2015).

146. PCAWG Group et al. PCAWG-1 Marker Paper. In preparation (2018).

147. PCAWG Group et al. PCAWG-3 Marker Paper. In preparation (2018).

148. Stegle, O., Parts, L., Piipari, M., Winn, J. & Durbin, R. Using probabilistic estimation of expression residuals (PEER) to obtain increased power and interpretability of gene expression analyses. Nat. Protoc. 7, 500–507 (2012).

149. GTEx Consortium. The Genotype-Tissue Expression (GTEx) project. Nat. Genet. 45, 580– 585 (2013).

150. Ashburner, M. et al. Gene ontology: tool for the unification of biology. The Gene Ontology Consortium. Nat. Genet. 25, 25–29 (2000).

151. Gene Ontology Consortium. Gene Ontology Consortium: going forward. Nucleic Acids Res. 43, D1049–56 (2015).

152. Fabregat, A. et al. The Reactome pathway Knowledgebase. Nucleic Acids Res. 44, D481–7 (2016).

153. Milacic, M. et al. Annotating cancer variants and anti-cancer therapeutics in reactome. Cancers 4, 1180–1211 (2012).

154. Durinck, S., Spellman, P. T., Birney, E. & Huber, W. Mapping identifiers for the integration of genomic datasets with the R/Bioconductor package biomaRt. Nat. Protoc. 4, 1184–1191 (2009).

155. Durinck, S. et al. BioMart and Bioconductor: a powerful link between biological databases and microarray data analysis. Bioinformatics 21, 3439–3440 (2005).

156. Yu, G., Wang, L.-G., Han, Y. & He, Q.-Y. clusterProfiler: an R package for comparing biological themes among gene clusters. OMICS 16, 284–287 (2012).

157. Yu, G. & He, Q.-Y. ReactomePA: an R/Bioconductor package for reactome pathway analysis and visualization. Mol. Biosyst. 12, 477–479 (2016).

158. Li, H. A statistical framework for SNP calling, mutation discovery, association mapping and population genetical parameter estimation from sequencing data. Bioinformatics 27, 2987– 2993 (2011).

159. Lippert, C., Casale, F. P., Rakitsch, B. & Stegle, O. LIMIX: genetic analysis of multiple traits. bioRxiv 003905 (2014). doi:10.1101/003905

160. Davis, J. R. et al. An Efficient Multiple-Testing Adjustment for eQTL Studies that Accounts for Linkage Disequilibrium between Variants. Am. J. Hum. Genet. 98, 216–224 (2016).

161. GTEx Consortium et al. Genetic effects on gene expression across human tissues. Nature 550, 204–213 (2017).

162. Kilpinen, H. et al. Common genetic variation drives molecular heterogeneity in human iPSCs. Nature 546, 370–375 (2017).

163. McLaren, W. et al. The Ensembl Variant Effect Predictor. Genome Biol. 17, 122 (2016).

164. Fan, Y. et al. MuSE: accounting for tumor heterogeneity using a sample-specific error model improves sensitivity and specificity in mutation calling from sequencing data. Genome Biol. 17, 178 (2016).

165. Harrow, J. et al. GENCODE: the reference human genome annotation for The ENCODE Project. Genome Res. 22, 1760–1774 (2012).

166. Quinlan, A. R. & Hall, I. M. BEDTools: a flexible suite of utilities for comparing genomic features. Bioinformatics 26, 841–842 (2010).

167. Lippert, C., Casale, F. P., Rakitsch, B. & Stegle, O. LIMIX: genetic analysis of multiple traits. bioRxiv 003905 (2014). doi:10.1101/003905

168. Roadmap Epigenomics Consortium et al. Integrative analysis of 111 reference human epigenomes. Nature 518, 317–330 (2015).

169. ENCODE Project Consortium. An integrated encyclopedia of DNA elements in the human genome. Nature 489, 57–74 (2012).

170. Wallace, C. Statistical testing of shared genetic control for potentially related traits. Genet. Epidemiol. 37, 802–813 (2013).

171. Rosseel, Y. lavaan: AnRPackage for Structural Equation Modeling. J. Stat. Softw. 48, (2012).

172. Tingley, D., Yamamoto, T., Hirose, K., Keele, L. & Imai, K. mediation:RPackage for Causal Mediation Analysis. J. Stat. Softw. 59, (2014).

173. Frith, M. C. et al. A code for transcription initiation in mammalian genomes. Genome Res. 18, 1–12 (2008).

174. Demircioğlu, D. et al. A pan cancer analysis of promoter activity highlights the regulatory role of alternative transcription start sites and their association with noncoding mutations. bioRxiv 176487 (2017). doi:10.1101/176487

175. Ge, H. et al. FusionMap: detecting fusion genes from next-generation sequencing data at base-pair resolution. Bioinformatics 27, 1922–1928 (2011).

176. Nicorici, D. et al. FusionCatcher - a tool for finding somatic fusion genes in paired-end RNA-sequencing data. (2014). doi:10.1101/011650

177. Fonseca, N. A. et al. Comprehensive genome and transcriptome analysis reveals genetic basis for gene fusions in cancer. (2017). doi:10.1101/148684

178. Kahles, A., Ong, C. S., Zhong, Y. & Rätsch, G. SplAdder: identification, quantification and testing of alternative splicing events from RNA-Seq data. Bioinformatics 32, 1840–1847 (2016).

179. Han, L. et al. The Genomic Landscape and Clinical Relevance of A-to-I RNA Editing in Human Cancers. Cancer Cel l 28, 515–528 (2015).

180. Li, Q. et al. Caste-specific RNA editomes in the leaf-cutting ant Acromyrmex echinatior. Nat. Commun. 5, 4943 (2014).

181. Liao, Y., Smyth, G. K. & Shi, W. featureCounts: an efficient general purpose program for assigning sequence reads to genomic features. Bioinformatics 30, 923–930 (2013).

182. Wang, K., Li, M. & Hakonarson, H. ANNOVAR: functional annotation of genetic variants from high-throughput sequencing data. Nucleic Acids Res. 38, e164 (2010).

183. Merico, D., Isserlin, R., Stueker, O., Emili, A. & Bader, G. D. Enrichment map: a network-based method for gene-set enrichment visualization and interpretation. PLoS One 5, e13984 (2010).

184. Yu, G. & He, Q.-Y. ReactomePA: an R/Bioconductor package for reactome pathway analysis and visualization. Mol. Biosyst. 12, 477–479 (2016).

185. Gu, Z., Gu, L., Eils, R., Schlesner, M. & Brors, B. circlize Implements and enhances circular visualization in R. Bioinformatics 30, 2811–2812 (2014).

186. Liberzon, A. et al. Molecular signatures database (MSigDB) 3.0. Bioinformatics 27, 1739– 1740 (2011).

